# An Atlas of Extrachromosomal DNA Structures Illuminates Its Evolution and Biogenesis in Cancer

**DOI:** 10.64898/2025.12.24.696443

**Authors:** Chee Hong Wong, Kuo-Chen Wei, Kuan-Yu Kung, Li-Hao Yang, Yin-Cheng Huang, Chiung-Yin Huang, I-Jiuan Chung, Meng-Fan Huang, Pei-Hua Peng, Meihong Li, Chew Yee Ngan, Kou-Juey Wu, Chia-Lin Wei

## Abstract

Extrachromosomal circular DNA (ecDNA) is a prevalent driver of oncogene amplification across diverse types of cancers. Leveraging single-molecule long-read sequencing and a *de novo* circular genome assembler, ecLego3, we charted a comprehensive atlas of ecDNA structures from longitudinal pairs of primary and recurrent glioblastoma (GBM). Our approach resolved 23 ecDNA genomes representing major oncogenes associated with GBM pathogenesis and revealed a dynamic configuration characterized by pronounced structural variation and multi-species co-existence within individual tumors. Evolutionary trajectories were shaped by hierarchical accrual of structural variants, predominated by deletions and inversions, giving rise to novel oncogene isoforms and enhancer repositioning. Such structural plasticity promotes intratumor heterogeneity and tumor fitness. Multiomic analyses at single cell resolution linked ecDNA abundance and structure to transcriptional output, uncovering non-linear scaling and synergistic co-selection among multiple ecDNA species. Comparative profiling revealed locus-dependent configuration, a contraction of ecDNA diversity upon recurrence, and convergent structural architectures across patients, implicating repeat-mediated genomic rearrangements and replication stress in ecDNA biogenesis. These findings elucidate ecDNA-driven mechanisms of tumor adaptation and highlight new molecular vulnerabilities, informing potential biomarkers and therapeutic strategies for high-grade gliomas.

## Introduction

Extrachromosomal circular DNA (ecDNA), also known as double-minutes (dmin)^1^, has emerged as a pervasive and potent driver of oncogene amplification across diverse cancers^2,3^. Its presence is often associated with aggressive tumor types, like glioblastoma (GBM) and sarcoma^4^, and poor patient outcomes^5^. In aggressive gliomas, ecDNA-mediated focal amplifications are among the most frequent oncogenic alterations^6^, strongly associated with malignant phenotypes, adverse clinical outcomes, and therapeutic resistance^5,7,4^. Ranging from hundreds of kilobases to several megabases in length, ecDNAs are packaged into chromatin complexes comprising oncogenes, enhancers, and immunomodulatory elements^8,9^. Within the nucleus, ecDNAs organize into phase-separated nuclear condensates and ecDNA chromatin hubs function as mobile enhancers, promoting oncogene transcription through long-range DNA interactions, thereby accelerating cellular proliferation and tumor progression^10,11,12^.

EcDNA arises from chromosomal DNA, often through extensive genomic rearrangements, generating multi-segmented molecular architectures derived from one or multiple chromosomes^13^. They are also heterogeneous within a population of cancer cells derived from the same tumor, differing in their complexity and abundance^14,15,16,17^. Such high structural variability creates new oncogenic modules encompassing oncogene variants^7^, splicing isoforms and non-coding regulatory elements^18,10^, which are critical for ecDNA’s tumorigenic function. Multiple, nonexclusive models have been proposed to ecDNA formation, including simple excision^19,20^, chromothripsis^21,22^, translocation-bridge amplification^23^, and breakage-fusion-bridge cycles^21^. Each predicts distinct structural complexity and configuration. Following initial formation, ecDNA molecules continue to evolve in response to microenvironmental cues, tumor progression, and therapy^24,21^, resulting in heterogeneity within tumors where distinct ecDNA species and subtypes could co-exist among tumor cell populations^16,17^. Such structural diversity, together with copy number variation and dynamic enhancer activities, fuels intratumor diversification and ongoing tumor evolution.

Our current understanding of ecDNA architecture largely stems from DNA sequencing, fluorescence in situ hybridization (FISH), and super-resolution microscopy integrated with computational analyses in *in vitro* cancer models^15,13,25^. However, the large size, complex multi-segmented nature, and near identical sequence to chromosomal DNA present substantial barriers to comprehensive characterization of ecDNA molecular configurations via methods like short-read sequencing. Additionally, the presence of multiple highly similar ecDNA species within the same tumor further complicates precise discerning individual ecDNA structures. Consequently, the *in vivo* complexity, diversity, and evolutionary dynamics of ecDNAs remain underexplored despite the mounting evidence of its functional importance in cancer.

The maturation of single-molecule, long-read sequencing (LR-seq) methods has been demonstrated in the analysis of complex rearrangements in cancer genomes^26,27,28^. Capitalizing on recent advances in LR-seq^29,30^ and a specialized *de novo* circular genome assembly pipeline, we systematically reconstruct the complete molecular architectures and evolutionary trajectories of ecDNAs in longitudinal GBM tumors from an extensively characterized patient cohort. Here, we establish an atlas of ecDNA pangenomes encompassing major oncogene amplifications, revealing pronounced intra-tumoral divergence alongside unexpected inter-patient structural convergence. Our data demonstrate that ecDNA structural variation drives the emergence of novel oncogene variants and regulatory landscapes that modulate oncogene transcription and tumor fitness. By integrating ecDNA structural dynamics with measures of abundance and transcriptional output at single-cell resolution, we establish direct mechanistic links between ecDNA structure and functional impact. These high-resolution insights illuminate new models of ecDNA biogenesis and evolution, revealing molecular features exploitable as prognostic biomarkers and therapeutic vulnerabilities in high-grade gliomas.

## Result

### ecLego3: a reference-independent assembler for ecDNA structures

To advance ecDNA structural characterization, we devised ecLego3, a customized circular genome assembler, to comprehensively reconstruct ecDNA molecular structures and dissect their diversity from long-read whole genome sequence (lrWGS) data. ecLego3 begins with k-mer spectrum analysis to empirically determine a sequence coverage threshold reflecting expected diploid abundance, allowing extraction of reads featuring k-mers above this threshold, reflecting high-copy, amplified loci. These reads are then subjected to metaFlye, a *de novo* metagenome assembly algorithm optimized for circular genomes and variable subpopulation abundance^31^, producing draft cyclic graphs that represent the circular genome (CG) candidates of ecDNAs. To elucidate sub-species variation, ecLego3 realigns long-read datasets to assembled cyclic graphs and identifies structural variants using Sniffles2^32^. Variant types and their quantitative abundance are integrated into the final CG representations, defining sub-species structures (see Methods and Figure 1a). We then performed rigorous manual inspection, including error rate analysis of assembled segment alignments to the T2T-CHM13 reference genome to confirm assembly continuity and validity.

**Figure 1.**
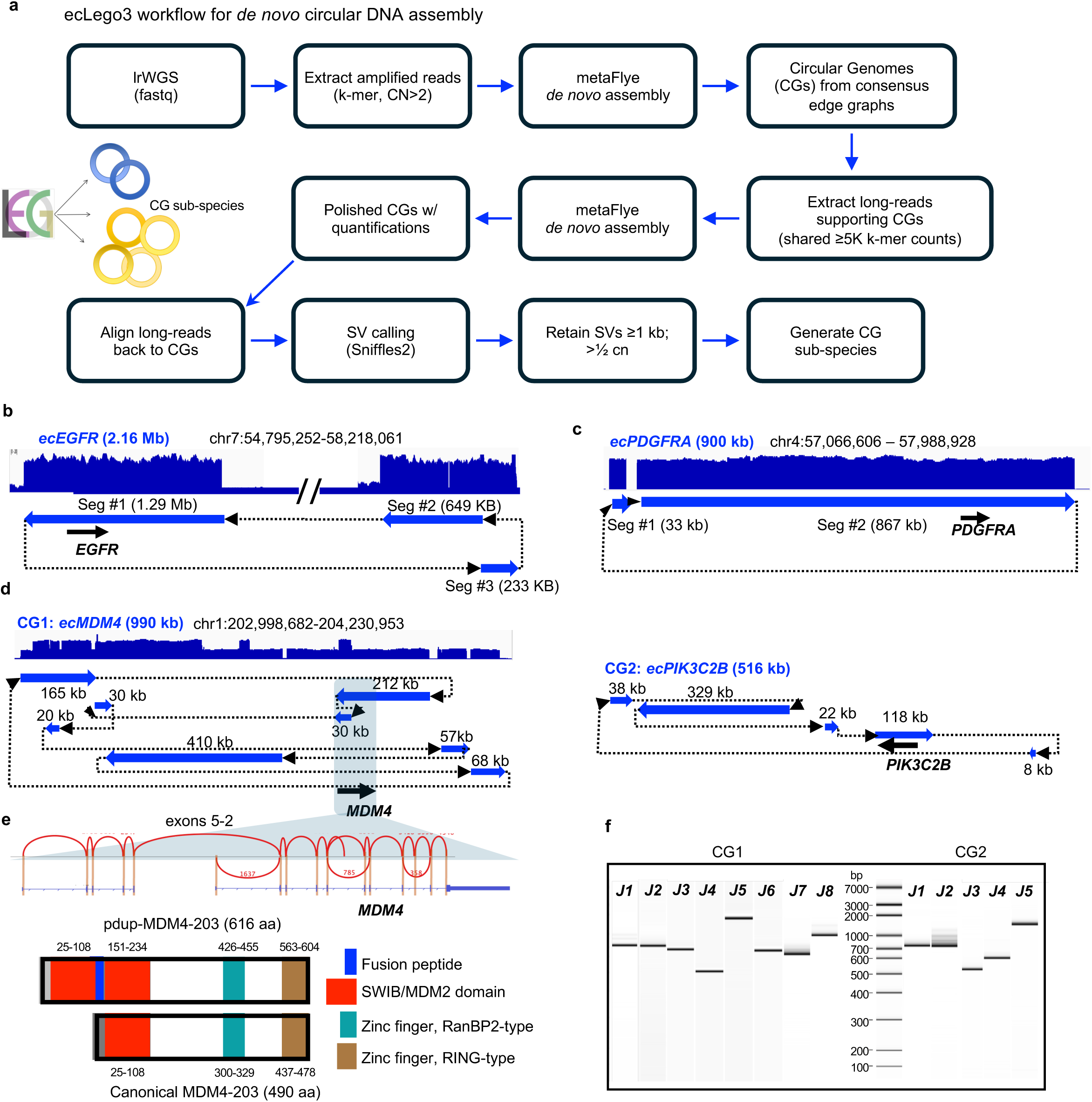
ecLego3 workflow and *de novo* assembly of oncogenic ecDNA structures. **(a)** Schematic of ecLego3 computational pipeline for reference-independent CG reconstruction. The workflow utilizes long-read whole-genome sequencing (lrWGS) to perform k-mer spectrum analysis, *de novo* assembly via metaFlye, and structural variant (SV) detection via Sniffles2. **(b, c)** *De novo* reconstructed architectures of CGs harboring *EGFR* (2.16 Mb) and *PDGFRA* (900 kb) from patient-derived GBM models. The top panel displays genomic locations and long-read alignment coverages of the originated chromosomal regions. **(d)** Structural resolution of co-existing CGs in a single GBM model, identifying CG1 (*ecMDM4*; 990 kb) and CG2 (*ecPIK3C2B*; 516 kb). Arcs indicate internal connectivity; the 30 kb tandem duplication within *MDM4* is highlighted. **(e)** Top: a zoom-in view of the 30 kb duplicated region highlighting *MDM4* chimeric isoform resulted through an in-frame fusion between exons 5 with 2. Bottom: domain organization of canonical MDM4-203 (490 aa) compared to the novel chimeric pdup-MDM4-203 isoform (616 aa), featuring a duplicated SWIB/MDM2 domain. **(f)** Experimental validation of predicted assembly junctions (J1–J8 for CG1; J1–J5 for CG2) via PCR and gel electrophoresis. CN, copy number; aa, amino acids.

The performance of ecLego3 was validated using three patient-derived GBM cellular models harboring extrachromosomal focal amplifications of distinct oncogenes, namely *EGFR*, *PDGFRA*, and *MDM4*, and their ecDNA status confirmed by fluorescence *in situ* hybridization (FISH). High quality lrWGS data (34-46X coverage, 23-28 kb N50) was generated and processed by ecLego3 to reconstruct three distinct CGs: *ecEGFR* (2.16 Mb), ec*PDGFRA* (900 kb) and *ecMDM4* (990 kb) for the respective models (Figure 1b-d). ecLego3 further uncovered a previously unidentified 516 kb CG of *ecPIK3C2B* co-existing with *ecMDM4*. Details on CG sizes, chromosomal origins and estimated copy numbers are provided in Supplementary Table ST.1 & 2. Notably, within *ecMDM4*, the *MDM4* coding region was partially duplicated by a 30 kb segment encoding exons 1-5. This partial gene duplication resulted a novel *MDM4* chimeric isoform (*pdup-MDM4*) through an in-frame fusion between exons 5 with 2 (Figure 1e) whose structure was validated by targeted PCR and Sanger sequencing (Figure 1f, Supplementary Table ST.3). This fusion isoform features duplicated SWIB/MDM2 domains (Figure 1e) and potentially confers enhanced oncogenic activity via greater p53 interactions relative to canonical MDM4. To demonstrate its robustness, ecLego3 was also applied to lrWGS data of lower coverage and read length (Supplementary Figure S1). From lrWGS data (4.3X genome coverage, 18 kb N50) of CHP-212 neuroblastoma cells line previously analyzed using reference-based lrWGS specialized tools CoRAL^33^ and Decoil^34^, ecLego3 successfully assembled a 1.69 Mb *ecMYCN* CG, revealing additional novel junctions and sub-species heterogeneity not detected by Decoil (Supplementary Figures S1 & S2), underscoring the power of reference-independent assembly.

### EcDNA displays locus-dependent configuration and structural heterogeneity

To delineate ecDNA structural configuration *in vivo*, we surveyed grade IV GBM tumors with IDH1/2 wildtype which were known with high likelihood of carrying ecDNAs. From a cohort of 36 patients with distinct (p <0.0001) survival outcome (short-survival, < 12 months, n=19 and long-survival, > 2 years, n=17), 55 tumors were collected, including 21 patients with primary tumors only and 15 patients with longitudinally paired primary and recurrent tumors (named as patient ID_P, R1 and R2) (Figure 2a, b & c). Patients’ metadata, clinical characteristics and treatment history are included in Supplementary table ST.4 & 5. Tumors were subjected to molecular profiling using short-read whole genome sequencing (srWGS) and RNA-seq to evaluate focal amplification and oncogene expression. Consistent with prior genomic studies^35^, widespread copy number alterations (CNAs), including the loss of tumor suppressor *PTEN* and *CDKN2A* gene loci and frequent focal amplification of oncogenes, were observed (Figure 2d). Among 36 primary tumors, 22 acquired oncogene amplifications (CN ≥ 5) and 16 distinct oncogenes were amplified in ≥ 2 tumors. Circular amplification of key oncogenes (*EGFR*, *PDGFRA*, *CDK4*) was detected in 53% of primary and 47% of recurrent tumors by AmpliconArchitect (AA)^13^, with copy numbers ranging from 5 to 116. Tumors harboring ecDNA consistently exhibited higher oncogene expression (Figure 2e), and gene dosage was positively correlated with transcript abundance (Pearson r=0.84−0.93, p<0.001), although nonlinear scaling effects were observed, presumably due to variable configurations of regulatory modules. The copy numbers and gene expressions from selected oncogenes were further confirmed by qPCR (Supplementary Figure S3).

**Figure 2.**
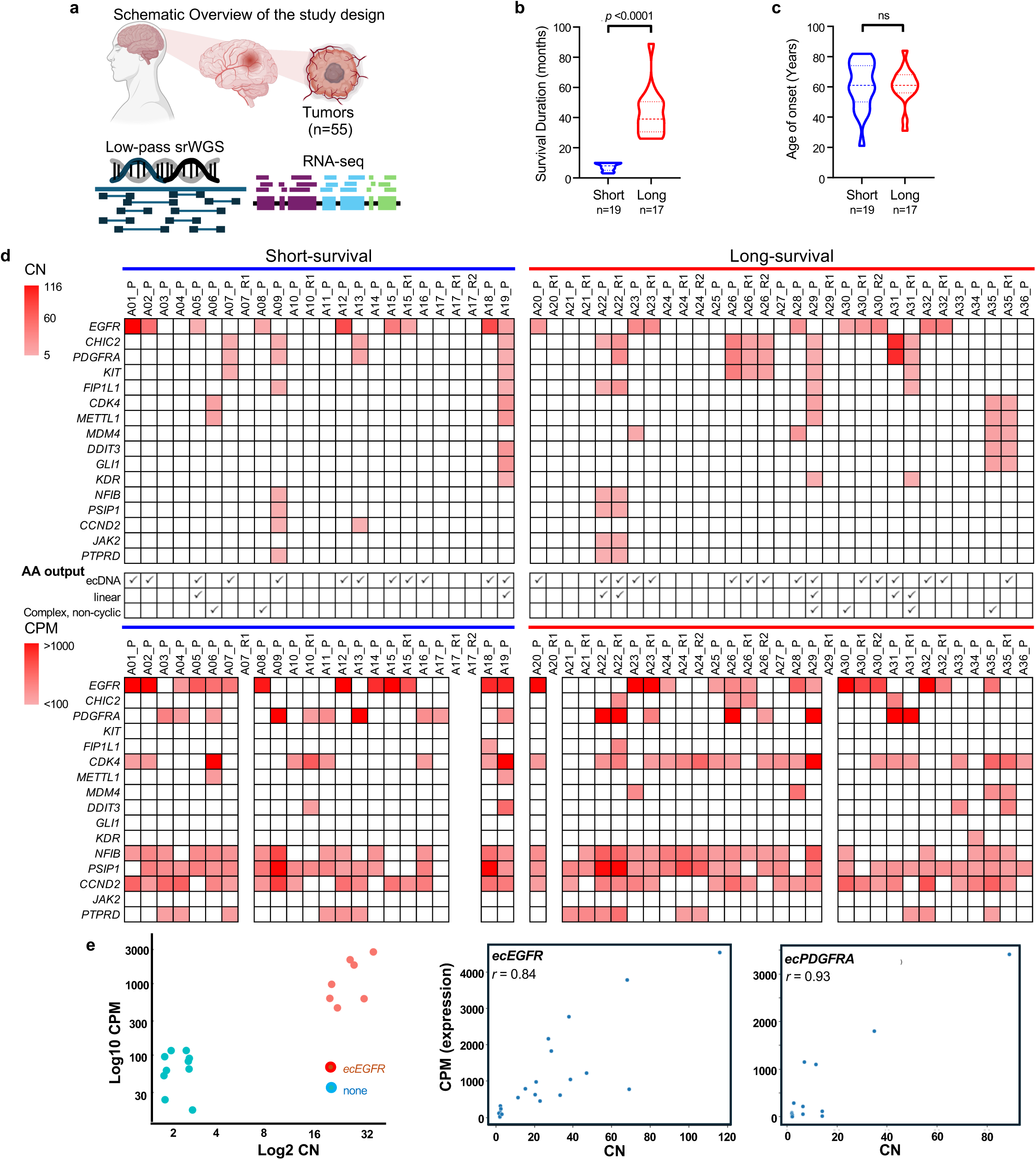
Genomic and transcriptomic surveys of oncogene amplification in GBM patient cohorts. **(a)** Schematic of the study design: 55 IDH-wildtype grade IV GBM tumors were analyzed using low-pass srWGS and RNA-seq. Clinical characteristics of the patient cohort. Violin plots illustrate **(b)** survival duration (months) and **(c)** age at onset (years) between short-survival (*n* = 19) and long-survival (*n* = 17) patients. Center lines represent the median; dashed lines represent the first and third quartiles. *P* values determined by two-sided Wilcoxon rank sum test; ns, non-significant (*P*>0.05). **(d)** Heatmaps showing oncogene copy number (CN, top) and gene expression in counts per million (CPM, bottom). Columns represent individual tumor samples (P, primary; R, recurrent). Row annotations indicate AmpliconArchitect (AA) classifications for focal amplifications, including ecDNA, linear, and complex non-cyclic structures. **(e)** Left: *EGFR* expression in tumors with and without *ecEGFR*. Right: correlation between oncogene CN and transcript abundance (CPM) for *ecEGFR* (*n*=16 tumors) and *ecPDGFRA* (*n*=12 tumors). Pearson’s correlation coefficient (*r*) is shown; all correlations *P*<0.001.

We selected twenty (ten primary and ten recurrent) tumors harboring *EGFR*, *PDGFRA* and *CDK4* circular amplicons and generated high quality lrWGS data (Figure 3a, Supplementary Table ST.1&2). ecLego3 reconstructed 23 ecDNA circular genomes (CGs) encompassing the expected *ecEGFR*, *ecPDGFRA*, *ecCDK4*, as well as *ecMDM4*, *ecMDM2*, *ecKMT2A*, and *ecHMGA2* undetected by AA analysis of the srWGS data. Multi-species ecDNA co-existence was evident in 22% of the tumors, a ratio similar to the rates observed from recent pan-cancer surveys^3,36^. The 3 distinct co-existing CGs of *ecMDM2*, *ecMDM4* and *ecEGFR* reconstructed in A23_P, their chromosomal alignment patterns and CNs were shown in Figure 3b. We aligned the assembled CGs to reference genome for in-depth characterization of ecDNA structures and observed diverse configurations, including simple circles (SCSS: single chromosome, single segment), multi-segmented amplicons from one chromosome (SCMS: single chromosome, multi-segments), and rearranged circles incorporating DNA from multiple chromosomes (MCMS: multiple chromosomes, multi-segment) (Figure 3c). Notably, majority (74%) of the CGs comprised of segments originated from single chromosomes and ∼ half of them were simple circles of one uninterrupted segment. Beyond the unexpected structural simplicity with limited rearrangements, there is a locus-dependent predisposition of ecDNA structural configuration. CGs derived from the *PDGFRA* locus (n=5)—regardless of tumor or patient origin—were consistently assembled as multi-segmented, single-chromosome circles, while half of the CGs derived from the *EGFR* locus displayed as an uninterrupted, single-segment circular structure (Figure 3d & e). The *ecCDK4* amplicons showcased maximal complexity as MCMS. An example of a CG comprising of two major segments (1.3 Mb, 1.14 Mb), bridging *MDM4* and *CDK4* via >500 Mb of intervening sequence originated from three distinct chromosomes is shown in Figure 3f.

**Figure 3.**
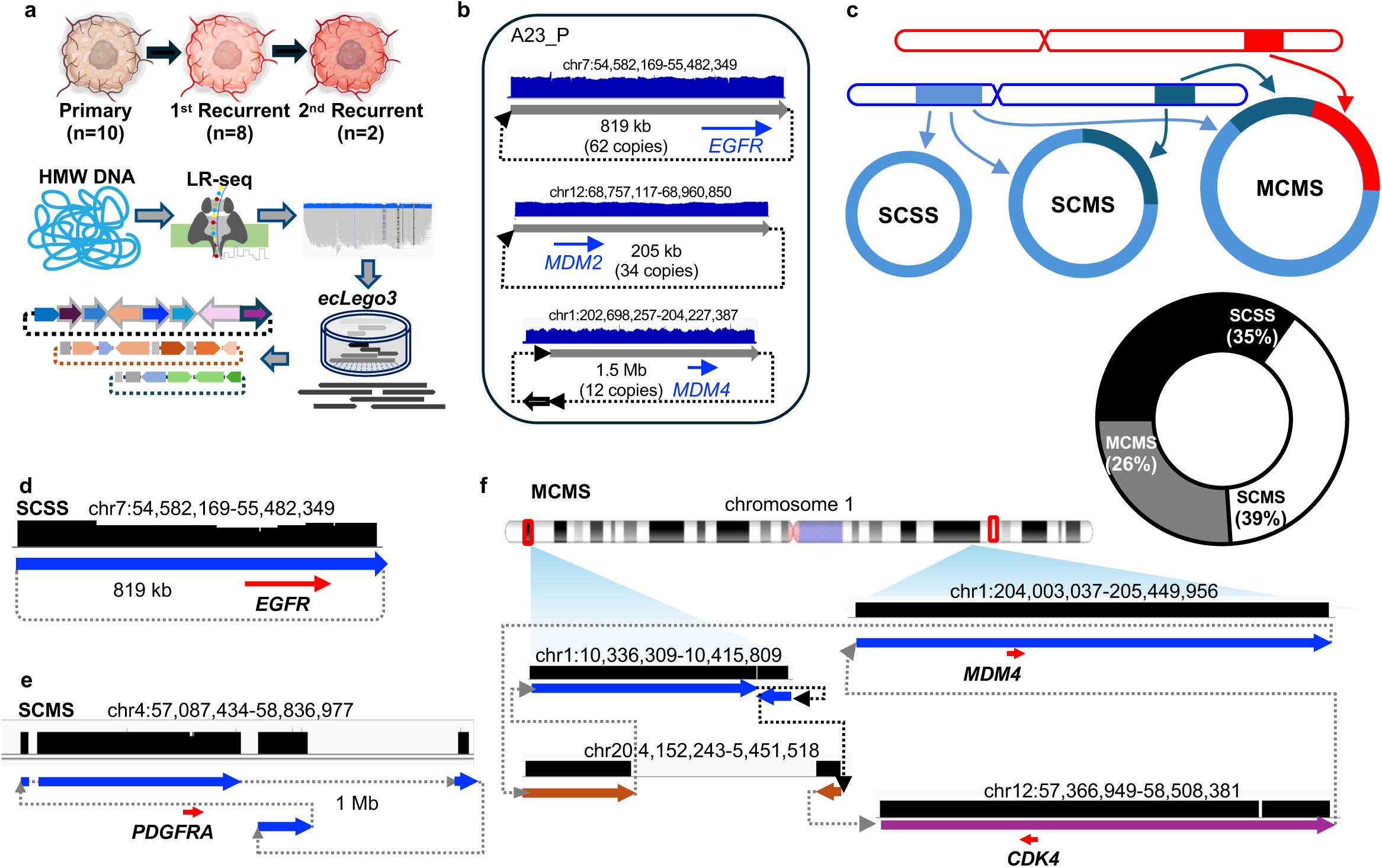
Structural configuration of ecDNA circular genomes. **(a)** high-molecular-weight (HMW) DNA was isolated from primary (*n*=10) and recurrent (*n*=10) GBM tumors for nanopore long-read sequencing (LR-seq), followed by computational reconstruction using ecLego3. **(b)** Co-existence of three distinct CGs (*ecEGFR*, *ecMDM2*, *ecMDM4*) in tumor A23_P, with coordinates (GRCh38), coverage tracks and estimated copy numbers. **(c)** Classification of the assembled CGs into three structural categories: Single Chromosome, Single Segment (SCSS; *n*=8); Single Chromosome, Multi-Segment (SCMS; *n*=9); and Multiple Chromosome, Multi-Segment (MCMS; *n*=6). **(d–f)** Representative ecDNA architectures: **(d)** simple SCSS *ecEGFR* CG in A23_P; **(e)** multi-segmented SCMS *ecPDGFRA* CG in A26_P; and **(f)** complex MCMS *ecCDK4* CG in A35_P, incorporating segments from chromosomes 1, 12, and 20. Colored blocks represent genomic segments; blue/brown/purple arrows indicate orientation; red arrows denote oncogene and transcription direction.

Analysis of intra-tumoral ecDNA heterogeneity by subpopulation structure revealed considerable variation. Most CGs existed as multiple subspecies with up to 11 sub-species observed, each characterized by distinct structural variants (SVs). Subspecies structures were corroborated by junction PCR, Sanger sequencing, and supporting short-read data from their corresponding tumors (Supplementary Table ST.2 & 3). EcDNA heterogeneity (CGs with multiple sub-species) was significantly higher in primary tumors (69% vs 40% in recurrences; Wilcoxon p=0.008), and tumors with highest subspecies diversity were exclusively primary (Supplementary Figure S4a). The number of subspecies displayed a moderate positive correlation with overall copy number (Spearman ρ=0.58, p=0.011), implicating ecDNA copy number expansion as a substrate for emergence of new variants. Such diverse configurations of ecDNAs, co-existence of multi-species ecDNAs and the prevalence of their subspecies underscore ecDNAs’ structural dynamics in cancer progression.

### EcDNA subspecies drive new oncogene variants that enhance tumor fitness

The analysis of subspecies variation and their abundance in individual tumors provides unique insights into the evolutionary trajectory and selective pressures shaping ecDNA adaptation. Among 67 ecDNA subspecies characterized, deletions (n=22) and inversions (n=14) emerged as the most dominant SV categories defining the diversity of ecDNA subpopulations. These subspecies are established by distinct and combinatorial SVs, illustrated by the *ecEGFR* circles in A12_P (cn=57) and A15_P (cn=61), which displayed as SCSS configurations of 620 kb and 1.33 Mb circles, respectively, dividing into 8 and 11 subspecies (Figure 4a, Supplementary Figure S4b). *EcEGFR* harboring 10 kb and 16 kb deletions constituted the most abundant subpopulations in A12_P and A15_P, accounting for 46% and 56% of total *ecEGFR* populations, respectively. These variants eliminated *EGFR* exons 2-4 and 2-7 and generated constitutively active, ligand-independent EGFR isoforms which are linked to therapeutic resistance to tyrosine kinase inhibitors in GBM^35,7^. Comparative analysis of acquired deletions among the subspecies enabled us to reconstruct their molecular phylogenies and reveal parsimonious evolutionary paths where a representative circular genome (P1) independently acquired a set of SVs, followed by hierarchical introduction of further variants through stepwise double-strand breaks (DSBs) and rejoining at distinct breakpoints (Figure 4b, Supplementary Figure S4c).

**Figure 4.**
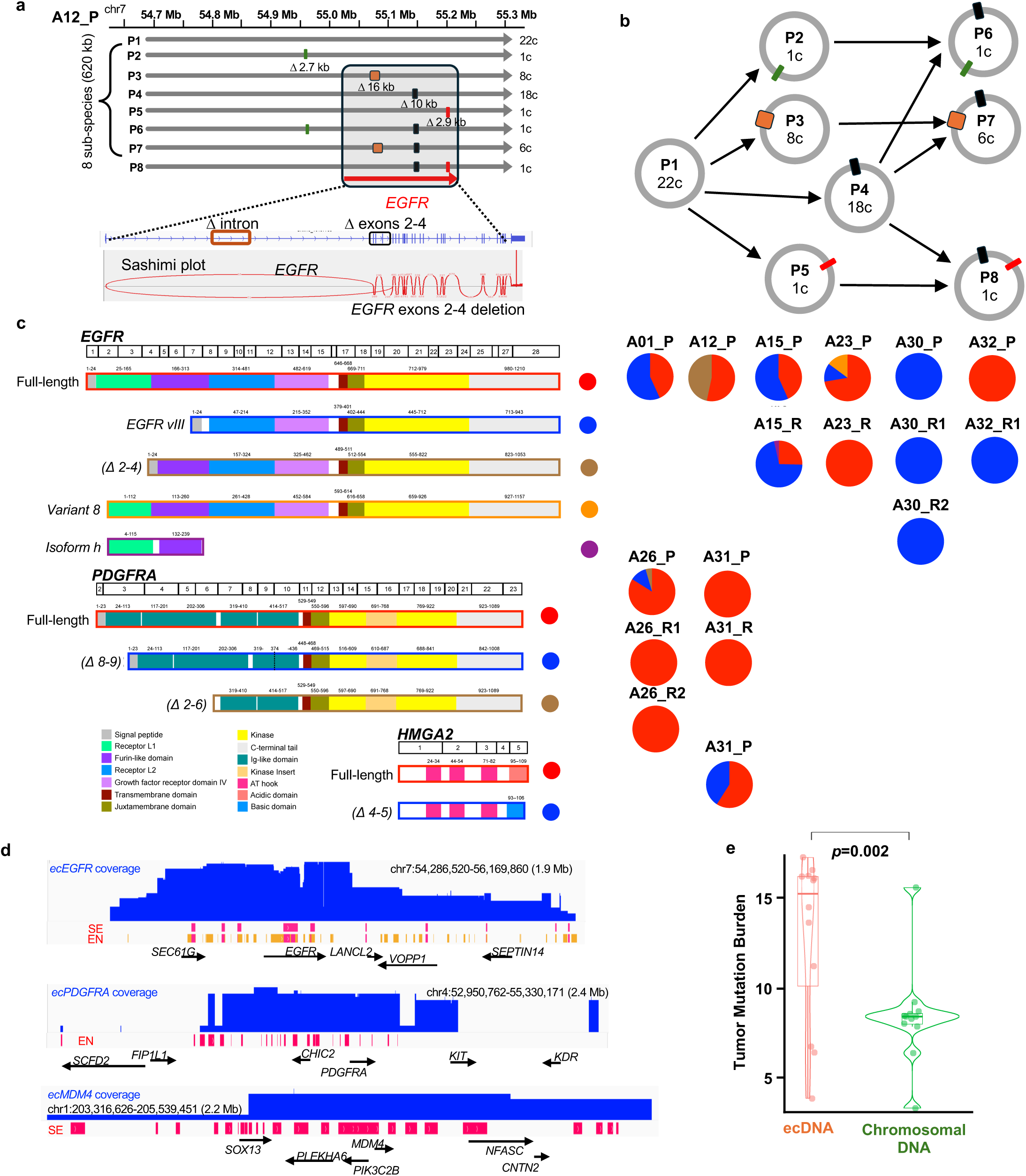
Structural evolution of ecDNA generates diverse oncogenic isoforms and enhances tumor fitness. **(a)** Architecture of *ecEGFR* subspecies (P1–P8) in A12_P. Internal deletions (colored boxes, sizes noted in kilobases (kb)) and sashimi-confirmed splice junctions define subspecies diversity. **(b)** Molecular phylogeny showing stepwise structural variant acquisition and copy numbers (*c*) from ancestor P1. **(c)** Truncated oncogene isoforms (*EGFR*, *PDGFRA*, *HMGA2*) driven by ecDNA variants. Domains are color-coded as indicated in the legend. Pie charts (right) quantify cohort-wide isoform abundance and co-occurrence. **(d)** Genomic coverage of *ecEGFR*, *ecPDGFRA*, and *ecMDM4* CGs, highlighting co-amplification with super-enhancers (SE) and enhancers (EN). **(e)** Higher tumor mutation burden (TMB) in ecDNA regions versus their corresponding chromosomal regions (*n*=12 tumors, *p*=0.002, two-sided Wilcoxon signed-rank test). The boxplot indicates the median and interquartile range (IQR); whiskers extend to 1.5 x IQR.

To assess how these subspecies shaping ecDNA adaptation, we evaluated their impacts on oncogene transcription. These structural variants modulate oncogene transcription by altering coding sequences or reorganizing regulatory modules. In 11 tumors, both known and novel *EGFR*, *PDGFRA*, and *HMGA2* coding variants were detected from distinct ecDNA subspecies (Figure 4c), each supported by RNA-seq junction reads. Variants included truncated *HMGA2* (Δ exons 4-5), *EGFR* (Δ exons 2-4, Δ exons 2-7, a.k.a. *EGFRvIII*), variant 8 (NM_001346900.2), isoform h and *PDGFRA* (Δ exons 2-6, Δ exons 8-9). Together with the *ecMDM4* encoded duplicated SWIB/MDM2 domains found in the patient-derived cell model (Figure 1e), all these ecDNA CG variants encoded in-frame fusion proteins with either intracellular or extracellular domains partially removed, while *EGFR* isoform h featuring only the ligand-binding domain^37^. These engineered oncogene variants confer positive selection, surpassing canonical isoforms in driving tumor fitness and resistance. In addition to *EGFRvIII’s* well-established drug resistance phenotype^7^, truncated *HMGA2* (Δ 4-5) is known for its enhanced DNA-binding affinity^38^, and *PDGFRA* Δ exons 8-9, present in 40% of the *PDGFRA* amplified cases, exhibits elevated tyrosine kinase activity and transforming capacity^39^, supporting the notion that ecDNA diversity nurtures oncogenic potency and plasticity. Thus, ecDNA structural plasticity is a key contributor to intratumor heterogeneity through the ongoing creation of novel, potentially gain-of-function oncogene variants. Enhancers on ecDNAs are central to oncogenic transcriptional activation and ecDNA’s tumorigenic activities^10,9^. In the assembled CGs, enhancer (EN) and super-enhancer (SE) loci are densely distributed across ecDNA circular genomes (Figure 4d). Despite the prevalent deletions among the diverse subspecies, SE-containing segments were largely conserved across primary and recurrent tumors. Deletions that remove SE-containing segments were only observed in minor ecDNA subpopulations in two tumors (23% in A15_P, 13% in A23_P, Supplementary Figure S4b & d), and were absent in recurrent tumors, emphasizing their functional importance.

SVs within ecDNA subspecies coexists with increased single nucleotide variant (SNV) burden. EcDNA CGs harbor significantly higher tumor mutation burden (TMB) (Wilcoxon signed-rank test, p=0.002) than corresponding chromosomal regions (Figure 4e), enriched with known GBM SBS1 and SBS5 (single base substitution) signatures^40^. Of 45 *ecEGFR* subspecies, a substantial fraction (n=23) carried the non-synonymous *EGFR* variant K521R (G→A, exon 13), a variant associated with increased glioma risk^41^, highlighting the confluence of structural and sequence-based evolution on ecDNA.

Collectively, the widespread ecDNA instability, in the forms of SVs and single base substitution, is integral to the generation of intratumor heterogeneity. Deletions and inversions are the major sources of subspecies formation, driving the emergence of new oncogene isoforms while the regulatory enhancers are largely insulated. Once formed, the circular genomes incorporate little new DNA, as evident by limited insertions. The stepwise, hierarchical accrual of SVs appears to balance oncogenic potency with minimization of “passenger” DNA, yielding ecDNA configurations that sustain selective advantages during tumor evolution.

### EcDNA structural dynamics shape regulatory plasticity and adaptive selection in recurrent tumors

Comparative profiling of ecDNAs and their subspecies between matched primary and recurrent tumors revealed that primary tumors harbor greater ecDNA diversity, with more subspecies and frequent co-existence of multi-species ecDNAs carrying distinct oncogenes. When tumors recurred, ecDNA displayed a contraction in diversity, characterized by reduced subspecies and increased uniformity among ecDNA species (Figure 5a). This shift suggests purifying selection acting upon ecDNA structures during cancer progression.

**Figure 5:**
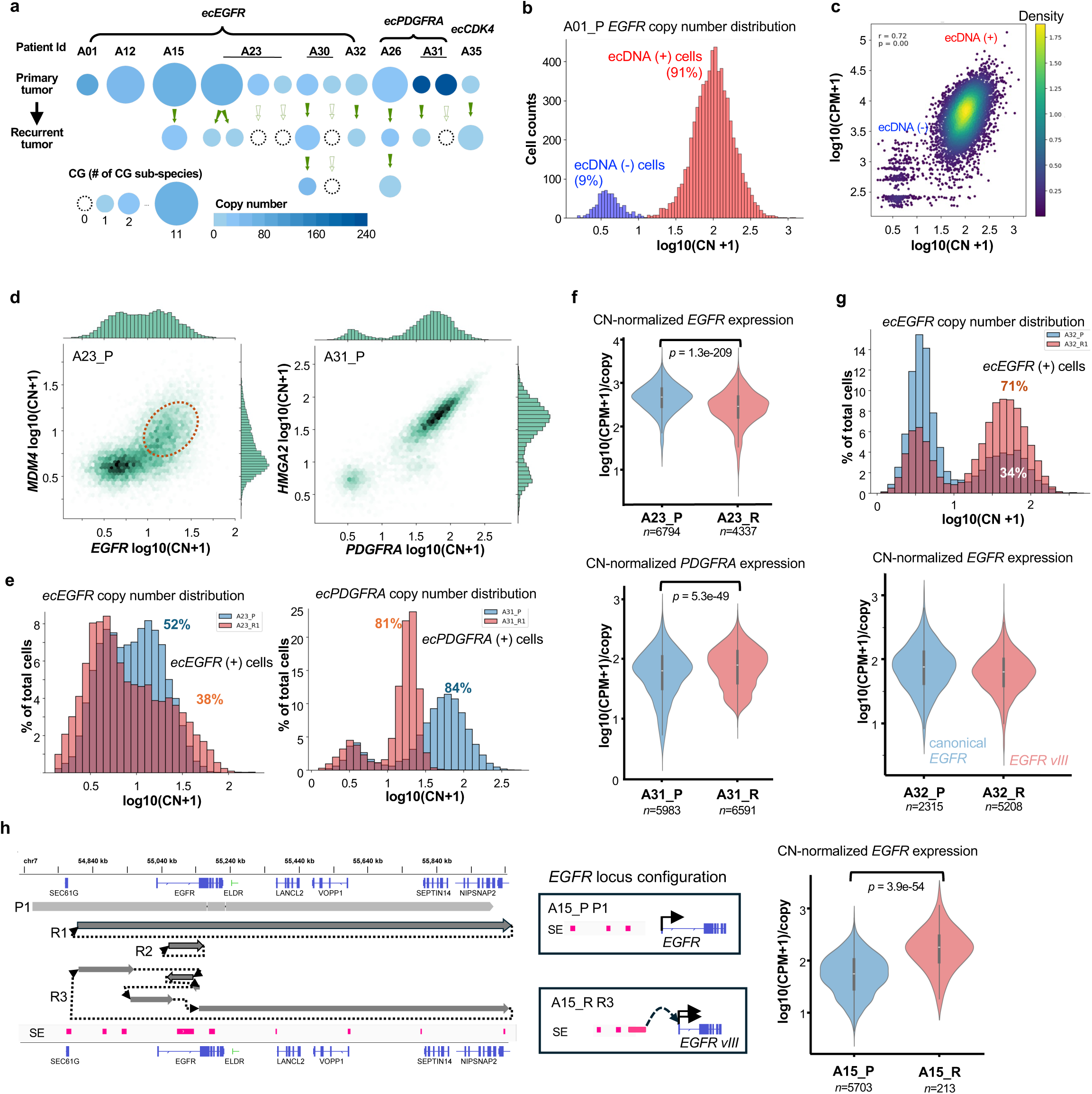
ecDNA structural dynamics drive regulatory plasticity and adaptive selection in recurrent tumors. **(a)** Longitudinal analysis of ecDNA species (*ecEGFR, ecPDGFRA, ecCDK4*) across matched primary (P) and recurrent (R) tumors (*n*=9 patients). Circle size denotes copy number (CN) inferred from long-read WGS and *ecLego3* characterization; vertical stacks and color-coded arrows illustrate the evolution and contraction of circular genome (CG) sub-species diversity in recurrence. **(b, c)** Distribution of *ecEGFR* CN per cell in A01_P tumor, showing a normal distribution of ecDNA(+) cells (red) and a positive correlation (Pearson *r*=0.72) between ecDNA dosage and transcriptional output. **(d)** 2D density plots of *ecEGFR/ecMDM4* and *ecPDGFRA/ecHMGA2* CNs within single nuclei in A23_P and A31_P, demonstrating the co-segregation of multi-species ecDNAs. **(e)** Longitudinal shifts in the CN distributions and fractions of ecDNA(+) cell populations. **(f)** Transcriptional output (CN-normalized expression) of *ecEGFR* and *ecPDGFRA* in primary (blue) and recurrent (red) states. **(g)** Clonal expansion of *ecEGFR*(+) cells in A32_R tumor, where the *ecEGFR*(+) fraction increases from 34% to 71%, despite the comparable overall *EGFR* expression per cell. **(h)** Enhancer hijacking upon tumor recurrence. Structural rearrangement of the *EGFR* locus in the R3 subspecies brings distal super-enhancers (SE, pink) into proximity with the *EGFR vIII* variant, significantly boosting its transcriptional efficiency (*p*=3. 9 ×10^-54^). Center lines in violin plots represent medians; *n* indicates nuclei counts. Statistical significance determined by Wilcoxon rank-sum tests.

To interrogate the functional impacts of structural variation on adaptation, single-nucleus multiomic (scRNA + scATAC) analyses were performed to measure longitudinal trends in ecDNA abundance and oncogene transcription at single-cell resolution. EcDNA amplification and copy numbers were inferred by normalized scATAC-seq read coverage as previous described^36^ at ecDNA loci, with non-amplified regions as controls and a Gaussian Mixture Model establishing thresholds for ecDNA positivity and relative copy number per nucleus. Notably, *ecEGFR* (+) cells comprised 91% of total cells (n=7,298) in a representative A01_P tumor, with a median CN of 100 (Figure 5b), which closely matched to the CG copy numbers inferred from lrWGS data (median CN 88), supporting analytic accuracy. As expected by the random segregation model during cell division^42^, ecDNA CNs displayed a normal distribution with marked cell-to-cell variability (log10(CN+1), mean 2, σ 0.26; Supplementary table ST. 6), strongly correlated with oncogene transcription (Pearson PCC=0.7, p=0, Figure 5c). Such varability enables ecDNA-driven tumors more adaptable in different tumor microenvironments.

CG distributions across individual cells further elucidated co-dependency among multi-species ecDNAs of distinct oncogene amplicons. In A23_P, ecLego3 independently reconstructed *ecMDM4* (1.5 Mb, cn=12) and *ecEGFR* (819 kb, cn=62). We revealed a 1.75-fold enrichment of cells containing both CG species (chi-square p=0). Similar significant enrichment (1.7-fold) of co-existence was observed for *ecPDGFRA* (699 kb, cn=228) and *ecHMGA2* (603 kb, cn=245) in A31_P (chi-square p=0). Moreover, their copy numbers also exhibited significant correlations (p-value = 0 for *ecPDGFRA-ecHMGA2* and 6.2e-144 for *ecEGFR-ecMDM4*) (Figure 5d), supporting the co-segregation model. Such co-selection likely contributes to higher fraction of ecDNA (+) cells and significantly elevated *EGFR* expression (Wilcoxon p=1.3e-209) in primary, compared to relapsed tumors with undetected *ecMDM4* (Figure 5e & f), suggesting regulatory synergy and fitness advantages conferred by multi-species ecDNA clones. Functionally, TP53 pathway disruption via *MDM2/4* amplification can escalate genome instability and foster ecDNA emergence, while HMGA2, a known transcriptional regulator, could modulate *PDGFRA* signaling, further potentiating oncogenic progression.

Recurrent tumors can acquire new ecDNA structural variants that further enhance tumor fitness. This is illustrated in A32_P and A32_R tumor pairs where a dominant *ecEGFR* subspecies with an 18 kb deletion spanning *EGFR* exons 2-7 (*EGFRvIII*) became prevalent upon relapse (Figure 4c), raising *ecEGFR*(+) cell fractions from 34% in the primary (canonical *EGFR* expression) to 71% in the recurrence, despite comparable overall *EGFR* expression within individual cells (Figure 5g). In rare cases, massive structural reorganization induces emergence of entirely new circular genomes (CGs), diversifying the regulatory modules of ecDNAs in relapsed tumors. In A15_R, a new *ecEGFR* CG was detected in three distinct subspecies, an uninterrupted 1.1-Mb circle (R1), an 82 kb circle (R2) carrying a well-characterized superenhancer (ecSE) critical for *EGFR*-mediated malignancy^43^ in the absence of any *EGFR* coding region, and a 1.2 Mb circle (R3) with rearranged regulatory modules, wherein the ecSE was relocated proximally to the 5’ of the *EGFR* promoter (Figure 5h). This rearrangement coincided with significantly upregulated *EGFR* expression in recurrence (Wilcoxon signed-rank test, *p*=3.9e-54), suggesting *cis* activation effects by ecSE repositioning.

In aggregate, ecDNA copy number variation, structural remodeling and co-selection contribute regulatory module plasticity and oncogene variants, conferring adaptive advantages. EcDNA abundance exhibits pronounced cell-to-cell variation, and oncogene expression is not strictly copy number-dependent, highlighting the impacts of regulatory architecture and multi-species co-selection in tumor evolution.

### Structural convergence reveals locus-dependent, repeat-driven ecDNA biogenesis

While individual CGs harboring prevalent variation as distinct subspecies, they also exhibit striking convergence in molecular architecture across diverse tumors from different patients, regardless of pathological manifestation or survival outcome. Common segments, breakpoints and rearranged junctions were shared among circular genomes carrying either *EGFR* or *PDGFRA* between tumors from distinct patients (Figure 6a). Notably, nine breakpoints originating from chromosome 7 (*EGFR* locus) and chromosome 4 (*PDGFRA* locus), and five junctions involving shared DNA segment order, were detected in CGs from at least two different patients (Supplementary Table ST.7). Comparative analysis with Glioma Longitudinal Analysis (GLASS) and Pan-Cancer Analysis of Whole Genomes (PCAWG) cohorts underscores the non-random nature of the observed ecDNA structural convergence. We identified significant breakpoint conservation, sharing 88 breakpoints with GLASS (12.9%; 2.7-fold enrichment; P<0.0005) and 199 with PCAWG (22.8%; n=155; 3.2-fold enrichment; P<0.0001). Conversely, dual-end criteria yielded zero overlapping segments or junctions. Thus, while genomic breakage hotspots are conserved; subsequent recombination generate unique, patient-specific ecDNA topologies.

**Figure 6.**
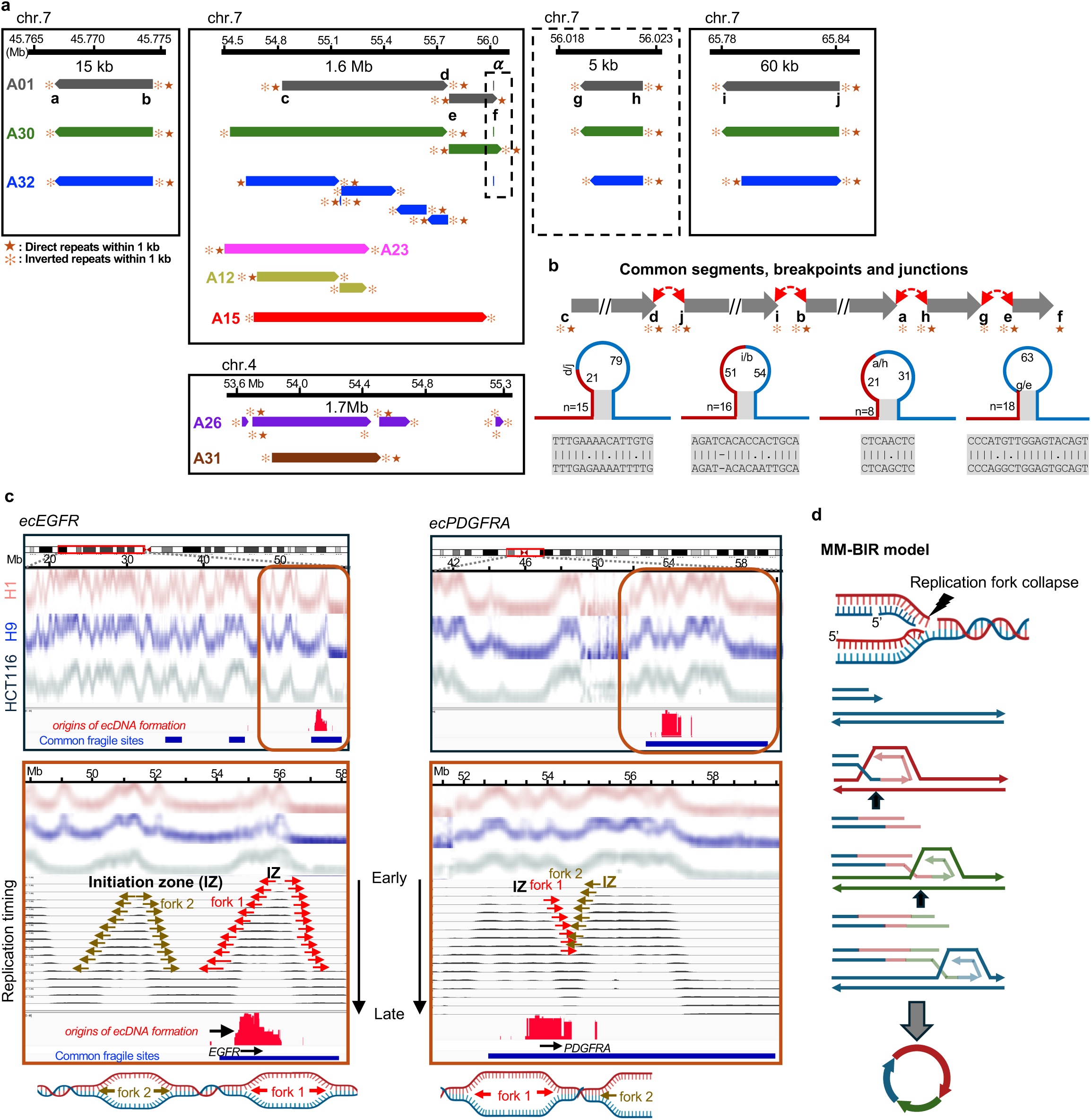
Structural convergence and MMBIR-mediated biogenesis of ecDNA. **(a)** Genomic architecture of recurrent ecDNA breakpoints on chromosomes 7 (*EGFR*) and 4 (*PDGFRA*). Tracks for patient-derived tumors (e.g., A01, A15) highlight conserved segments and structural hotspots flanked by direct (gold stars) and inverted (orange asterisks) repeats. The α label denotes a region of structural convergence within the 1.6 Mb overview (dashed box), which is resolved in the adjacent high-resolution inset to distinguish clustered focal breakpoints (vertical dashed lines) from flanking segments (*e*,*f*). **(b)** Characterization of shared junctions. Schematics depict the potential stem-loop structures formed within the junctions (e.g., d/j), where values (e.g., 21, 79) denote loop lengths (bp) between the junction breakpoint and the base of the flanking inverted repeat stem. Alignments reveal 8–18 bp microhomologies (*n*), a defining hallmark of microhomology-mediated break-induced replication (MMBIR). **(c)** Spatiotemporal correlation between replication timing and ecDNA formation. Repli-seq profiles in hESCs (H1, H9) and HCT116 cells demonstrate that ecDNA origins (red) and common fragile sites (CFS, blue) coincide with late S-phase initiation zones (IZ) and regions of replication fork convergence. **(d)** Proposed mechanistic model. Replication stress-induced fork collapse at repeat-rich loci triggers microhomology-mediated strand invasion and template switching, facilitating the circularization and complex structural rearrangements characteristic of ecDNA.

The structural convergence suggests genomic features or mechanisms pre-disposing specific regions for ecDNA formation. To parse the underlying genomic determinants, sequence analysis of common breakpoints and junctions revealed they were frequently flanked by clusters of annotated direct and inverted repeats (Figure 6a & b). Extending the analysis to a broader set of breakpoints (n=50) from all CGs, we detected a significant enrichment for inverted repeats (1.5-fold) and Z-DNA motifs (2.8-fold, p < 0.05), which are the genomic features known for enhancing structural DNA instability^44,45,46^. Manual inspection further showed that these junctions often involved microhomologies of 8–18 bp, a hallmark of microhomology-mediated break-induced replication (MM-BIR), previously implicated in the creation of complex DNA copy number variants^47^. Transcription-replication conflicts (TRCs) are known to cause DNA damage and provoke replication fork collapse at late S-phase^48,49^. MM-BIR initiates DNA synthesis at short regions of homology flanking breakpoints, yielding complex rearrangements and copy number variants. Therefore, we hypothesized that active transcribed loci flanked by repeat sequences and replicated late in S-phase could be susceptible to MM-BIR in replication stall to drive ecDNA formation. Analysis of high-resolution replication timing maps created through Repli-seq approach^50^ revealed that *ecEGFR* and *ecPDGFRA* originate in regions bordering late-replicating forks (Figure 6d). These locations overlapped with common fragile sites (CFSs), known as genomic loci hypersensitive to replication stress and known hotspots for chromosomal rearrangements^51^. Crucially, this specific timing signature was corroborated by independent datasets from the GLASS and PCAWG consortia, where breakpoints driving *ecEGFR* and *ecPDGFRA* formation were significantly enriched within the late-mid S-phase (S7–S10) relative to background models restricted to these amplicon loci (*P* < 0.001 for *ecEGFR* across all cohorts, Supplementary Table ST.8). The robust persistence of this temporal profile within genomic contexts deriving extrachromosomal amplicons underscores the universality of these replication-timing boundaries as potential drivers of ecDNA formation.

Based on such pattern, we propose TRCs induced MM-BIR as a mechanism of ecDNA biogenesis. By annealing single-strand 3’ tails from broken replication forks, MM-BIR primes DNA synthesis through repeats on nearby single-stranded regions, creates R-loops and generates complex rearrangements with singular or multiple template switches, leading to eventual re-establishment of processive replication and ecDNA circles (Figure 6d). This mechanism is supported by the identification of intervening DNA sequences embedded within segment junctions by the reference-free algorithmic design of ecLego3. These novel insertions ranged from 100 bp to 1.6 kb, many of which have poor or no alignment to the human reference genome and displayed high enrichment for repetitive DNA elements (65–98%), predominantly derived from mobile element insertions (MEIs) such as LINE/L1 and SINE/Alu subfamilies (Supplementary Table ST.9). PCR with Sanger sequencing reaffirmed the sequences of these insertions (Supplementary Information), further validating ecLego3’s ability to resolve convoluted regions inaccessible by reference-guided approaches.

## Discussion

GBM, a highly aggressive brain tumor in which ecDNA is both prevalent and functional important. By generating a high-resolution atlas of ecDNA molecular structures across matched primary and recurrent patient tumors, we revealed the structural diversity, subspecies variation and longitudinal dynamics of ecDNAs and associated their effects on tumor fitness and adaptation. Contrary to the expectations based on the chromothripsis model, which predicts highly complex, fragmented rearrangements, ecDNAs in GBM tumors display unexpected structural simplicity and common genomic features. The shared structural signatures are potential hallmarks of GBM-specific traits that may serve as valuable biomarkers for prognosis or therapeutic targeting.

The application of LR-seq enabled accurate, reference-independent reconstruction of ecDNA structures. Computational tools like AmpliconArchitect (AA) and AmpliconReconstructor (AR)^13,16^ efficiently map circular amplicons, but they are limited in studying their variation and subpopulations. While long-read specialized tools like CoRAL^33^ and Decoil^34^ can inform ecDNA structural variation, their reliance on reference genomes restricts the discovery of non-templated synthesized sequences flanking the segments within ecDNA circular genomes. The ecLego3 pipeline, which leverages k-mer based read extraction and metagenome assembly, facilitates *de novo* ecDNA assembly, allowing unique insights into the diversity of ecDNA architectures which are critical for understanding tumor heterogeneity.

EcDNA originates from chromosomes. Multiple, nonexclusive models have been proposed to explain ecDNA biogenesis, including excision, chromothripsis, and breakage-fusion-bridge cycles^19,20,21,22,23^. Here, integrative structural and sequence analysis supports a locus-dependent model where ecDNA arises through the interplay between replication stress associated DNA fragility, and repeat-mediated alternative end-joining repair pathways (alt-EJ)^47^. In this context, MM-BIR is triggered by TRCs, primes DNA synthesis through single-stranded repeat pairing and, with multiple template switches, leads to processive replication of circular DNA. This proposed mechanism is supported by findings in other organisms like yeast and drosophila, where activated transcription and replication stress prime the formation of circular (ecc)DNAs^52,53^. Furthermore, prior studies have shown that elevated TRCs and single-strand DNA are recognized features in ecDNA (+) tumors^54^. The recruitment of alternative DNA repair pathways, shaped by DNA lesion types, cell-cycle stages, and chromatin states, suggests ecDNA genesis is context-specific, with diverse therapeutic implications.

Once established, ecDNA molecules are highly dynamic, giving rise to diverse subspecies that encode new oncogene variants and reconfigure regulatory modules. The combination of large intercellular variation of ecDNA copy number and oncogene expression amplifies intratumor heterogeneity. The co-existence of multiple ecDNA species, each bearing distinct and potentially synergistic oncogenes, even with random segregation during cell division^55,56^ highlights the adaptive advantage conferred by ecDNA co-selection. Notably, previous genomic studies revealed that the evolutionary trajectories of ecDNA-marked and chromosomally marked tumor subclones diverge^57^. Clonal selection of ecDNA-harboring cells enables rapid cellular adaptation and enhances tumor aggressiveness and therapy resistance. Therefore, systematic characterization of ecDNA structural variation provides a new dimension for understanding tumor evolution, heterogeneity, and fitness in tumor progression. Ultimately, expanding ecDNA structural studies to larger tumor cohorts and additional cancer types is expected to transform our understanding of oncogene regulation and inform new avenues for early detection, and precision therapies targeting these DNA circles.

## RESOURCE AVAILABILITY

### Lead contact

Further information and requests for resources and reagents should be directed to and will be fulfilled by the Lead Contact, Chia-Lin Wei.

### Materials availability

All unique/stable reagents generated in this study are available from the lead contact with a completed materials transfer agreement. Information and inquiry for sequencing data and patient tumors should be directed to co-corresponding author, Kou-Juey Wu.

### Data and code availability

Data deposition is in progress and the (reviewer) accession id will be available when ready. ecLego3 is available at https://github.com/cheehongsg/ecLego.

## Acknowledgments

We thank the Genome Sequencing Core at Chang Gung Memorial Hospital, Linkou, and the Genome Technologies Service at The Jackson Laboratory for their expert technical assistance. We are grateful to Albert H. Kim and the Tumor Bank of the Brain Tumor Center, St. Louis, for providing the patient-derived glioblastoma cell lines (B168, B171, and B186), and to Yi-Hsin Tsai for technical support. This work was supported by Chang Gung Memorial Hospital grants OMRPG3I0011, OMRPG3I0012, and OMRPG3I0013 (to K.J.W.) and CORPG3N0271 (to K.C.W.).

## Author contributions

C.H.W., C.L.W., K.C.W. and K.J.W. conceptualized the project. C.H.W., C.L.W., C.Y.N., K.J.W., K.Y.K., M.F.H., M.H.L., P.H.P. and Y.C.H. designed the experiments. C.Y.H., C.Y.N., I.J.C., M.F.H., M.H.L. and P.H.P. performed wet lab experiments. C.H.W., I.J.C., K.Y.K., L.H.Y. and M.F.H. conducted bioinformatic analyses. C.H.W., C.L.W., K.Y.K. and K.J.W. wrote the manuscript with input from all authors. C.L.W., K.C.W. and K.J.W. supervised the project.

## METHODS

### EcLego3 *de no*vo assembly of ecDNA circular genomes

The ecLego3 pipeline (https://github.com/cheehongsg/ecLego) performs the k-mer based enrichment of ecDNA reads by identifying k-mers (k=29) present in the long-read data. A coverage threshold is established for the k-mer spectrum, representing the expected abundance for sequences within the diploid nuclear genome. All k-mer analysis are performed with KMC3^58^. Next, these enriched long reads are assembled using metaFlye^31^. MetaFlye utilizes a de Bruijn graph (DBG) approach^59^ to assemble contigs (disjointigs) and construct a repeat graph. Read mapping is employed to further resolve this repeat graph. ecLego3 traverses the assembled graph and eliminates edges likely belonging to the linear chromosomes (i.e., edges connecting to terminal vertices). The remaining graph is expected to be cyclic, representing the potential circular structure of ecDNA molecules. Comprehensive set of long reads supporting this cyclic graph are identified using the k-mer approach. These reads are subjected to *de novo* assembly, potentially leading to longer contigs and simplified assembly interpretation. The long reads are subsequently mapped back to the re-assemblies, and structural variations (SVs) are identified using Sniffles2. Finally, the identified SVs are integrated into the re-assemblies to generate assemblies representing the various ecDNA sub-populations within the sample.

### GSC Neurosphere Culture

Cells were plated using DMEM/Ham’s F12 (Gibco) with Glutamax and B27 without vitamin A supplement (Life Technology) for neurosphere culture. Murine EGF and Human FGF-2 (Peprotech INC), each at 20 ng/ml, were added to culture media for neurosphere cultures. Neurosphere GSCs were grown on ultra-low attachment dishes. Media were replaced with fresh GSC media every 2–3 days. If sphere become large, then use Accutase for cells dissociation. Cells were routinely used between passages

### GBM Tumor Collection

The frozen tumors from 36 patients with primary and recurrent glioblastoma (GBM, grade Ⅳ) were collected with written informed consent from patients, using a protocol approved by the Institutional Review Board (IRB) of The Chang Gung Medical Foundation Institutional Review Board (104-2656B). The study cohort including 22 male and 14 female patients with a median age at diagnosis of 61 years (range 21–84 years) (Supplementary Table ST. 4 & 5 for patient metadata, clinical characteristics and treatment history). Each tumor was mechanically dissociated with BeadBug™ microtube homogenizer. Genomic DNA and total RNA purification from GBM patients using the AllPrep DNA/RNA Micro Kit (Qiagen).

### Short read whole-genome sequencing (srWGS) and copy number analysis

WGS DNA was produced using the KAPA Hyper Prep Kit (KAPA Biosystems, Roche, Basel, Switzerland) with 200 ng input genomic DNA. srWGS was performed at the Genome Sequencing Core Facility of Chang Gung Memorial Hospital, Linkou, Taiwan.

The raw sequencing reads were trimmed using fastp v0.23.2 (options: -r -W 4 -M 20 -l 36 -5 --cut_front_window_size 1 --cut_front_mean_quality 3 -3 --cut_tail_window_size 1 --cut_tail_mean_quality 3). Subsequently, the trimmed reads were aligned to the T2T-CHM13v2.0 genome using bwa mem v0.7.17 (option: -M -I 250, 150). The resulting SAM files were then converted to BAM format and sorted using SAMtools v1.15.1. After alignment, the duplicated reads were marked using Picard v2.27.4 (https://broadinstitute.github.io/picard/). General sequencing and processing metrics for all samples are provided in Supplementary Table ST.10. The genome-wide DNA copy number profiles of 36 GBM patients (Supplementary Table ST.11) were determined from the WGS data using readDepth v0.9.8.5. Pearson correlations of genome-wide copy numbers (exclude chrX. chrY and chrM) between different samples were determined. Copy numbers greater than 4 of 733 COSMIC (the catalogue of somatic mutations in cancer)^60^ cancer driver genes (v96) and 256 NCG (the network of cancer genes) oncogenes version 7.1. 737 defined were selected as the oncogene amplicons.

### RNA sequencing (RNA-seq) and data analysis

Frozen tissue samples (25 mg of GBM tumor tissue/patient) were homogenized using MagNA Lyser Green Beads (Roche, Basel, Switzerland) with 350 µl RLT plus buffer (QIAGEN, Hilden, Germany). The lysates were pipeted directly into a QIAshredder spin column, placed in a 2 ml collection tube and centrifuged for 2 min at full speed. The homogenized lysates were transferred to an AllPrep DNA spin column, placed in a 2 ml collection tube, and centrifuged for 30 s at 10,000 rpm. The flow-through was used for RNA purification. 1 volume of 70% ethanol was added to the flow-through, mixed well by pipetting, transferred to a RNeasy MinElute spin column, placed in a 2 ml collection tube, and centrifuged for 15 s at 10,000 rpm. 700 µl RW1 buffer was added to the RNeasy MinElute spin column and centrifuged for 15 s at 10,000 rpm. 500 µl RPE buffer was added to the RNeasy MinElute spin column and centrifuged for 15 s at 10,000 rpm. 500 µl 80% ethanol was added to the RNeasy MinElute spin column and centrifuged for 2 min at 10,000 rpm to wash the spin column membrane. The RNeasy MinElute spin column was placed in a new 2 ml collection tube, and centrifuged at full speed for 5 min. The collection tube with the flow-through was discarded. The RNeasy MinElute spin column was placed in a new 1.5 ml collection tube followed by adding14 µl RNase-free water to the spin column membrane and centrifuged for 1 min at full speed to elute the RNA. The RNA-seq library was constructed using the KAPA mRNA HyperPrep kit (KAPA Biosystems, Roche, Basel, Switzerland) with an input of 300 ng RNA. Sequencing data were generated by Illumina Nextseq 550 (Illumina, San Diego, USA) with 150 bp paired-end sequence.

The raw sequencing reads were trimmed using fastp^61^ (v0.23.2, options: -r -W 4 -M 20 -l 36 -5 --cut_front_window_size 1 --cut_front_mean_quality 3 -3 --cut_tail_window_size 1 --cut_tail_mean_quality 3). The trimmed reads were aligned to the T2T-CHM13v2.0 genome using HISAT^62^ (v2.1.0, option: --rna-strandness FR). The resulting SAM files were then converted to BAM format and sorted using SAMtools^63^ (v1.15.1). After alignment, the duplicated reads were marked using Picard v2.27.4 (Broad Institute, https://broadinstitute.github.io/picard/). Gene counting was performed using HTSeq^64^ (v2.0.3, options: -a 30 -s yes --secondary-alignments ignore --supplementary-alignments ignore --additional-attr gene_name). Total 40,806 genes were selected from UCSC GENCODE^65^ (v35) annotated in the following transcript biotypes: protein coding, processed, miRNA, lncRNA, misc_RNA, nonsense_mediated_decay, non_stop_decay, and retained_intron. The raw counts for all samples were generated using HT-seq count. To filter out low count genes, only genes with more than 5 raw counts in at least 2 samples were retained. General sequencing and processing metrics for all samples are provided in Supplementary Table ST.10. Counts were then normalized by upper quantile method to account for differences in read depth across samples. The raw counts were normalized by edgeR with the calculated k number (k=3) from RUVseq. Counts Per Million (CPM) was generated with edgeR^66^, and genes located on chromosomes X, Y, and M were removed to eliminate sex bias. ERCC spike-in controls were introduced to estimate the number (k=3) of unwanted variation factors for subsequent analysis, which was performed using RUVseq^67^. The design matrix was created based on sample conditions (short-survival vs. long-survival). Autosomal, protein-coding genes were then retained for Spearman’s correlation analysis, and the results were visualized with ggplot2 (Wickham, 2016).

### Long-read whole genome sequencing data generation

Library construction, sequencing and basecalling of 3 GBM cell lines and 20 GBM tumor samples were performed by Genome Technologies Service at The Jackson Laboratory, Connecticut, USA and BIOTOOLS Co., Ltd., Taipei, Taiwan respectively. In brief, 25 mg of brain GBM tumor was extracted using Monarch^®^ HMW DNA Extraction Kit (NEB) following manufacturer’s instructions. After extraction, the obtained genomic DNA was sheared with the 27G needle, ultimately yielding DNA fragments with a size of approximately 30 kb. The fragmented DNA was further enriched for fragments larger than 25 kb using the Short Reads Eliminator Kit (PacBio). DNA purity was checked using SimpliNano™-Biochrom Spectrophotometers (Biochrom, MA, USA). DNA concentration was determined by Qubit 4.0 fluorometer (Thermo Scientific) and the fragment size was monitored by fragment analyzer (Agilent technologies). The long-read sequencing libraries were constructed through the steps of end-repair, A-tailing, adapter-ligation. In brief, 100 femtomole of high molecular weight genomic DNA was end-repaired and added dA-tail using KAPA HyperPlus Kit (KAPA Biosystems, Roche), following the adapter ligation with Ligation Sequencing Kit V14 (SQK-NBD114, ONT) according to the manufacturer’s instructions. DNA libraries were clean-up by Long Fragment Buffer (LFB) washing to enrich DNA fragments >3kb. Libraries were sequenced with FLO-PRO114M flow cell (R10.4.1) on ONT PromethION 24 device. Sequencing raw data were decoded using Guppy (v6.5.7) with Super-accurate basecalling, 400 bps model. Reads with average quality above Q10 were considered as “pass” reads. Sequencing result checked by NanoPlot (v1.40.0) to validate read length profile. General sequencing and processing metrics for all samples are provided in Supplementary Table ST.10 and Supplementary Figure S5.

### Oncogene copy number and RNA expression heatmap

Integrated heatmaps represent oncogenes with recurrent high-level amplification (maximal copy number ≥5; n≥2 tumors), grouped by patient survival. RNA expression (CPM) and maximal copy number data are displayed using identical sample and gene hierarchies to ensure topographical consistency.

### Somatic SNV Calling, Mutational Signatures and TMB Analysis

Somatic single-nucleotide variants (SNVs) were identified from WGS data using Clair3^68^ and filtered for quality; variants present in the 1000 Genomes Project catalog were excluded to remove potential germline variants. Mutational signature exposures were attributed to the identified SNVs using SigProfilerAssignment^69^ (v0.1.9) based on COSMIC v3.4 SBS signatures^40^. For each tumor (n=12), the genome was partitioned into: (1) ecDNA, consisting of extrachromosomal amplicons aligned to chromosome 7, and (2) Chromosomal DNA, comprising autosomes excluding chromosomes 7, 9, X, and Y to minimize bias from the chromosomes of origin or sex-linked heterochromatin. Tumor Mutational Burden (TMB) was calculated by normalizing the somatic SNV count to the total size (Mb) of each genomic territory. Differences in TMB between ecDNA and paired chromosomal regions were evaluated using two-sided Wilcoxon signed-rank tests.

### Breakpoint validation

Primers for breakpoint junction validation were designed using Primer3 Plus (https://www.primer3plus.com/index.html). Primers were selected approximately 500-1000 bp upstream and downstream of the junctions. PCR amplification was performed across the junctions, and Sanger sequence was deployed to confirm the exact breakpoint sequences. Amplicon sizes varied between 400 and 2000 bp. Sequences of all primers used in this study are provided in Supplementary Table ST.3. PCR reactions were performed using 2X SuperRed MasterMix (Tools-biotech) in a 30 µl reaction volume and with 100 ng of genomic DNA as a template. PCR reactions were performed using the following parameters: initial denaturation at 95°C for 3 min, (95°C × 30 s, 58-62°C × 30 s, 72°C × 2 min) for 35 cycles, with a final extension at 72°C for 5 min. PCR products were analyzed on an Agilent High Sensitivity DNA Bioanalyzer, PCR products showing single bands were purified by Gel Extraction Kit (Qiagen). PCR products were sequenced using Sanger sequencing (Genomics).

To validate the *EGFR*, *PDGFRA*, *MDM4* and *CDK4* expression levels in selected tumors, RNA was extracted from GBM tumors using the AllPrep DNA/RNA Micro Kit (Qiagen). Quantitative real-time RT-PCR was performed using KAPA SYBR® FAST (KAPA Biosystems, Roche). Relative expression levels were normalized to *GAPDH* with controls from A11_P.

### Sequence Analysis of ecDNA Breakpoint Junctions

Breakpoint junction sequences were analyzed relative to the local genomic architecture. Direct and inverted repeat coordinates were retrieved from established datasets^70^. Repeats located within 1 kb of segment boundaries were mapped onto the structural reconstructions (Figure 6a). The enrichment of direct repeats, inverted repeats, and Z-DNA motifs was assessed against genomic background levels using permutation testing (1,000 simulations). To identify non-canonical structures, a 100-bp window flanking each junction was manually inspected for imperfect inverted repeats (allowing for at most 3 non-canonical base pairing or mismatches or gap), which are detailed in Figure 6b. Recurrent breakpoints were defined as those located within 10 kb of one another across different samples (Supplementary Table ST.7).

### Replication Timing Analysis

To investigate the temporal landscape of ecDNA formation, high-resolution replication timing (RT) maps—derived via Repli-seq across 16 S-phase fractions^50^ (S1–S16)—were utilized. Genomic coordinates for ecDNA segments (length ≥100 kbp) were mapped to these RT profiles to calculate empirical mean replication signals for ecEGFR and ecPDGFRA assembly segments. The statistical significance of the observed RT signatures was evaluated using a permutation-based approach. For each cohort—including the internal GBM discovery cohort, GLASS, and PCAWG^6^—we generated a null distribution by performing 10,000 Monte Carlo simulations. During each iteration, empirical ecDNA segments were randomly shuffled across their respective chromosomes using BedTools shuffle, maintaining segment length and chromosomal context. The empirical mean RT for each S-phase fraction was then compared against the distribution of 10,000 simulated means. P-values were determined by the frequency with which simulated means exceeded or fell below the empirical observations (#sim > emp). Enrichment within specific temporal windows (e.g., mid-to-late S-phase, S7–S10) was defined as a significant deviation from the simulated background model, ensuring that the observed temporal signatures were not a stochastic property of the local genomic architecture.

### Single-cell multiome procedure

30-250 mg of GBM tumor tissue were used to perform nucleus isolation using the nuclei Isolation Kit (10X Genomics). The nuclei were examined under high power microscope to check their integrity to be free of debris. 10,000 nuclei were used to go through enzyme reaction, mononuclear formation, and labeling by barcode. After QC, ATAC library and gene expression library were prepared separately.

### Single-cell multiome data processing

The cellranger-arc (version 2.0.2) software was used for alignment, filtering, barcode counting, peak calling, and counting of both ATAC and GEX molecules. The cellranger-arc output data were processed using the Seurat (version 5.2.1) and Signac (version 1.14.0) R packages. Use Signac to calculate the NucleosomeSignal and TSSEnrichment. TSS positions were acquired from the T2T GTF file used by the cellranger-arc. Filter cells by 1,000 < nCount_RNA < 25,000, 500 < nCount_ATAC < 100,000, nucleosome_signal < 2, TSS.enrichment > 1. General sequencing and processing metrics for 29 tumor samples are provided in Supplementary Table ST.12. scRNA data were normalized by using SCTransform to normalize the sequencing depth across all cells and using PCA to reduce the data dimension. scATAC data were normalized by using RunTFIDF to normalize sequencing depth and peak frequency and reduced dimension using RunSVD. To integrate the scRNA and scATAC modalities, weighted nearest neighbors (WNN) analysis was performed using FindMultiModalNeighbors. Doublets were filtered using scDblFinder package (version 1.16.0). RNA-based doublets were identified using the scDblFinder method. scATAC-based doublets were detected using AMULET. scATAC fragment data in the known blacklist regions (lifted over from the hg38 v2 blacklist^71^ and sex chromosomes were excluded. P-values from both methods (scDblFinder and AMULET) were combined using Fisher’s method. Cells with a combined p-value > 0.05 were retained as singlets.

### Single cell resolution of ecDNA copy number analysis by single-nucleus ATAC-seq coverage

We followed the protocol described^36^. In brief, we created a 3 Mb sliding window every 1Mb across the T2T reference genome and calculated the GC content in the windows. The reads mapping to the regions in the T2T blacklist (lifted over from the hg38 v2 blacklist^71^ were excluded. Transposition events within each genomic window were then counted. For each window, calculated the events per bp and calculated the fold change to the 100 nearest windows according to the GC content. Calculated the mean log2[fold change] of nearest windows and used the formula copy-number = 2 x 2mean log2[fold change] to compute the copy-number. For gene copy-number calculation, we used windows overlapped with the gene coordinate and calculated the mean copy-number of those windows. For each ecDNA locus, we used the Gaussian Mixture Models (GMM) to group the cells into high-copied and low-copied groups and used Chi-squared test to determine the significance of co-existing of multi-species ecDNAs.

## SUPPLEMENTAL INFORMATION

### Supplemental Tables

ST.1: ecDNA assembly summary

ST.2: assembly subspecies

ST.3: junction validation

ST.4: patient metadata

ST.5: study metadata

ST.6: ecDNA CN variablity

ST.7: ecDNA breakpoints

ST.8: Replication timing signature

ST.9: ecDNA novel insertions

ST.10: sequencing summary

ST.11: copy number summary

ST.12: 10X Multiome Statistic (T2T-CHM13)

### Supplemental Figures

**Supplementary Figure S1 | ecLego3 demonstrates enhanced specificity and structural fidelity in ecDNA reconstruction from CHP-212 long-reads. (a)** Comparison of high-abundance amplicon reconstructions by CoRAL (colored arcs), Decoil (dark gray blocks), and ecLego3 (light gray blocks). The top panel displays long-read alignment coverage (black trace; peak depths of 154x and 144x indicated) alongside GENCODE v42 gene annotations within chr2: 15,942,184–15,946,347 region, hg38. Structural variation, including insertions (purple) and deletions (red), are correctly identified in the ecLego3 assembly but are omitted in the Decoil reconstruction despite presence in the raw reads. **(b)** Resolution of consensus ecDNA sequences by ecLego3. Bar plots indicate variant frequencies (red: alternative; blue: reference) at five genomic positions within the *ecMYCN* region, demonstrating the accurate capture of sub-species heterogeneity. **(c)** Evaluation of assembly specificity. Two additional ecDNA candidates reconstructed by Decoil lack corresponding long-read coverage support (zoomed insets) are shown, highlighting ecLego3’s specificity compared to existing methods.

**Supplementary Figure S2 | Comparative reconstruction of complex ecDNA architectures in B171 GBM model. (a)** Genomic landscape and ecLego3-reconstructed assemblies of *ecMDM4* and *ecPIK3C2B*. Tracks (top to bottom): long-read coverage; UCSC gene annotations; and alignment of two circularized ecLego3 assemblies of *ecMDM4* and *ecPIK3C2B* to the T2T-CHM13 reference genome. **(b)** Evaluation of Decoil-based ecDNA assemblies. Top track: split-read coverage at structural junctions. Middle and bottom: alignment of two representative Decoil-assembled ecDNAs to the T2T-CHM13 genome. Unfilled red ribbons denote segments in the Decoil assemblies unsupported by split-reads, specifically within the region of the partial *MDM4* duplication (exons 1–5). This absence of split-read support for Decoil-assembled segments highlights the algorithm’s limitations in accurately resolving complex, multi-segmented rearrangements compared to the high-fidelity reconstruction achieved by ecLego3.

**Supplementary Figure S3. Validation of copy number (CN) and transcript abundance of oncogene amplification in GBM tumors. (a,b)** Genomic CNs and relative mRNA expression levels of *EGFR* across a subset of GBM tumor samples, as determined by quantitative PCR (qPCR) and RT-qPCR, respectively. **(c,d)** Validation of *PDGFRA* CN and relative expression levels. **(e,f)** Validation of *MDM4* and *CDK4* CN and transcript abundance in selected GBM tumors. For all expression plots, data were normalized to *GAPDH* and are represented relative to the A11_P control sample. Error bars denote the standard deviation (SD) from experiments performed in triplicate. Statistical significance was determined by Student’s t-test (*p* < 0.05). **(g)** The list of oligonucleotide primer sequences used for qPCR and RT-qPCR assays.

**Supplementary Figure S4 | Structural heterogeneity, evolutionary trajectories, and transcriptional consequences of ecDNA subspecies. (a)** Numbers of subspecies found per CG between primary and recurrent tumors. **(b)** Linearized structural map of the 1.33 Mb *ecEGFR* CG in A15_P. Eleven distinct subspecies (P1–P11) are defined by combinatorial SVs across genomic segments A–H. ΔC: a 16 kb deletion (Δ exons 2–7) within the *EGFR* locus. Pink bars at the bottom denote super-enhancer (SE) elements. **(c)** Reconstructed molecular phylogeny illustrating the parsimonious evolution of ecDNA subspecies in A15_P. Nodes represent individual subspecies; values indicate relative copy number abundance (*c*, estimated copies per cell). Edges denote the hierarchical acquisition of specific SVs (e.g., ΔG indicates loss of segment G as defined in **(b)**. **(d)** Genomic architecture and transcriptional profiles of 10 ecDNA subspecies in A23_P (total circle copy number = 62). Top track: lrWGS read coverage (blue) across the *EGFR* locus. Middle/bottom tracks: Grey and colored arrows represent the structural configurations resulted from inversions and deletions, respectively with copy numbers (*c*) indicated for each subspecies. Pink bars denote SE regions. Note that deletions (represented by dashed lines) removing these SE-containing segments are restricted to minor subspecies (e.g., P9), while dominant subspecies retain the full SE repertoire, underscoring their functional selection.

**Supplementary Figure S5 | Long-read sequencing quality control and performance metrics. Violin plots showing the distribution of primary sequencing statistics for the ONT WGS cohort (*n*=21). (a)** Mean read length (median≈27 kb). **(b)** N50 read length (median≈37 kb). **(c)** Genomic depth of coverage (median≈32×). **(d)** Alignment accuracy (median≈97.6%). Internal boxplots indicate the median (white line) and interquartile range (IQR; dark grey box); whiskers extend to 1.5 × IQR. Kernel density estimation (blue) represents the distribution density of the samples.

## Notes

### Competing Interest Statement

The authors have declared no competing interest.

