## Supplementary Figures for "An Atlas of Extrachromosomal DNA Structures Illuminates Its Evolution and Biogenesis in Cancer"

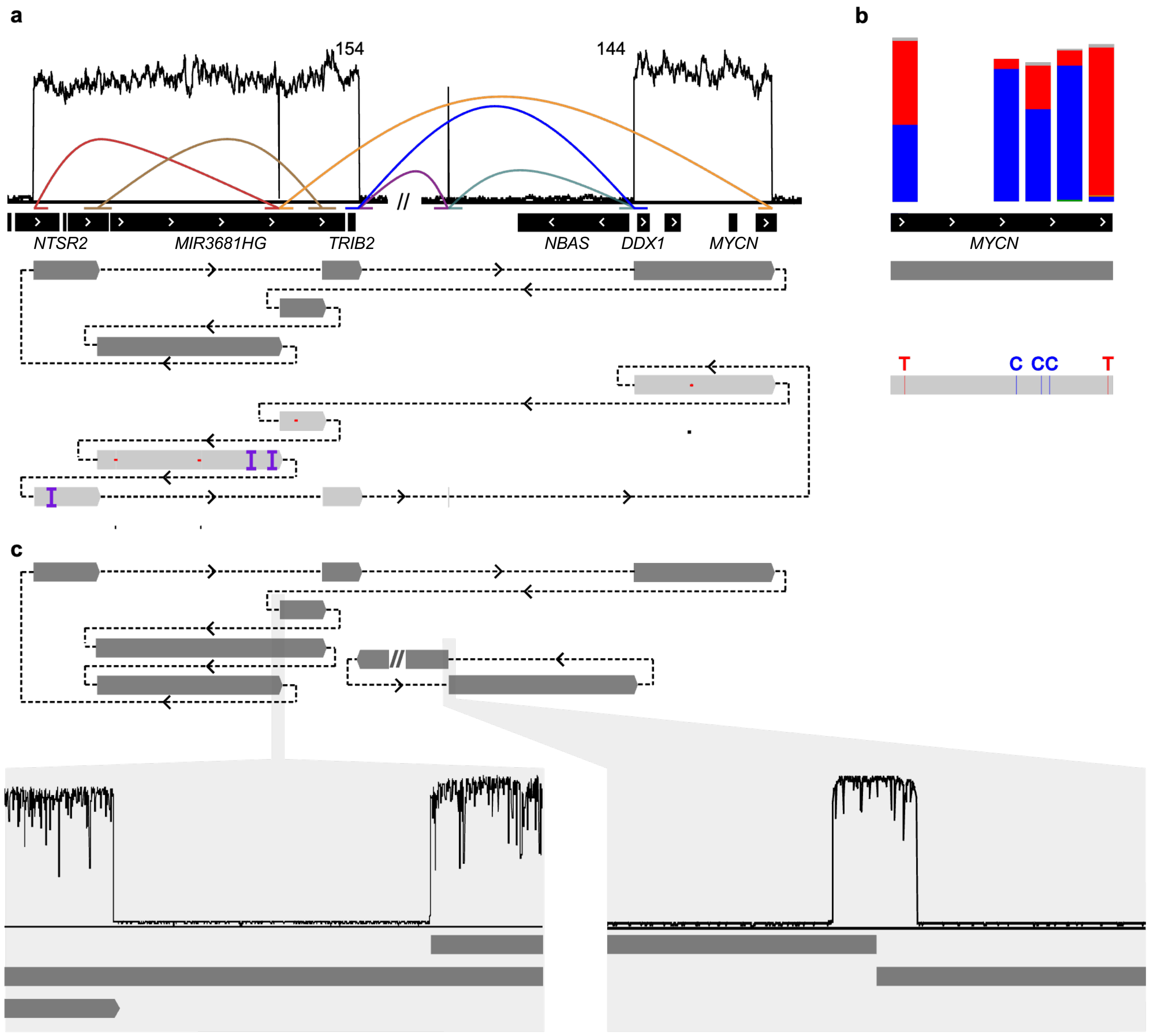


**Supplementary Figure S1 | ecLego3 demonstrates enhanced specificity and structural fidelity in ecDNA reconstruction from CHP-212 long-reads. (a)** Comparison of high-abundance amplicon reconstructions by CoRAL (colored arcs), Decoil (dark gray blocks), and ecLego3 (light gray blocks). The top panel displays long-read alignment coverage (black trace; peak depths of 154x and 144x indicated) alongside GENCODE v42 gene annotations within chr2: 15,942,184–15,946,347 region, hg38. Structural variation, including insertions (purple) and deletions (red), are correctly identified in the ecLego3 assembly but are omitted in the Decoil reconstruction despite presence in the raw reads. **(b)** Resolution of consensus ecDNA sequences by ecLego3. Bar plots indicate variant frequencies (red: alternative; blue: reference) at five genomic positions within the *ecMYCN* region, demonstrating the accurate capture of sub-species heterogeneity. **(c)** Evaluation of assembly specificity. Two additional ecDNA candidates reconstructed by Decoil lack corresponding long-read coverage support (zoomed insets) are shown, highlighting ecLego3's specificity compared to existing methods.

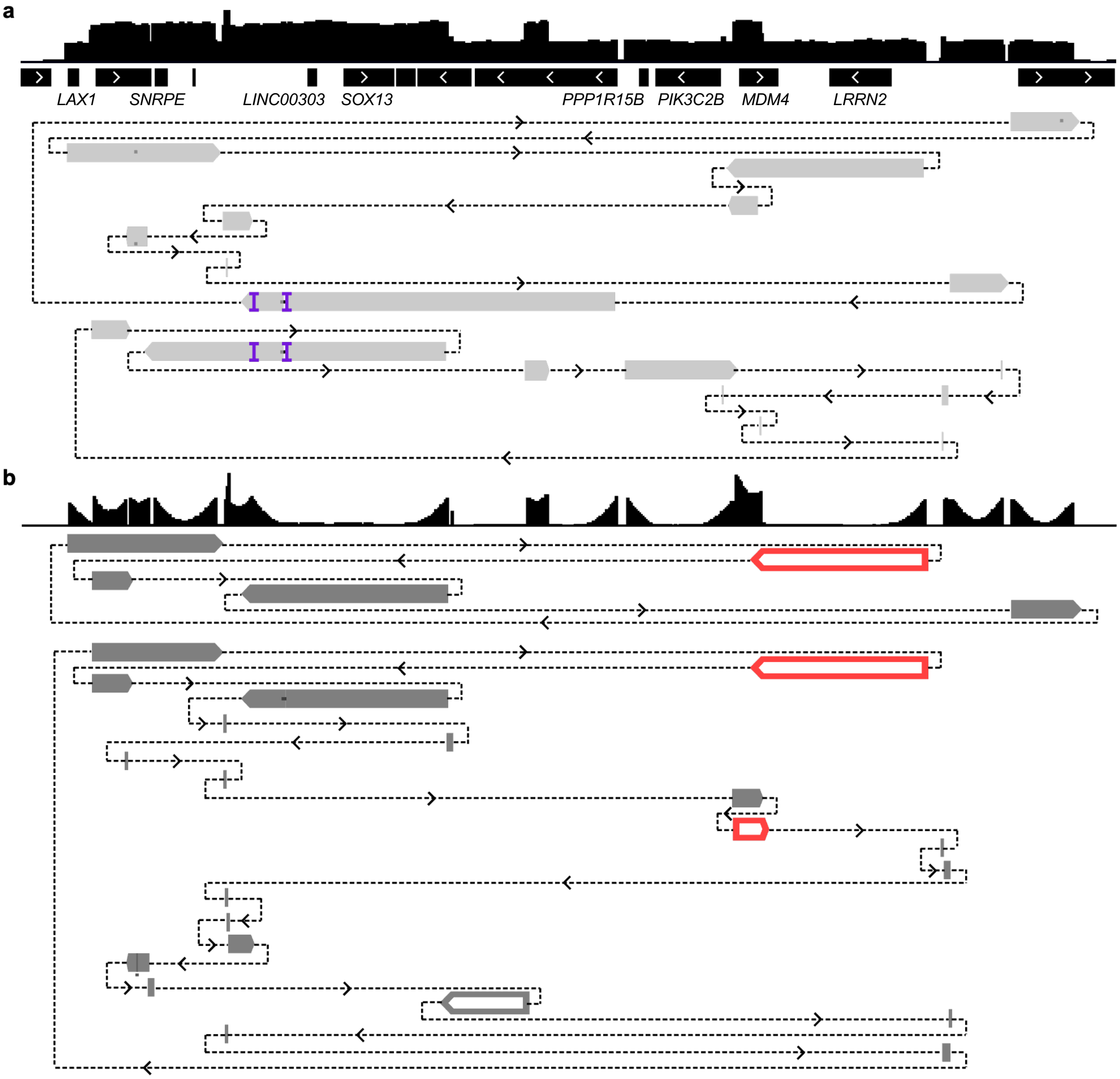


**Supplementary Figure S2 | Comparative reconstruction of complex ecDNA architectures in B171 GBM model. (a)** Genomic landscape and ecLego3-reconstructed assemblies of *ecMDM4* and *ecPIK3C2B*. Tracks (top to bottom): long-read coverage; UCSC gene annotations; and alignment of two circularized ecLego3 assemblies of *ecMDM4* and *ecPIK3C2B* to the T2T-CHM13 reference genome. **(b)** Evaluation of Decoil-based ecDNA assemblies. Top track: split-read coverage at structural junctions. Middle and bottom: alignment of two representative Decoil-assembled ecDNAs to the T2T-CHM13 genome. Unfilled red ribbons denote segments in the Decoil assemblies unsupported by split-reads, specifically within the region of the partial *MDM4* duplication (exons 1–5). This absence of split-read support for Decoil-assembled segments highlights the algorithm's limitations in accurately resolving complex, multi-segmented rearrangements compared to the high-fidelity reconstruction achieved by ecLego3.

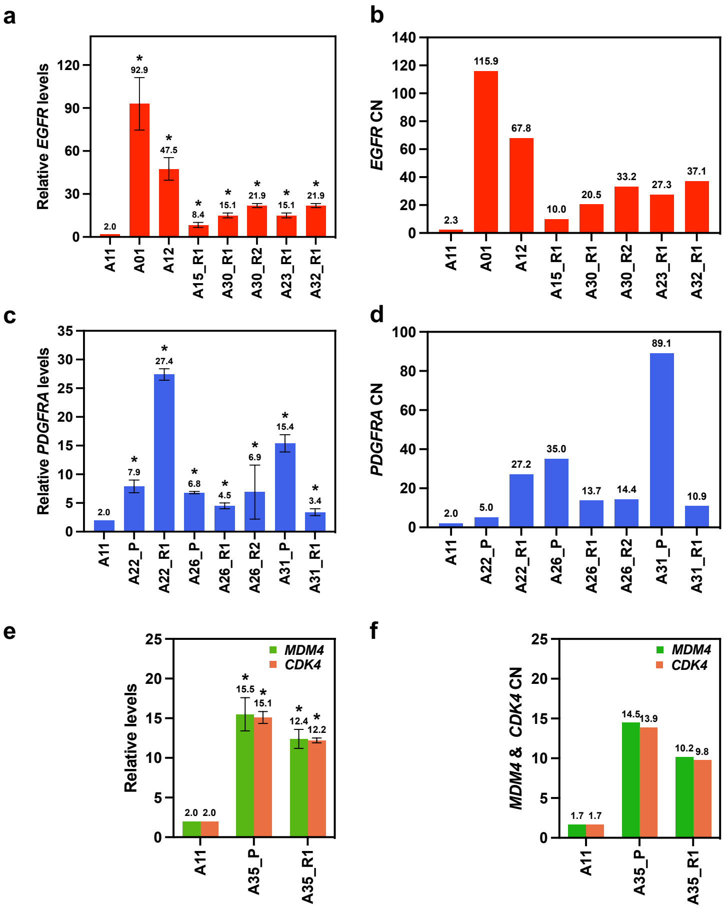


g

| Primers | Sequence (5' to 3') |
|---|---|
| PDGFRA-F | TGATGGCTTGCTCAGAGGAC |
| PDGFRA-R | TCCTGGCCGACATGATGAA |
| EGFR-F | AACCTCTGGCAAACTCGCT |
| EGFR-R | TGCCACTTTCTGGTTGGTCA |
| MDM4-F | ACTCTTCTGAGCACTGTGAGTTAT |
| MDM4-R | AATGGTTCTGGAGTGTGGCA |
| CDK4-F | TTCGCGAGCTGGTTATGGAA |
| CDK4-R | CTGCAGCAACAAAGGGACTC |
| GAPDH-F | GTCTCCTCTGACTTCAACAGCG |
| GAPDH-R | ACCACCCTGTTGCTGTAGCCAA |

**Supplementary Figure S3. Validation of copy number (CN) and transcript abundance of oncogene amplification in GBM tumors.** (**a,b)** Genomic CNs and relative mRNA expression levels of *EGFR* across a subset of GBM tumor samples, as determined by quantitative PCR (qPCR) and RT-qPCR, respectively. **(c,d)** Validation of *PDGFRA* CN and relative expression levels. **(e,f)** Validation of *MDM4* and *CDK4* CN and transcript abundance in selected GBM tumors. For all expression plots, data were normalized to *GAPDH* and are represented relative to the A11_P control sample. Error bars denote the standard deviation (SD) from experiments performed in triplicate. Statistical significance was determined by Student's t-test ($p < 0.05$). **(g)** The list of oligonucleotide primer sequences used for qPCR and RT-qPCR assays.

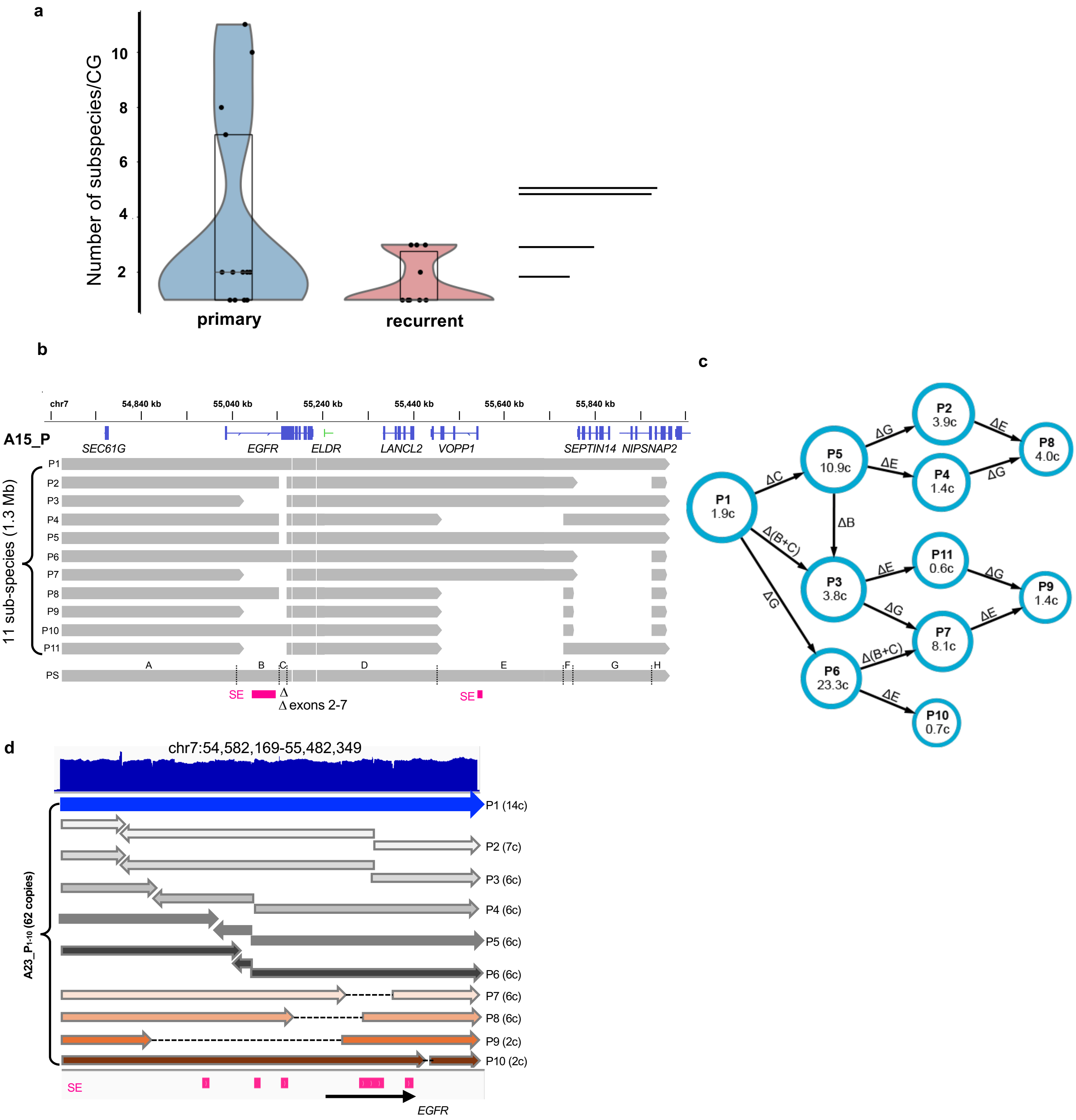


**Supplementary Figure S4 | Structural heterogeneity, evolutionary trajectories, and transcriptional consequences of ecDNA subspecies. (a)** Numbers of subspecies found per CG between primary ($n$=13) and recurrent ($n$=9) tumors. **(b)** Linearized structural map of the 1.33 Mb *ecEGFR* CG in A15_P. Eleven distinct subspecies (P1–P11) are defined by combinatorial SVs across genomic segments A–H. ΔC: a 16 kb deletion (Δ exons 2–7) within the *EGFR* locus. Pink bars at the bottom denote super-enhancer (SE) elements. **(c)** Reconstructed molecular phylogeny illustrating the parsimonious evolution of ecDNA subspecies in A15_P. Nodes represent individual subspecies; values indicate relative copy number abundance ($c$, estimated copies per cell). Edges denote the hierarchical acquisition of specific SVs (e.g., ΔG indicates loss of segment G as defined in **(b)**. **(d)** Genomic architecture and transcriptional profiles of 10 ecDNA subspecies in A23_P (total circle copy number = 62). Top track: lrWGS read coverage (blue) across the *EGFR* locus. Middle/bottom tracks: Grey and colored arrows represent the structural configurations resulted from inversions and deletions, respectively with copy numbers ($c$) indicated for each subspecies. Pink bars denote SE regions. Note that deletions (represented by dashed lines) removing these SE-containing segments are restricted to minor subspecies (e.g., P9), while dominant subspecies retain the full SE repertoire, underscoring their functional selection.

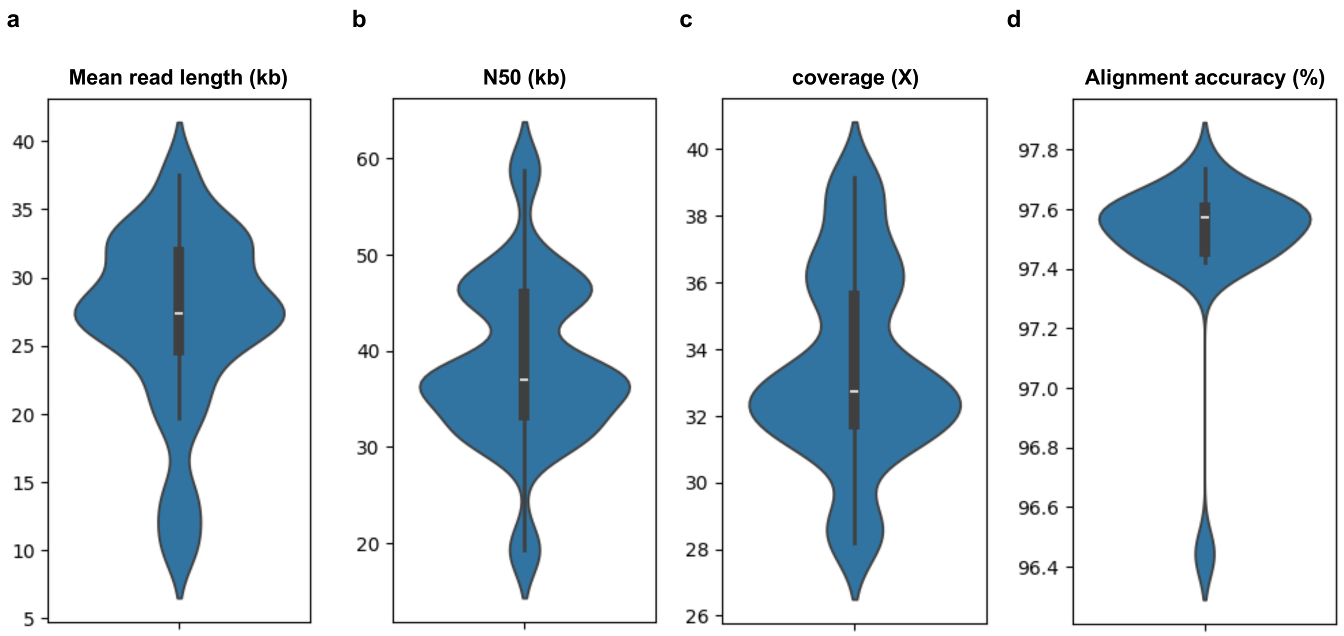


**Supplementary Figure S5 | Long-read sequencing quality control and performance metrics. Violin plots showing the distribution of primary sequencing statistics for the ONT WGS cohort (*n*=21). (a)** Mean read length (median≈27 kb). **(b)** N50 read length (median≈37 kb). **(c)** Genomic depth of coverage (median≈32×). **(d)** Alignment accuracy (median≈97.6%). Internal boxplots indicate the median (white line) and interquartile range (IQR; dark grey box); whiskers extend to 1.5 × IQR. Kernel density estimation (blue) represents the distribution density of the samples.
