## Supplementary Information for "An Atlas of Extrachromosomal DNA Structures Illuminates Its Evolution and Biogenesis in Cancer"

Supplementary information for the inserted sequences (bold) in the validated junctions with 10 bp genomic sequences at the breakpoints on both 5' and 3' ends.

CHP-212_156bp_cG12.1.1

**CTACCTTTTGATTTAAAGTGAGAGACCTGAGACTCTTCCTTTCACTTGAATGCTTAGAGAC**
**CATTGTAGGGTTATTAACTGGCCTAATTTCAGTATTGCTGTGTCTCAGGGAATAGGGAGGC**
**CCAAGGAGAGGGGGAGAGAGATGGGAGAATGGCC**

A01_P_4.3kb (4297bp) CG1.1_chr1:223184658-chr7:55923308

ATACATCAAC**TCTGATGACCTTTGTTTCCACTCCCCTTAAGCCATTCACAGGAACAGCTACT**
**TTACATAGAAAAGTCAATCTTGGAGGGTTGAGTAATTTGCCCAATCACTGTAAAAGGTAGA**
**GCTTCAATTATCACCCAGGCCCATCTGTTGCTGGTATTCCCTCTCCACCCCTCCGTGTGGC**
**TTCCCTTTATTTATTTATTTTAATTTTTTTTGAGATGGAGTTGCACTTTTGTTGCCCAGGCTGG**
**AGTGCAATGGCATGATCTCGGCTCACTGCAACCTCCACCTCCCAGGTTCAAGCAATTCTC**
**CTGCCTCAGGCTCCCAAGTAGCTGGGATTACAGGTGCCTGCCACCAAGCCCAGCTAATTT**
**TTTGTACTTTTAGTAGAGATGGGGTTTCACCATGTTGACCAGACTGGTCTTGGACTCCTGA**
**CCTCAGGTGACCCACCCTCCTCAGCCTCCCAAAGTGCTGAGATTACAGGCGTGAGTCAC**
**GGTGCCTGGCCTTGGCTTTCCTTTAATAAATCACCACTCAATTTGATGTGGAGGCTCTGTT**
**CATTCCCAGAATATAATGAATGGATTATATATATATTTTTTCTTTTTGAGACAGAGTCTTGCTT**
**TGTCTCGCAGGCTGGAGTGCAGTGGCGTGATCTCAGCTCATTGCAGCCTCTGCCTCCCAG**
**GTTCCAGCAATTCTCCTGCCTCAGCCTCCTGGGTAGCTGGGATTACGGGCGCTCACCACC**
**ATGCCCAGCTAATTTTTTTTTTTTTTTTTTTTTTTTTTTTTTTCAGGGTGGGGCTAACTCCTGA**
**CCTCTGGTATCCGCCCCCTCGGCTTGGAAAATTCTTGGATTATAGGCATGAACCCCCGCCC**
**CCAACCTTAACATTAAAAAGATTTAAAAATTGGTAACAACCAGGGGCAGTGGGTTCCCCC**
**TGTAATCCCAGCACTTTGGGAGGCCGAGGTGGGCAGATCACCTGAGGTCAGGAGTTCAA**
**GACCAGCCTGGCCAACATGGTGAAACCCCGTCTCTAATAAAAATAAAAAATAAAAAAAAC**
**AATTAGCTGGGTATGGTGGCACGTGCCTATAATCCCAGCTACTCAGGAGGCTGAGGCACG**
**AGAATCACTAGAACCCAGGAGGTGGAGGTTGCAGTAAGCCAAGTTCGTGCCACTACCCT**
**CCAGCCTGGGCAACAGAGTAAGACTCCATCTAAAAAAAAAATGAAGAAGAAGAAATTAG**
**TGTAGTGTGGGAAGTGAAAAAAAAAAAAAAAAAAAGGAAAAAAAAAAAGGATTGAATTCTT**
**GAAAACCCTCTTATGGGGCCGGCCCCGACTTTGACCCAAATTAAATGGGTTTAATAGGCA**
**AGGGGGGGATCTTTTCATAATTTTATTTGATGCCTAAAATACTTTTATCTTTTTTTCTTATAGG**
**AGGAAAATTCTTCCAAGGAGGATCTCCCACTCAAAGCTGAGGAAGCTTTTCTACTCAAAA**
**AAGGCTGGGGGTTTTGATGTTGACAGCACGGCCATCAGGGAAAAAGGAATCGGGGAGCT**
**AGCCAAAATCTGTGGGGTAAAGGATGCGGGGTCAAAAAGGTAGGGATAGCATTTATTCCC**
**TTTATGAAAGGATAAAAAATTTCTTTTTTTTTTTTTTTTTTTTTTTGAGACAGAGTCTCACTCT**
**ATTGCCCAGACTGGAGTGCAGTGGCACAATCTTGGCTTACTGCAACCTCCACCTCCCAGG**
**GTTCAAGCGATTATCCTGCCTCAGCCTCCTGAGTAGCTGGGATTACAGGCGTGTGCAACC**
**ACATCCGGCTAATTTTTTTATTTTTAGTAGAGATGGAGTTTCACCATGTTGGTCAGGCTGGT**
**CTCAAACTCCTGACCTCGTGATCCACCCACCTCAGCCTCCCAAAGTGCTGGGATTACAGG**
**CGTGAGCCACCATGCCCGGCCATGAAATGATAAAGAATTTCTAAAGAGTGGCTGTTTTGG**
**ATGAAGTGCTGGACCCTGGCTATGGAACATGAGCACTAGGCTTTTTCTCTCACCCCTTAGA**
**GTTGAATTTGATATATTCATTCATTTATATAAATATTTATGGCAAAAAAATATGGATAAGACAT**
**GTTCTTAGCTTGATTAGGGGAATGATGACCATCACTAACCAAAGAAGATGGCCTAGGGGG**
**CAGGGCTCAAGTGTCCAGTGGCTTTGTGTCATGGTGAAAGTGTGACTCCGTTGAAAGTGT**
**GGCCTTAGTGGTTCCCTGGAAAGGTTCAGAGTCTCTTGGTGGATCCTGGGTCAGAGCCCC**
**TCTCTCCCTCCCTCCCTCCTGCCCTCCCACCTCCTGCCCTGCAGCTGGGCACCACCCTCT**

**GCAGCCCCAGTCCCCCAGTCATGCACCATGTCATTTTCTTTTTTTTTTTTTTTTCAGGACGG**
**AGTCTCGCTCTGTCACCAGGCTGGAGTGCAATGGTGCAATCTCGGCTCACTGCAACCTCC**
**GCCTCCTGGGTTCAAGCAATTCTCCTGCCTCCACCTCCTGAGTAGCTGAAACTACAGGCA**
**CGCACCACCACACCCGGCTTATTTTTGTATTTTTAGTAGAGATGGTGTTTCACCATGTTGGC**
**CAGCCTGGTCTTGAACTCCTGACCTCGTGATCTGCCGGCCTCGGCCTCCCAAAGTGCTG**
**GGATTACAGGCATGAGCCACCACACTCGGCCACCATTTTATTTTCAACTCCCTTTCATGGA**
**AGAACGTTTAGCCTTTGGTCTCTTTCTTGATTTATGACAGCTCGCGGCTTTCAAGAAAACT**
**ACCTATGAATAGGCTGTGTAACTTTTATTTATTTATTTATTTTTTGAGATGGAGTCTCTGTCTT**
**CTAGGCTGGGGTGCAGTGGCATGATCTCGGCTCACTGCAACCTCCCCCTCCCAGGTTCC**
**AGCAATTCTCCTGCCTCAGCTTCCCAAGTAGCTGGGATTACAGGCATGCACCACCACACC**
**TGGCTAATTTTTGCATTTTTTTTTATTAGAGATGGGGTTTTGTCATGTTGGCCAGGCTGGTCT**
**TGAACTCCTGGCTTCAGATGAGCTGCCCACCTCAGCTTCCCAAAGTTTTGGCATTATAGGC**
**ATGAGTCACTGCACCCAGCTTGTTAATGTTATTTTCAAGCACACCTTCAAAAGTTTATTCCA**
**AGGCCTTGCTCTCATAGCAGCAAAGCCTGTCTGTTGCATAGTGGGGCTCTTGTTAGCATCT**
**TCCTTTTGGTGGATGTTGTGTAGACCCGGGGCAGTAAGGCATGTTAACCTCGAGGACATT**
**GGACCTGGCTGGCCTGACTGTTGGGTCTCCCTCCTAGGACATGGCGAGCCATGGGTGGG**
**GCAGTGTCTTTCAAAGCTGCTCTCACGGAGCACTTAGCCCCAATCCAGCCCTCCAGGGA**
**GCAGGTGCAGAGACTCATAGCAGAGCACCCCCCACACCTGACCCCCAGCATAAGTAAGA**
**GGAGCCGCTGCTCCAGGTGTATTTCAGCACCAGTGTTGGGGGGCATGTCCTCCTGAGAG**
**CATCTAACGATTGCTTTTAGAGAGCCCTCTGGGTGGTTTATTTATTTCATTTTTTATTTTTTAA**
**TTTTATTATTTTAAAAAAAATTATTGATTGATTTTATTTTATTTATTTATTTATTTATTGAGGAAA**
**GTCTGGCTCTGTCGCCCAATACGGAGTGCAATGTTACGATCTCGGCTCACTGCAACCTCT**
**GCCGCCCGGGTTCAAGTGATTCTCCTGCCTCAGCCTCCTGAGTAGTTGGGATTTCAGGCA**
**CCCACCATGCCAGGCTAGTTTTTGTATTTTTAGTAGAGACAGGATTTCACCATGTTGTTTAG**
**GCTGGTCTCGAACTCCTGACCTCAGGTGGTCTGCCCGCCTCAGACTCCCAAAGTGCTGG**
**GATCACAGGCGTGAGCCACTGCGCCTGGCAAAATTTTATTTATTTATTTTTTTTGAGACAAA**
**GTCTCACTCTGTCGCCCAGGGTGGAGTGCAGTGGTGCAATCTCAACTCACTGCAACTTTT**
**GCCTCCCAGGTTCACACGATTTTCCTGCCTCAGCCTCAGGAGTAGCTGGGATTACAGGCG**
**CCTGTCCCCACACCTGGTTAATTTTTGTATTTTTAGTAGAGATTGGGTTTTCCCATTTTGGC**
**CAGGGGGGTTTTGAATTTTTTTTTTCAAGGGATCCCCCCCCAGCCTCGGTCTCCAACGTG**
**CTGAGATTACAGGTGTGAGTCACCGCACCTGGCTTATTATTTTTTTTTTTTAGATTCAAAGTC**
**TCATTCTGTTGCCCAGGTTGGAGTACAGTGGTATAATCATGGCCCACTGCAGCCTCGAACT**
**CCTGGGGTCAAGTAGCCCTCCCACCTTAGCCTCCTTAGTAGTAGGGACTG**CAGGTGAATG

A15_R1_475bp_CG1.1_chr7:56180237-54972777

GGGCGGATCA**TGTATATAAAATTGTATAGTATATACTCATTTTTGTCTGGCTTCTTTCACCCA**
**GCGTATTATTCTGAGATTTGTCCATGTTGTGTGTATCAGTAGTTCATTCTTTTTTGTTGCTGA**
**GTAGCATTCAGTTATATGGACAGACCATATTCTATTTGTCCATTCACTGATTGATGGTCATTT**
**GGGTTGTTTCCAGTCTTTGGCTGTGACAAATAAAGCTACTATGAACATGTGTGTACAAGTC**
**TTTGTGTAGACGTACACTAAGGCAGGGAATCTCTGGAGCCCAGGAGTTTGAGGCTGCAGT**
**AAGTTATGACTGTGCCACTGTGCTCCAGTCTGGACAACAGAGTGAGACCCTGTTTCTAAA**
**AAAACAATAAAAATAAGTGACTTAAGATACAGCAGCTAGTTATCCTTAGAGATGTCACTGG**
**ATTATGAACTGATAGGCTCATGTTTATGTTTATAAATAGTATAAATAATATGTA**TAATAACTCC

>A15_R1_81.6kb (81679bp) CG1.3.1_chr7: 54960334-54960678

ACTTTCCATTTTTTATGGGCAACCTGGATATGGCTGGGGTGGGCCTCACTGTGTCCTGAAT
CCTGGAACTGGACCTGCATGTGGCTGGGGTGGGCCTCGCTGAGTCCTGAGTCCTGGAAC
TGGACCTGTGTGTGGCTGGGGTGGGCCTCACTGAATCCTGAGTCCTGGAACTGGACCTG
GATGTGGCTCGGGTGGGCCTCACTGTGTCCTGAATCCTGGAACTGGACCTGCATGTGGC
TGGGGTGGGCCTCGCTGAGTTCTGAGTCCTGGAACTGTATCTGTGTGTGGCTGGGGTGG
GCCTTGCTGAGTCCTGACTCCTGAGCAGGGGACATTGGAGAAAGGACAGGATGCTGGCT
TCGATTCAAAATCCCTCTGTGTGAGAGGATAGATAGCAGTGAGCGTGCTGGGCCTGGCT
TTCCTTCCTAGACAGAGGGTGAGTAAACGCCTCCTCTCGGGCCAGCATTCTACTTGTCTA
ATCTCTGCGCTGAGACTTTTGCTTACTCATGTTCAGGGGCTCTCTCTTTACACACAGGCAT
ACTTTTGAACTGAATAATACCACTTCCCCTCCAAAAGTCTGTGGGCCAGGCCGGTGCTTG
CACTCTGTGCCTCCACTGTGTCCAGGGCATCTGCCTTGCTGCTTCAGGGCCGTGTGGCTC
CTCCAGTGGCGCCTCAGACCTGCCAGAGAAGGGCAGTGCAGAGCCAGGGCGGCAGGA
GCCCCAGCAGAGCTGGCCTCTCCCTTTCTGGCTCGCTGGGGAGCCGACAGCACCACAG
AACCCGGGGTGGGACAGAATGGGGCAGTGTGGACGTGGGGATGACGGAGATTGCATGA
TTATCACCAAAGGTGAGCTCAGAGACATCTGGTAGAAGGTAGGGGGAGATACAGAATTC
CTAAATGGGGTTGGCATCAAAGATCACTGCTGGAAAATCGGCTTCAAAACCCGGCCAGC
TGACTTCAGGAAGCCAGCTCTTTGTTCAAAGTCAAGACTGCCCTGTGCTACCTCCCTTCC
CCTCTGACTTCACGTCCCTCTTTGTTACTGGAGAGCTGGTGAGCACCAGGCAGTAGAAA
GGTGCCAAGGGCTGCTGGGGGCTGCAGACTGCACAGCCCAGCCCAGCCTCCACCACCC
CAGCAGGCTCACTGGGCAGCTCACTCTTTCTTCCTCAGCTTTGCATGAGACAGCTCAGTG
CTGCCTGCCAGGGTGCAACTCTTCTCTTCCTAGACAGGCCCTCACAACCGAAGGCACCC
TAGCAAGGTGATGTGCAGGTAAAGACAGGCAGGGCTGCCGGGAGTTCACCTCGCTGGG
CGCTGTCTCACCCAGTCCACACACCTTTTCTCCTCTATCTTCTATTCAGTTCAGGCCCGTG
ACTCACTCCTTGATGACAAAGGCTGAGTAATATTTCCCATATGAAGCTGTTTATTTGCTGG
TTGAGCTGTCGCCCTAGAAGTCATGTGAATACCAAGCATGGCTGAGAAGCTCCGTGGCC
AGATGGAGTCAGCCACAGAGGTTGCACGATGACCCTTTGGCCAGGCTCACTCCACCCTG
CTCTGAGATGCACACCAGGCACCAAAGTTGCTTGTTCTTTTGTTCATTAATTCAACTACAA
CATATCTAGTAAAGTAGCGCGAAGGTAGCTGCTAAGGATGAATCAGAAGTACTCACTGC
CCTGAAGGAATACGTGAGTCCTTTGGGGAGGCAACTTTACACATGAAACAGTCACAGGA
CATCTCAGAGCCATGCAGTGTAGAGGTGTGCTAAAGGAACACACGTCAGAAAGAGAAGT
CAGTCAGCTCGAGAACTGGGACGAAAGCACCAATATTCCTCATTACCAAGCAAATGAAA
ACATAATACTTCACATTCAGATTAGAAGACTACCTTGCCTACAAGTGCCAAGTGTTGGGC
TTGGTGACTTCACCATACGTAAAATACATAGTCCCACCATTTACCACTAAAAATAAAGTTT
CTTCCATTTCTTTTTTTTATGCCTTCCCATCCTCTGGATTCGAATTGGGGGATACAAGTTAT
TATCCTTCCTGGTTTGTTCATAGTTATCTGTAAGAAACACATCTTCATTCATTCTTCTTATA
GTCAAAAATCCTGAAACAGATATGTAATTTGCAAGGAAATAAAAGATCTCTGACTCAAAA
ATTGGGATTGGAAATTCTTTTTGCAAATGTTTATCTGCAGAGAGATTTTGTTTGGCTTCCG
TGGGGTGGTGCATATCCACCCCTAGCCCAAGATGGTTTAAATGTTCTTTTAAAATTTTTCT
TCAATTTTTACCCATGTTTTTTAAAAAATCAATCTCTACTGCAGAATTGCGATTTAAAGCC
ATCTCTATAAACATACACACATTTACATTGGTTTACCCATGAACATTAGGAAAAAAGCATA
AAAGCCCCTCTTTCAAAAAAAAAAAAGACAACCAAAAATCGAAAGAAAAGAGCAATCACA
AAGACACCAGGAGCAGGTCAGTGCACCAGTGTGCTTCTTTTTTTTTTTTTTTTTCTGAAAG
CAAAGATGCTCCCTGCGCTTGTCCCTTTGACCTCTGCAGCACTGGGCCACAGAATGGCC
ACGTCTAGCTGCTCCACTGTTCCATGACAGCTGTTAGCCTGGGAGGCTGTGCTGTGTGAA
TGTCTCTGAATGCTGACAGCGCAAGGGGTCAGCAGCAGGGACCTCAGAGCATGCTGGA

CACCCCAGGGCCTCTGGACTTTTCCCCCGAGGGTCAAACATGCAACTCTCAGTACTGAC
AGGAGCAGGACAGCCAGTGAATTGAATTGCTGGGGGGCAGAGGGCGCTCAGTGCCTTT
GGGGTAGCTTGGGGAGAAGCAGCTTCGAGAGGGAGTGTCCGAGGGGCCCAAGCCGCC
CCGCCTGTGGTTTCTAAGCCGACCCTAAGGTCTAAGCTCGCATGGTGCTGCTTCAGGCTG
GAGGTGTTCCCTGAGTCGTCGTCATCTTGTCCTCTTTACTGCGGGGAACAGATGAGAAGC
AGTAAAAGGAAGAAAAATAACCTTCCTAAAAGATTACTGACTCAATTTTACCAATGAGCA
GTCCACTGCTTGCTTTTTTCAAGTTATAGAAACATGTCATGTTGTCAGCAGAACGATGCA
GCCAGTAGTTTATGTTCATGTCATAAAGTGTTTGAAGAGTGAAAAAAAAAAAAAGAAGGT
GGAAAGCCAGCCTCTGCCGGCAGTTTAGTCGACAGAACCACAGGCAGCTGCACTGTTAC
TGGCACATGTCAGTGGGCACTGGGCTCAGTGGAGTCCATGGTTTGGAATCCTGGTGCCA
CCCCTATGAAGGCAACATCTTAATCATGTTGGGTCTGTTTTCTTCATTTGTAAAACAAGGA
CTCATGTACCTGTGACTCCTGGGAGGGAAGTGAGAGGGGAGAGTCCCTGGTCAAAGCA
CCTGGCCCCTTAGTCAGACTTGAGTCTGAGTGAGACTAGAGGGGTTGCACAGGAGGAAT
CCCGTCCCTTTCTCTAAGCACTCAGAGGTTGCCGGCCTGCCCTCCTGCTCACATGTTGTG
ATGACCTCACCATGGAGCGGGGAAGCTGGTGCTGACCGCAGCTGACCAGCCCGCCTGG
ACACAGGGATGCGTCTGCGGCTTGAAAGGAAGGCAGCTGGCCTTGTGTCTAGCAGGGAT
CCTGTGCGGGGCACATACAGCTGAGAATTCCCTTCAGGGAACCCAGCATGCTCTCAGAA
CAGTCTCCACCCCTGTTCTTTTTCATTTAGAAAAGGGCCTCGCCTGACCCTCTGTGATGGT
TTTGTAAGCGTGGTTGGATTTGTGTGCTCTGAGTGTGTGTGGGTTTTCATTTAACGCTTGT
TTTAAAATGAGCCCTCTTCTCACTTCCTACTCAACTGAAATCCTCCTTTTAAGTTTCATTTT
AAAGTGGGTGAAGGGATGTAGCAGTGTGAAGACAGACCTTGTCCCGGGGGCGGGCGTG
AAGCTTTCCCTTTCCCTGGCTACCCTGTAATCCCAAACTCATTCCTGGGGTCGTTGACAA
GGAAAAACTGACTCAAAAATCACTAACAACTACCAAGTTAACTCAGGAGAAGCCCCAGC
CAAGTTTCTGCCATAGAAGAATCTTAATTTTGGCTCAGGACATTCTTGCAATAGGGCCGC
ACTCCTGTTTCTAGAATATATTATCCCTTCCATGAGGAGAGCTGCTAATTAACAGACCAGT
TCCTGCACGTAGGCACTTGACATTAGATGGCTCCCCTGCCTGAAAAAATGAGTGAGTCG
GGAGTGCTATCCTGAGCTGGCTCTTTACTGCCCCAGGCCTGAATCTAGTGAAGCATGAG
CCTGAAGCAGCGCTTGCCATGTCTCAGAGAAGGGCTCAGTAAACCTCCTCTGAATACAG
CCAGAGAGTAAACCCATCATATGGTCCCTGTCCCAATTATTCAACTCTGCTGGGATAGTG
TGAACGCAGCCACAGACTGTAAGCAAATACATATAGTCATGCTCCAATAAAACTTTATTT
GCAAAAACAGGTGGCAGGCTAGGCGTGACCCACAGGTTGTGCTTTGCCAGCCCCTGTTT
TAGGGAAAGGCCTAGCAGGAGGTCAGGAAGTATGTGATGAGGAGCCCTTTCATCCTATA
CTTTCTAGAAGACAGACCTGGTTGGAGTGATTCATTTTCCCCCAAGGCCAAACTCCAAAA
GGATTTTCTTAGTGCTAAGCTCTTCCTATTTCATCATCAGCTAAATAAAGGAATTGAAGTA
AGATACAAGGTCCTGAAACCAGAGAGGGAAATGCAGGGGCTGGGGATTGTTATTTGTGT
GTGTTTGTGTGCAGGGCAGTGTTGCAGCTCCGTGCTGGGCACAATTTACTTCTAGATTGG
GATACTCACTCCCTACCCTCACCACTCCCACCCAGGGCCCCTGCCTTCCCATACTCTACC
AAATGGTTGTCCACTCCTCCTGGACCATTCTGAGTGACTAGGAGTGTGTGAAGAAGCAGT
CTACTCTTGTTATTGACAACCCTATCACATAATTGTTCCTTATGTTAAGCTGAAACCAGTC
CCTTGATAATTTTGTTTGACTTAACACAGATGCAACGTATTGCTTCTCTACCTTACAGCCC
TGCAGATATTTTAAGACAATAATTGTTTCTCATCACGTTTCTCACCCCAGTGCTTTCCTGC
AGATTCCCCGCCTCCACAGCCCTCCTGGGAATTCCACGTGTGTGGCCTCTCCACACAACT
ACATCCTCTGCAAAGGCCTCCCCAGCTAGATCTGCTTCTTGGCTTCTGTAGTCCCCACGG
CTTTCTCTCATGTGTGAGCACTCACTCACACTTGAGTGCTGAAAGCATTCTATTATTTCAC
AGCACATTTTGTCCTTTAAAGAGATAACATGTCATCTGAGAACAGCACAAACATTTTGTTT
TCTTTAGCTTATTATTTTGAAAAATTAGCAAGCAATAAAAAGATAAATAAAAATTGTTGAA
TTACTGGAATGCATGCATGAACATCATATATAAGTTTTCAACAGATGTGATTCCCTTTTCT

GTACAGTGTAATAATGAATATGGCAGTTGAGTAATGAAAATGGAATCTGTATTCCATCTA
TTAATAGTAGGCCTGACGGTAAATTGCTCTCAAGTTTGGATGAGGAAGTATTTTTCCTTGC
TTTGGTCTCCCCTAGGTTTTAGTTTCCTATCACATCCTACTAGCTTAATAGAATTATATTTG
CTATTGACTACAGATATAGAATCTTGTAGAAAACCCATTTTCTGATTGAATATATTTGAGA
TCTAACTAGAGTGATATTTTCCAAAAACATGGAGTCTTCTAGAATACTAGACAAGAAAAG
ATATCATTGCCATGCTTTGCTTCATTATTAATTACTATGCAAGGGAATACACACAAATAAC
TTGCAGGGGTGAGTCACAGAACACACGTCGTATATTCTTTAAAAAATTCCTACTTTCAACT
TTTCTCTTAGGTTTTCTTTTGCTTTTTGAAAATGTTTTGTGATTCTCTTCTTTCATTAAACTTT
TTAAAATAGCTTTCATATTTAAGGGCAAACTTTCTTAAATAGCTTTCATATTTAAGGGCAA
TGACTAAGGTGAAGCTGCTCCGCCATGACAGTAATTTTAGACCAAAGAATATTTCAGTGG
ATTTCTTTAACTCTCAGTAATAAACTATTTGAATTTCTCCTCCTTGGGGACAATTAGAAAT
GAACCACAAAGGGACACATCAGCTGCTTTGATCAGCAGAAGTGTCTTTCAAATGCAGGC
ATCGTTCGGTGGTCTAGGAGACTCGTGGTAATGAAGCAAGAAGAGGAGTGGAGCAGCA
GGGCTGGGGCCAAGTCCCTGGGTCTAGAAAGCCACTATGAATGAACGCTGCTTGCACCC
ATTATAGGTCTAAGCACCAATCCAGGCTCAACCGCTCACTCCAGGAGGATGGGAAGCCT
TTGTTTTGGTTCCTGTGCCAACTCTGGCTTCTCTGGCTGCCCCATTCAATCTGACAGAACA
TGTTGACGGAGATAAGGTTATTACTGGCTCCTGCAAGGCCCTACCATGTGAGGTGGTTTC
CAGTGGCCTTTCCCGGGGCTGAGAGCGGGGCCTGGTGAAATGGACATATTATAAGAAGT
CTGCAATCTGTAATCTGAAGCAAGCTTTTGTTTTTCTTTTTAAGTTGACTGCCTGGTTATTG
AGGCCAGTAACCAGAGAAGGGAAAAACTGGAAGTGTGTACTTTGTGTCTCAACTGAATT
CATCCTGAAGTTGTTTATTCTGGCCCACTGTCAAGTCTTTATAAGGGGGGTGTTGGCGGC
AGAAAGGATTGGTCCTAATGGCTATATGAGTGCCTTCTACTCAGCTAATTTCTATGCTGTT
GAACAGCCATTCAAAGCGTATTATTTTTCCTTGAAGCCGAGAAAGTTTCTATACAAGCATT
CCTTTGAAAGTGCAATTAACTTTTGTAACAATTTGTGCTTTAGTTTAAGAATTGCAGATGTT
GTGTGGGTGGTGGGGGGTGAGGGGGTGGGAAGGAGAGGATGAAGGAGTAAGGATATA
GATGCCTCCCACTTGGAAAATGTGGTAACACTTCCTTTCGTGGAAAAAGCGCATCAATTT
TTAGAATTACTAAAACATTTAGATGGTTGGTGAAACAACTGAGAAGGCCGTGATGTTTGG
TTTGGGGGAGGTTGGACTGGATCATCTCCCAGCTTCCTCTCCTTCCTAATGAGACCCAGA
TCTTTGTGACAGAATTATCTGGAATTGTGATTTTATGCTTGGAAACTGTCCAAAGGTTGGG
TAGCAGAGCCCTGCCACTCATCAAAAATGCTAGCTCAATTTGCAAAAGAAATACAAAAC
CCAAAATGTCACATCACCATTGGGATGTAAGTATTGATGTGCGTGGGTCAGAGGAGCTCT
TATTAACTAGATGGTACATTAGTTTTCACTGGGGAATGGTTTTCACCATTAGAAATACACT
TGCACTGCCCCCAAGTCTATTCCTTATTATTTAGTGTTAGTTTGAGGTTAAGCGGGGGTGT
TTGGTTTCACTGAATGTTTGTGGCTATTTCATACTACGGTATGATCCTTAGTAAACCTTCTA
CCTCAAAATCCACACTACTTTTTGTATAATGAATATCTCACTCATCCCCTGACAAACAAGA
TAACATTTATAAAATACTTTATAGTTTGCATTTGCTAATAATCTAGATTTTCTAATTCATGT
ATATCCAGTACTTAAACTATTAACAGAAATCAACCTAATAAGGAAACTATTGAGATTCAAT
CCCACTAGAATTAAATCCCTTTTATATGTGTGACTTCTAGGAAAGCAGTTAATCAGAGGC
AGTGCAGGAGCATTTCTCTCCCCCTCACTGGTATGTGGGGATGAGGTGTGCAGTGGGGG
CACCTGGCATGGTCTTTGAATGCATGGGTCCTGGCTTCACAAGCCTGCGTTTGCACCCAG
CTCTGCCAACAGTAGTGGGGTCCCCTTGGGCAAATGACTTCTCTTTACTAGCTTCACTGT
CCTTGTCTGTTAAGTGGGAGTAGAAGAATGCCTCCCTCGTGGATTGTTGCAAGGATTAAA
TTGTTAATATTTGCACAACACCCAGAAGAGCATCTAGGTCATGTAAGTGGTTGCTGTTGT
CATCGTCCTTTTAGGATTAGTATGTGCAAAAACGAACAGTCCTGGGACATAGCATTGACT
GGAACCAATGCTGTTCCTGCCCTCAGGAAGCTTACCATCCAGTGGGGAGACAGACATTC
ACTGAATCATCACAACATCACAACTGTGATTATGCCATGGGGATGATCTAAGAGTGCAGA
AACCTGACCTGGTCTGAGAGGTTCTGGAGGGGTTCCTTCAGGAAGCAGTTCAGCTGGGC

TTCTAAGGACAAGCGGGGCCCCGCCACATGTGGGGAAGAGTTGGGGGCAGGGGACCTT
CCTGGCAGGGCAGGAGGAGAGGGACTGTGAAAGGCTGCCTGGCTGAACAGAGCCTAGC
AGAGATGAAGCTGAGAGTGTGAGGGGAAGTGGGATGGAGAGGTTCCTGCTGCAAGGCT
TGTAAGGACTTTAGATTTCATTTGAAAGGCACTGAGAAGCCACTGACAGATTCAAATGAA
GAGGAACATGTGCAGATGTGTGTTTTTAAGAATGCCTCTGCTTTTTGGAGGGGGTCCTAG
ATGGCATCTGGAAGTGGGCGTGTCTGTGACAGCACAGTGCAAGGGGCCCTGTAGCGCA
CAGGTGCTCTGGACTTCGCCTGGGTCAATGCCTCATCCTGGCTGTGACGCTGCAAGACG
TCTCCGTGGGGAAAGCTGGTACGCTGGATTTATCCGCATCATTTCTTACAGCTAGTATGT
GTGAATCTACAATTTTCTCAAAACAGACAGTTTAATTACATAACTTCTTGAGATGTACAAG
CTGTAGGGCAGAGCCCTTGCAAAGAAATATCATCCCTAGGTATGTGTTTGCTTTATGGGA
GGAGCTGGCACCCCAAAGCTGGAGCACCCAGTGCTGTTGGAAATGAGCCTGGCTTCTGG
CAGTGCCCTGGTGACAGGTGACTCTTCTTCTCTAACTGAGATAAATGGAGTTAGCCATAA
TAGCTTGTAACTTTCAAATACTATGATTTTTTCTTATAATTTTTTAAAGCATGTGGTTTCAAT
GTTGCATACTCAACTTATGACTAATAATTAGTTTTCTCAATCCCATGTTGCACATTTTAGTA
TTAAATAATCTCCTTCTCCCAACTTGAAACACAAACACAAAAACAGATCTTACTGCCATGT
GGAGAAGGGCCCATTAGGCCTCCAGGGAGTGTGGAGAAACCCCTAGGAGGCGACAGCA
GAGCCTGGAGTCCCACCACTGAGAGTGTCATCTGCAGCACAGCAGCGGGGACATCCCC
AGGAGCCTGCTGGAAATGCAGCCCGAGGGCATCCCAGATTTGCTGAATCAACATCTGCA
CTTTATTAAGGTTCCCAGGTGACGCTCATGTATGCTGAACTTTGCAAAACAGTGCACCAG
GGAGATGGCAGGGTGAAGGGGCACAATGGACACCCCGACAGAGGTTAAAGAGGGTGA
ATCGACAAACTTGGGGATGGCTTAGCTACAGAAAGATAGAAGAGAGTGGCATCAAGTAG
CTGTTGACATCTCCAGGTCAGAATCGGGAATGTTCGAGTGAGTATGGTAGTAGGAGCAC
TGGAGGCTCAGGTTTATAGACTGTGAATTTGGTGTGGAGCAAGGCATGTGGGGCAGCCA
GCCTGAGATAGTGTCTGGTCTTTCAGCTAGGGCTGTAGAGGAAATCAGGGCACCCAATG
GATGATCAGAAGGGAACGGATCCTGACTGTTGTCATCTTGGCTGGAATCAGCCAGTTGA
ATTGCCCCCGAGGGATCCATCCTTTGGATCAGTTATGCCCCATCTAAACTTGCAGATGAT
GATAATGCAGATTTCATGAAAATCCTAAGATCCACAACTCTTAGCATAAAGCCCTCTATA
GTTTTTCAACCATAAGTTGCAATCAAAGAATATCATACACAGCACCATGATTTTAGAGCTT
AAATTTGTAATTTTGAAAATCTGAAAATTTAAGAACTGAGTTAAGCAAGTCTCCAAGCCTT
GGCCCAGGCTTATTCATTTTTTATTTTGAATAATTTAGTCAATTTTATGGCCAAAATACCTC
TCCAAAAACCTGGTAGAAATTTGAAATATACTCATATCAATAAAACATGTTGGAACAAAG
GTTTTAGATTTGATTGAAGTAGTATGGAATGTCATAGGGACATTTTCTGACAATAGCACTG
GGAAACATATCAAGTCTGAACATTTTAAAACCATGACTGATAGGGAGAAAAGTTTTTCCTA
TAATTTAAGCTATAGCTTTCTTGTTAATTTCCCTATTTCTTTATGCACTTTCTCTAACTAGACT
CTAGAACTACACGCATACATTAATACACACACATACACCAAGCACACATGTACATTCACAC
TTTTATGTGCATACACACATACTTGCAAACAGATGCATAAAACACACATACATGTACACAAT
ACATGCCCACACACGTATACACACTTACTTGCACACACATGCACACACACACACATCCATT
TACACACGTGCATGCATGCACATACGGGCACATTCACATGCACACACACACAGAGATTTA
ATTCTAGTTTGAGTTAAGAAAAACCTGCTGCTCTAGTCTTTTATTACTCTCCCCTCCTGCTT
GATTCCTCTAGGTTTCTTCTCTTCTAGGCTTTCCCCCTCATAAACTGCCACTGGGATTTGAA
TTGACATTCATATCCATGAGATATTGTAGCTCTAGACCTCCTCTTGCAAAGCCTCATGCAAT
AAGGCATATGATAAATGTGAGATGGAGACCCTCAAGCTGCTGCCAGCAATTGCTTATGTGC
ACCAGTAGACTCTGAAGATATTTCACAAGGAGGTACACTTAGCTATGAAGAAATGCATTTC
CTACAGATGCTACACCAAAAAGGAAGACCAGAAAGGAGAAAAATATATCCTTTCTGGGAT
ATATAATCTGAAAGAGATATGCAGTATTTTTAGTTAAGTTTTCTTTATATTTGTTTGTAAAAGG
AATGATATATCAATTACTCTAAGCAAATAAAATTTGCAAATTAATTTCTTAATTGATTGCACA
GATATTTGTTTCAAAGAATGACTCTTTAATCACCTGGCTTTGACCGCAATATCACCACAATA

**CCTTGAGTTAATCTGAAACGTACTATGTTGTACCAGGAAAGGTCTGTAATCACATATCACAT
TTTCTTCCCCCAGACGTGATATTTAATACACAACCTGATAAAATAAAGGTAATTAGGTAGAT
GATTTTCAATATCTTCTATAGCAAGCCTGTAGACAAAAAAAGAATTCCTGACCTAGTCTTGT
TGATTTTTTCTTACTACAAACTAGCCTGGTCATCCTTCTAAAATATCCCACCATGTGAATTAT
TTTGATGAGATCTTTGCATTGAAGGTTGCACTGTGCTAGTGTGAACCATTAATTAATCAATT
CATCAATTAATCAGCTATATACTGAATACAAGACACTGCACGCTGGCCCCAGCACTGAGG
GAACTCAGTTGAATGGCAGAAAAGCAAGTCAGCCACTACCCCACATTGTGACAGATGCT
GTTGTGGCAATACTCATGAGACATGCTGGAAAAGAAGTTGAGGCAGCTGCTGATTTTCAG
CTAAGAGAGGATGAGGGACACCTCATCAGACTGAACAACTAAGGAAAGTGTCCCAGAGG
ATCTGATGCTTGAGCTGTCCTGTGATGGATGGCTTGGAGGTAGGCAGAGGAGAAAAGGG
GAGAAAGGGCAAAGGGAACAGAGAGTGAGAAGGCACAGAGAGGGGAGGCAGCAGGA
GGAGTCAGCACATCTGTGAGAGAGTGAAGATCCCATGCAGGTTGGAACGGCTGCTGCCA
AAAGTGAGCGGAGCTCAGAAGGGCAGGAATGCGCAGCCATGGCTGCCTCTTGAGGGCG
AGGGTAGGAGGCTGTGCATTTCTTCAGCAGCACACAGCGTGGCTGTGCGGTTCCCCCCA
CTGCCACCAGCAGCCCAATGAGGACACATGCTGCACTGGCCCAGGCATGGACTTTGCAG
GTTGGGCAGTAAGAGGGGAGTGAGGGAACACTAACGATGTGAGGCTGTGTCATGCAGAC
TGGACAAAAAGAGGGAGTGAGACCAGATGGGAGCTGCCAGGTGAAAAATCACGTTACC
AAAGGCCAAGGTCGACGGAGGACTAGAGTGCTCCATGAGCAAGGCTGAGAGTGCTGGG
AAGACAGCAGGCCAGAGAGAAATGTGGCCCGAAGACTGGAGAGGTAGAGCCTTCCCAG
TTATGGTGGGGTCACAGAGGCTCCTAGAGAGCAGATCTCTTCAGTGAATTGACTTGTTGC
ACAAAGATTATCTAAATAAATTGTTATCATTTCCTTCCAGAAGTGGGCTCAGGTCTCCCCTC
CAAGTCTTTTTGGATAGTGAAGAATCAGTAAAGAGCCCCTAATGCGGATTAAAGAGAGAT
GCCCAGATCCACTGGACTGAACTGGGCAGCAGGGAACTGCCTTTGAAGAACAGGCCCC
ACATGGGACAGCTTCATTCCCAGTCAAGGTCACTGTCGCACAGAGGCCATATATATTTGTT
AAATGACTGAATTACTTATGGTACAAACTTGACTGCATCACTTAAATGCTTTTCGATTCTAG
TAAGAGGAAGTTCTTCTAATATTAACTTCGATTTAGTGGAAGTTACCCAAGAAAGCTCAGT
AATCCAAAATAGAAGACAAAGTCACCATTATGTGATAGAGTGTTGTATGTCTAGGGGTCTG
TTTTTTATGGTAATTCTCTGAACCAGGAGTTCCAATGGAGAGAACAAAACTATAGTGGTGG
TGGCTGGGCAGAAGGCTGACAGGGAGACAGAGCAGAAGACAGCACAGTCACGCTGCTG
GCTACTAGGACAAATGACAGCTCTCGAAGGGCAGTACCTCACATTCCTCTCATCACAGAA
AATGAGAAGGTGTGTAGGACATTCTTCTAATGGCTGAGCTGTTACCTAAGAAGGGTAACA
CACAATGCCACTGAAACGCCATATGATATCTGCCAGAGGCACGGGGCTCCCATTGACTCC
TCCTCTGTTGAAATGGATTCTGAAAGAGCAGCAGCAATGAGAGAAGAGAGCCGCCTTTTC
CTGGGAATGTCTGCCAAAAAATGTTTACCTTCTCTTCCATATCAGAGTTTTATCCCTCAATC
TATGTATGGTTGTAAGCAGCTAAGCATTCAGAAGTTTGGATTGTATGGGCTTTCTGAAGAG
GACAGTGTGAGGAGCTCTTGTTTTCATGCTAAATTGGAGACTTCATCCCTGCACAGTATGA
CACACTTCCCTGTGTTGTAAGACTTGGATTTAGTTCATGGAATCTTCCAGGCTTTCAAAAC
AGATGACCTCTCACTAGAAGATCCACTTATCTCTTTCCCCTTGGCCCTGTAAACCAGCCCA
GCAAAGTCCCAACTTGCAACAAACAGGCAATTCACTCTACCGGAAAACAGTTATCTAATC
TCTCAAAAATGACACCTGGCCAGTTTCTAACCGTAATGCCTTTGGACTCATTTCCATCAAA
TGTCTTCCTATAAGCAATGTTAACAATGTCTTTATAACTGTAGGTGCTGTAACCAGTGGAGA
ATACAATTAAAAATAGGAATATGCAGCCCGATGAATGCCCTCCACTCCAAACCAGTGTTAT
GAGATGTGTCAACAACGTTCCCAGGAGAGCTAATCTGTTCACCATTCAGCAAGTGCTCAC
TGAGCCCCGGATGTGTGCCAGGCCCTGTGTGTGCTGTGGAGATACCAACAGTCTCCGCC
CGGATGGAGCTGACAGTCTAGGAAGGGGGATAGATGTTAAATACTCTGGTGAATATTTCTC
CCACCACCACCCATTGGCTGATTTAAAAGGAAAATGGGGGATGCTTTGCAGATGGTTCAT
TTCCATGTAAACGTCCTTGTGGTTGTAATACATTGCTTTCTCTACTAACAAGAGTTAACTTT**

CAAATGAAAAAATGGAGTGATTATTTGTCAAACCAAAGTGCTTTCTTCATTATTAATAAAGT
TGCTCAACATCCCTTTAGTAAGTGAAGTTATTCACACAGGGTACCAGGAAGTGGAGATAGT
CACATATTAGCCCTCTTGGGCCCTCCACGTGTAGGCAGATGAACAGGAACCACGAGGGT
GGGTGGCGTCCCCCAACACTGGAGAGCAGGCTGTGTAGAGTGCTCTTGGTCATCCTGTC
TGCAGTCCTCCTGCCAGGGAGGGCAGCACCTGATCACAGAGAGACACAGCGATGCAAC
AGAAACACTGAAAAAGAGTAGCTTGTCCTGGCCCACGTCAGCGGCCTCGGGTCCATCCA
GGTTTGCAAAAAACAGGAAGGTTTTCACTTGTATCTGGCTACCCAGTCAGCATACTGGCC
TCAGCACTATTTGTATGTCAGGAACTCATGAGACCAGTACTTCTCAAACTCTCATGGACAT
AGGAGTTACCCAGGACTTTGCAAAAATGCAGATTCTGAGATTTTGGAGGACCTGGCCAAC
TGCATCTCTAACACACTCCAGGGATGGTTGTGCCCATGGCCCTGGGGAGGTGAAACAGAT
TTGGGGCAGTGGTCACCTGCTGCAACAGAATGTGTCCTCCAATAGAAGGACCTTTCAAGA
CACTGATAAGGATGGGTCAAGAGGAGAGGTGAACCAGGGCAGGGTGGGTTCACGCAGC
CCAGGAAGACAAAAGCATAGGACCGTATTGTCAAATGCAGAGGGGCCAGGCCACGCCA
GAGCGAGGGGAATCCCACAGATGTAGCTACAAAGGGATCCTTGGGATACCTTTTTTTGAA
CATTTCCTAGATCAGTTTCAGTAGAGTGGCTAAACAGAGATATTACATTCTGAACAAATTCA
GGCTTTAAACAGGAGAAGATACACAGTGAGACTCTCTTCCCACAGCATTTGCTAGGGTAA
CACCAACTCCAATTATGCTGAAATTCTAGTTTCCAAATAGTTGATGGTTGTTGGGGATGTAC
AGCATCAGAGCCAACATTGTCCCCTATATGTTTCTGGAACCACTCGGAAGATTGTTTTGGT
TTCAGGCTCTATGTCATGGCTTATTGTCTTAAACCAGGAGAGTCTGACATCCAGTTCCAAG
GCTGTAATCTCTCTCTGTTACACCTCATGGGAAAAGGGTTTGTAGGTCATGATAGACACTG
TGTATGGAGAAATAGGTACTCAATTCATTAGGAGGGCACTCTCTCAAAAGGCCTGTAGTTA
TTGATCAAAAATTAATTTTATAGCTTTCACATTCTTCTCAGATCAGATGTATTAGAAAATTAA
GGTCTTGATGATGCCTTCCTCTTCTTGCCAGGATAAATTGAATGGCACTTGAACTGTAAAA
TGAGCAGGGGAAAGAGCAGAGTTTTAGGCTTACTTGGTCCATTGTTCACTTCTATTCTTTT
CTCCTTGACTTTGGAGGAAAGGTAGGATTTGTTAATAAAAATGTTTTTAAAATTTATTTTTAA
AGTTAAACTAGCTTTCTAGTATGAGTTTTTTAAGTAAAATAACCAATTTTCCCACGAGTTTG
CTACTGAGAAAACACTGAATGGGAAGATTTGGGGGAGCATTTCAGGCTGATGGGCTGACT
TCTTATGTGTCACTTCACCCCTCCCTGTGCAGTCCTGGTTCCCGCCCGCCCTGGGACACT
TCTCTGTGGATCTGTGAGATCACAGCTCCTTTGTCGGCCCCTGGAGCTGTCACCCATGGC
TCTGAATGACTTAGTGACTTGTCATGTGCCAGTGGACATGACATTTCTGGAGGGCAGAAA
TGGTGTCTATCCAGCTAGCCACAGCCAGTGCCCAGTGGTGGCCTGACACATAGTGGACA
GTTGACACGTGTTTGTTGAATGACTACCCCAAAGGTGGCAACCTTCCACACAGCCGGGG
AGTGGGGGAGAAATGTGTTCAGAGGCTGGTCCACCTGCTTCTACCACTCTGGTCTTTGGG
ATGGTGCCTTCAGGTACCTACTGAGCCTATCTTACTCTAGTAATCTTTTAAAGTATTTGAGAT
TACCAGTGCTTCTCAAACTCCAATGGACATAGGAATTACCCAGGACTTTGCTAAAATGCAG
ATTCTGAGATTCTAGGAGGGCCTGACCATCTGCATCTCCAACATGCCTCAGGTGATGGTTG
TGCCCGTGGTCCTGGAGAGGTGACATAGATTCAGAGCAGTGGTCACCTATTGCAACAGAC
TGTGTCCTCCAACAGGAGGTCCCCATTAATTTCAGAAGCACTGGCAAAATGCTCAAAAGG
GTAATCTCCATCTAAGCATTTTATGAATAAAGGATTCATTAATGAAAAATTAACTTGTGCTGG
GCACTTTAAAAACCACTAAAATGTCAAAGTGTAATTGTAACGCAATTCTAATTTATAGACAC
GCCGAGTCTGAGTGGTGTCAGGCCCTACTCTGGTCTGGATTCCCTTCCTTCTCCTTTTCTG
GGGGCCCAGTTCTGCACAGGTGCATGTCTCAAACTAACTCCAGTCTTCCCAGCATCAGCA
CTGAAGTTCTAATTGCAAATTTTCAGTGTTTTGGCCTAGGAAATCTAGCCTTTTCAAACTTC
TGAAGCCTCAAATATACACTTTACCAATGCTTTTCCGCAGCCCTGACTGTGTCATTCACCAT
GACTGCAGAATCACATACACATATGCAAACCCTCGGGAGACTTGGTCTTCCTTGCTCCTTC
GTTGAGGGCACTCTGCACAGGGTCTATGGCGCAGAAGGGAATACCGCCTAGGACTTGAT
CGCAGCAGTATGCCCTGGTGGCTGCTTTGGGATTATTTTATATAACAGAGATCTATTAAAGG

**GCCATGTTCCCTGGAGGAGTTCTCCCATCATTCCAGGTGGTTTCCTCCAGGGTTGCCCCA
GCTCTGCTAGTATCAAATATCCTCCTAGGGCAGCAACCATGGCCAAGAGCCGTTCCTCTTT
GACAAGCTGCCTCGTTTTCTGCAGCTTACCTATAACATGCTAATTGAATGTGGGAGTTACA
GAGTGTGGGTCCCCCCCACCCCGGGAAATGCCCCTCCTTGTAGACTGCTATGTCTTTCAT
TGGGTACCATTATTTCTCATCTTTTCTTGCCTGCATTCAACAAACTTTTACCGATCTAAAAA
GGTGGCAAGACAACTTTTACCCACATCTCTATTCCACTTTATAAACTATTTGAATAAAAAGA
TTAAATATATTTCCCTTCAAATTTGCTGAATAGTTAAGTAATTGATATGGAAGATGTATGTGG
CAATCTTTGGGGGTTTACTGTGAGCAATTAGGAGATGTTTTCTACTGCTGGTGGTGTATAAT
TTGCTATAACTGCGCTGGAAAATGTTTTGGCAATATCTTTCAAAGCTACTAAAGCGACCAG
TACCCTCTGACCCAGCAACGCCACTCTGAGGTACCCAACAGAAATGCAAACAAAAGTTC
ACCAGGGACACGTGGGACAATATTCATAAACTTGTAACTGCAAGAAAAGAGAAAGAACC
CATATGTCCATTGGCGGTAGAATGGACGAACGCAGTGTGGTATTCTCCTATGATGGAACTG
CCCAGTTATGGAATGCAAGAAATACAATTTTGCAGCACAGCATGGAAAAATATCAACACAT
AATGTTGAGTGAAAGAACCTGACCCAAAGGAGTACCTACTGGAGGATTCTATGTAAGCAA
AATTCAACAGCAGGCAAAGCTGGCCCATCGTGCTGGCCTTAGGATGGTGGTAGCCCTGTC
CCAGGAGGCAGCACGGGGTGCTTCTAGGGTGCTGGTATGGTCAGCTGTCCTGATCTGGG
TGCCACTTACAAAGGGCTAATTGGTTTGTGATATTGGCCAAATTACACACCTTTGACTTGT
GTGTGTGTGTGTGTGTGTGTATGTGTTTTTTTTGAATTTAGTTATACTTGAAAGTTTACTTGA
AACATTTCTAGAAGGCTGCTCCCTCTGAGCAAAGCTGCCTGGGCAGCATCGAATTCCCAG
TCAGGAACTCATGTTTTTAGAAGGGAGTTGTGGTGAGGCTCTTAGGAGTACATAAATGAG
AGGTGCAAAAAGTGGGGTGCTTGGTGCAACTGGATTTCTTGTGGACAGCGGCCAGGGGT
GGGATGTGCGCATTGCGTGGGCACAGGAATCTCAATAGCTGGCCCCAGAGCAGAAGCAG
AGAAGGTGTGAAGGCCTGGACTCTTCTGAGCTTGAGGTCAGGGGATCTACTTGCACATTA
GAGTTATTTAAATGAGGAAAATGCATGCCCTTTTTGAATCGATGTTGTCAAGCCTTTCTTTT
CCTAGGAGAATCTTTCGATTCCCAGCATGGGGAAAGCATGTTTTCAGGTATTAAAATGAGT
AAGGTTGTAACAGTTACCATGAATCCACATGTTGTGCCCCATTCTAAATCAAGTATGATGAT
AACAAACACTGTAACCCGTGCCTAGGAACACTTGTTTCAAACTTCAGACCCCTCTCCAGT
TACATGGGAAGCATGGCTGTGGGCATGCATCTCTCTCCGCATAATCTGCGGTCGCTTAAAA
CTTAAAGTTATGGCAAACCATGTTGTCCTGATGATTATAGTTTCCTGATGTCACTCCCCATT
CCAGTTCTGCAATCCCAGGGACATCCAGTCCAACTACCCAGCAAAGACACCTGCCTGCTA
CTGTATGACGGCATTGGCTGAGTTTGAGGAGGGCACACAGGAAGCTCCAGGTGCTATCTA
CAGCCTCTGCATAGGTAAGTCACTGGGTCCTGAGAGCAGACCGGGATATTTGCAGGGAG
CTCATGGATGGGAATGTGAAGCTGAAATTTCACTATACCTTCAGGAAGGCTTCCAGGAAC
CCCTTGAGATTCAGAGAACTGCGAGTTACTCTCCCTTACTGCCATTGATTATAAAATCTATT
TTTAAGAGAAGGAATAGCTAAAATCTTAGATGATAATAAATATAACTTCTTAAATCCAGTAG
AGGACAATAAATATATGTTAAAACGAACTCATGAAAATAGCATCCATTGACTGAATATTTAG
TTTGTGTAGGCAATGTGCTTGGCGTTTTTACAATAATCCTGGCAGGTAAAATGAAATGTAAA
TAAACCTTATTCACAGGACTATAGTGAGGTTAAGAGATCATGCCTGTCTCACTTTTATAAAC
TATAAACTATGTGAGTGCCTGGCTCACTTTTCTGATACTTTTGTCTGAACAGAGTCTTGGTG
ATATTTTAACTTATGGTATTTTTTGAAGCCATACTAAATTCTCATGTGAATTTAAAGAATAATA
TAATGCCTGTAATCCCAGTGCATTGGGAGGCTGAGGCGGGCAGATCATGAGGTCAGGAG
ATCGAGACCATGGTGAAACCCCGTCTCTACTAAAAATACAAAAAATTAGCCCGGCGCGGT
GGCGGGCACCTGTAGTCCCAGCTACTCAGGAGGCTGAGGCAGGAGAATGGTGTGAACCT
GGGAGGCGGAGCTTGCAGTGAGCAGAGATCGCACCATTGCACTCCAGCCTGGGTGACA
GGGCGAGACTCCATCTCAAAAAAAAAAAAAAAAAAAAAAGAATAATACAAATACAAGTTGCAG
TGTTTCATTATTTCTACACAATGTCCTTCTCTCAAAAGCTGATATTTATACAGGCGGTAACAA
TATATTTAGAATTTTTTGTAATGCCAGAAAAATAACAAAACATAAACACATTGATTAATACTT**

TAAGTCCATTGGACATGGATACTATAGGAAAGAATGGATTCCCCAGTCATCTCAAAGCCAA
AGGTTCCTTGTTATAAAAAGTTAGAAATACAGATTTTTTTAAAAATAGTATTTTTAAAAAGGT
ACTAAAAGTCCAATATTAAGAAACTTAAAAATTATTGTATACATAGATATAAAATTGAGTCCT
CTAATAATTAAACATCATGTATTTTTAAATGTTTTAATTTATTTATTATTTATTTTAGAAACAGG
GTCTCACTGTGTCACCTAGGCTGGAGTGCAATGGTGCAATTATAGCTCACTGCAGCCTTG
AACTCCTGGGCTCAAGTGATCCTCCAGCCTCAGCTTCCTGAGTAGCTGGGACTACAGGCA
TGTGCCACTACACCTGGCTAATTTTTAAGCCTCAAGTGATCCTCCTGCCTTGCCCTCTAAA
GTGTTGGGATTATAGGTATGAGCTACTGTGGCCAGTGTAAACATCTTTTAAAATATTAAATG
GTATAGGAAATATCTCAGGATATACTGTTAATGAGCAAAAACACACACTAATATATTAATAAT
CATTGTTCCTAAATGACAAAATTCTGTGAGCCTTTTATTTACTATAACATGCTTTTCTATATTT
TTGTAATATTCCAGAGTGCCCATGTCTTACTTTTTACATTGAGGCTGTGAAATGTGAGAACA
CAGGAAGGAACCTGGTTTTGGCACCTCTGTGGAGCAGTGCTTCCCACCCTGGCCACAAG
TTAGAGTCACCTGGCAGCGTCCAAGCCCCTGCCACCATGGCCGTGCCCATCAGAGTGTC
GGGTGAGGCCTGGCTTGGCAGCAGCTGCTCAAAGCCCCCAGCAGGCTCCAGGGCTGCT
GCAGGAAGGCAGGCACCTACTTTCTAGAGGTCTGTGGGTTTTGGATGGGGCTTGAAACC
ATATCAAAAACTCCCAGAGCCCAAACACAGCCCTGCTTCTTATCAAATCACCTTGGAATCT
TTGTCAAAGCTAAACATTCAGGGATGTTTGTATCAGAATTAAAGTTCTGTGCACTCATAGG
GCAAGAGTGTTGAACAAAAACTAGAGTTCTAAGCTATGCTTTTGGATGAAATAGTGAATCT
AATTGAGATTGTCAATGATTTTTTTCCTTATCGATCCTCTATTGAAGGAGGTACATTGACAA
AACATTGGGAATGTGAAGAGAATGTGGTTTACTTGTCTGTGGGGGGAAAACATCACACTT
GTACACATCAAAGATGGAGGAACCAAAACCACCATCCTGCAAACACATGGGAGAGGGGA
ATTGCTGTCAGGTGTTTTCAACTGTATATTTCACCTGGGAAAAAAAAGAAGGAAATTTCCT
GGGTGCCAGGTGATTTGAATAGGACATTTTTGTGGGATTATCGAGGCAATACCATGTTTAC
TGAAAGCGTACTGTGTAGCAGGCACTCACATAGTCTCACACAGTCAACACGACCATGAGG
CAGAGCTGGCAGGCTTGGCAGTGTTGGGTCATTACTAGAATGTTTAGGGAAGGGAGTTGT
TTTCATTCACGGAGCTTCCTATTTAACAGAAGGTTAAGGAACAAAACCTCTTTCCAAGGTG
ACTTTTGAAAAACTTTCTACAACATACAGGAATCGCTGCTCCCCTGGCAGACAGCCTGTC
GGTGCCTGTCATCCAGCGTCTGGTACAGACATCTCCTCCTTTTCATTCTCAAAAGTGTCCT
TGTGTGGAGGGAAAATGATATGGTCACTCTGAGTCTAAGGGGATAATTTAATCTTCCTTGA
TGACTATCTCTGGATCGTCAAGATAAGGTTGATTCCCATTATAAAAGACAAAGGCTGGAAC
CTGGGAATCAGGTGTTGCACTGGCACTTGACTGTAAGGCTGTATGGGATAAAGAAAGGTG
CACAGTCCTGGAAGTTGGGGGTCTGGGATTCCAGTCCCTGCTCTACCAATGGCTACCTGT
GTGACCTTGGCCGGTTACAGAGCCCTCCGACAGCAATTCCATAGTCTACAAACTCACTGC
CTTGGTATCACAGGGCAGAGGGATTCCCATCCAGCGTTTCCTACTTGTGCATTCCCATACT
GATCTCCAGAGGCCGGCAGGAAGACCCTTCCTCGCTTCACAGGAGGCTGCTGAGGCTCG
GCCAGCCACTCCACAGGGCCTTCTGAGGGGGCAGCCATGGCCCTCGGGCCACACTTCCT
GTCTACTGTGAGATCAGCCCCTGGCCACACCAGGCCTGATTCTGAGGCCATGTCTAGGCG
ATGTGAAAACTCTTGATTGAAAGTGACCACCCCAGCTAATCATCTGTTTGTTTTGACTCCA
AATTATGGGTTTGAGCAACGTGCCACCCCGGGCAGCTGCCTCAGTCGCCTGGCTGCTGT
GTGCTCATGGTTGGCCTGAGACTTTTCACCCACCATGCAGGGCATGAACTTCTGGAATTT
GAAATGCTTGGGCAGGGGAATGACCCTGAGCCACAGCACTCATCAAATCACTACTGAAC
CACCCTTAAATCGTGTGGCTATCAAAGTGAGTCACAATGAGACCCGAAGGGGTGAGACG
CCGACTGCAGAACTGCTGTCTAAGCTGGGTATCGGTTTGCAGTGGGGACCTCTGTAGGG
GGACCTTGCTGCCCTGCTTCCGGTGAGTCCTGTGGGTCACTTAGTGAGGGCGTCGTGGG
TGAGGCTGGGTAGCCCAGGTGTCTGCCTCAGTCACGCCAGGAGTTCCAGTCCCAGCTCT
GGCACCCATCAGCCAGAGACCCCAAGCCCATCACGCAATCCCACCCTGGGACCCCACCC
CACATCAGCCATCCGTCAGGTGGTGACTGCTGGTTAAACGAGATGCTGTAAACAGTGCCT

GGCACACAGTGATGTAGAAGCAAAGGAATATGTTTCACCTCCTTCCTGTTTCAAAAAGGT
CTCCATTAATTGGCATCATCATAAGCCCTCCTAAATGTGGTATTTACACGTCATTTATCTGCT
GGGCTGCTGACATAGAAGGACAGGTCTTTACAGAGGAGCTGGCTATCATAACATCCACAC
ACTCCTCAGGTGGGTGCTCGCCTGTGCACCATCTCATCTTAGCTCACTTAACTGAGCCCA
CTCACTCCCGAGCCAATGCCAGCTTGCTACATATCAAGTCCAGGAATTCATTTCTATAGGA
TTTTATTCAATCTAGAAAGTAACATGATTTTTAGTGTTTTTTAGACAGTATTTTGAGCAATTTA
AAAACCCCTAATACAATCACTGCTGCATTTTATTCAGGAAGCCTGGTGGGTTCTGCTTCAA
GCCTGTCTAATCAGTGGCACAAACAGGCTCAGGAAACTGGAATCCAAGTCTAATAAATAA
AAGGTGTGAGGTTTAGAAAAAAAAAAAAACAAAACGTCTAGCAAGCGAGATGAAACCTGC
CTCGCTGGGCAGCTGCTCTGAAACCGGTTACTGTGGATTGCAAATCATACCCTGGCTGAT
TGAGCCCTAAACCACAAAATGTCTAAAGAATGAAGCAAAACACTTTAGTTCTAGTGCACTT
TGGTGTAAGAAGGAATCCATTAGAGTTAGAGGACTCAGCGGGCCAGGCAGAGAGAGCAC
ACACCGCGCGATACCCGAACCCAGCTGGCAGGGTTTCTTGGGAAGAGTGCAGTCAGCTG
TCTGCTAGCATTCAGCGATTCCAAATCCAATTTCTATGGTTTCTGAATCAGTTCCTGGGATA
TAAATGGAAGGGGGGAAAAAGCAACAATGATGCTTCCAAGAATAAAAGCAGTTGCAAGT
CTGGAAAACCCCCAAAACTCCATTCGCTGGAAGGAAGCTTTTCTGAAGGCACTTGAATCC
CCTTCATTTCTGCATAGTGTCAATGGAAAATGCAATCACTCCCATCTTGGCCCTGAAATGC
TTCTCCAGCGACTTCACACCTGGGCCCAAGATCTGCCAGGGGCCAGCTGGTCATTGTGC
AGAGGCTGGTGGCTAATGGCCAGCGAATGGCAAGAGAAAGGAGAGCGGGGCTAAAATC
GGTCCCAAATAAAGCATGAAAAGCAATATTTGAAATTCTACTTCCGGATCCGAACAGGAA
ACAGAATGGAAACCCAGTAAATCACAGACTGGAAAGTTGCTTTGCTCCTTAATTCCCAGT
CACACAGTCTCCTATGAACTTCCTGGTTTTGGACACATGCTAATTGAGCTATAAATAATTTA
ATGATCGTAAGACATTTAACATGGGAAACAAAGGCCTATAAATCATCGGGGTCAGTGCCA
ACGAAAATTGGGGGCTTTGCCCAGCCTCTGTCCGCCTTTCCCAGTGTGTGGGGTGGGAG
AGCTGCCCCATGTGTGAGGACGCCCCGCAGACCGGGCAGGCTTGGCTGCAGCAACCTG
GATACATGCTTCTGCTTTATTATCTTCTTGGGTTAGCAGTGATCAAGTCACAGAGACATCCA
TGTCTGCAGAGACCCCTTTTCTCTCTTTTTGGCTACATTCCTACTGGCACAGAATACATAAA
TAGGAAAAGAAAATAGATATTTATATTTTCCATTATGCTGTTTTAAATCTACTATGTCTAGAAT
TTTATAGCATGGGATATATTACATATTCTTTTAAAAATAAGGTAAATTTTCTAAATTAAAAAAA
AGCTCAAAACTTGTTGTCTCTGGAAATTCTCATAAGATTCCTAGGAATCTCCTCACTTCTTC
CCTCCCCAGCAGTATCTCATCATGTGATGCCTCATAACAACATACACTGTGTGTTTATGCTG
TTATCTCTCACATTTTTCTTTCTGTGCCTCCTCATGTACTTTCTTACTTGGTTATCCGGATAAC
CTTCTCAGCGCGGAGAGGTCGCCTGCCGGAGGGAACCCACAGCAAATGCACAACCAGC
CAGCAGCCCAGAGCTCCCTCTTGTTCTGAAGTCCAGTGGAGAACTGTTTTAGAGCTCATT
CAAAAGGAGATGGGCTTTATATTTTTGTTTTCATTTGCTGAAACACCAGACAGGCCCTTTT
GTCATGGAAATAAATGCAATCCTAGGCCCAAAGGAATGATAGAATTTGTCCTGTGAGTCAA
AAGTTTATACCTAAACCAGGAGAGGATATCCCTTTAAGAGCAAATTTGGAATTAAGCCTCT
CTGCAAAGGTAATATTCTGACTAGCAATGACTCACATGGGAAAGCGAGGAAGGCTTTGAA
TTACTGCTTCAGAACCTGTCTTGTCTTGCTTTCTGCTTGGTCGCTGGCAAAGAGGAGCTGC
CCTAACCAGTAAGCAGCAAGCATTGCAGTCGGCTTTGTGCTTTGTAGCAGGTCCTTGATTT
TTTTTGGAGAGGGCCATGAGGCCACACCTCAAGGGCCAAATCAGCCTGCAGTCATTATTA
CCTGGGAACACATCCCAACACCCGATTCTTCCATTGCTGTAGATAGTCCTTAAAATTTTCC
AAGAATAAAAGCATCTCAGTTTAACTATAGAAAGACTTGATGCTTGGGAAGAAGATGAGG
GTATCCAGACCCCTTTTGTTTAGGTTGCATTAATATGTTTTATGTGGAAATAGATTTTAAATG
ACTTTGCAGATCACATAATTTAATCCTCTGTGCACATTAATCTATTTCCTTTCTGATGTTCAA
GGTTTGGGAATTTATTTGCCTAAGGAGGCAATAAATTCTTCTTCCTATGGGACTGTGCTGCT
CATTCTGCTAAATGCTGTCCCTCCCACCTTCTCCCTTGCAGCCCCCTCCAGCGTATCACTC

TATTGCAGTCTCACTGAGCAATGACGGTCCTCACAATGTATCCTGCTGATTTATATGTTTAC
ATAGGCATTTCTCTTTCCCCACTAAGATGTAAGCTCCAAAAGAACAGCAACTTGCCCAGC
CTGTGTTGAGGTGCTTAGTTCATGTTTGTGGAGTTCAAATCACTCCTTAGGAAGTTGCACA
TGAAGTCAAACTTTGGCCCTGTTTCGCTTCCACCCATCCATTGCCTCTTATTCTGCCACCT
GGAGTCCTACAGATTAAATCGAATTTCGGTTTGGAGTGACTGTGCTTTGAGAATGGAGTCC
TACAGATTAAATCGAATTTTGGTTTGGAGTGACTATACTTTGAGAATTTGAAGATGGCCCTC
CTACCTGCAACTCCCTTCCCCATCTCTCCTGTCATGACTAACTCACGCCAGTTCTTTCAAC
TAGTGTTCATGAATTCCAGGCATGGCGACTTGCCTTCAGCCTACTGCTATTTCTCAACTTCT
GAAAATGTAGCATGTAAAACTCACAAACCTCTGACCACAGATTTCACTTCCCAAAATTATT
CCACAGACACTCACTCAAGTACCTAAAAATGTGTGAACAAGGATGCTCATGGCAGCCATT
TATGGAGATAACCCACACATTCGACAACAGAGGAAGAGACAGTCACTCTCGGACCCCTCT
GCACACCGGCACCCAGGGCAGCTCTGGCGGAGTGATACGGCCCCTGTGCGTGAACATG
GACAGTTCCGAACATGAGTTATAAGTTCTAAGTCATGACTAGAACATAAGCATGATTCTATT
TTTGTTGAAAATTCTGTGGGAATGGCCTATGGCCATTTCTTATCACTCAGGGCTTTTCTAGG
CTCTTGCATGACAAATAGACTAACTCAGGGTCAGAGATTTCTGCATTTGGAATGAGTCACA
CTGCAGCTAAAAATCACCTTTCAGAAGTTCTGCCAACTTAAAAAGAATAGAACTTTTGTCT
CAACTTCATCTTTCAAACACAATAACTAAAGAGGAAATACAGATTTTTCTAAGGCATCCAAT
AGGAGCCAGAAAAAAGATGGTTCTTTTATACAGCAGTGGAGGTGTTCATGAGCTGTTCTG
TGTTTCTTCACATTTTTATACATAGTATAGTACCATTTACATGAGATTTAAGTATTTTCCTTAGT
TTGCAAACCATTTATTTTACAAGTAGGCGTGTCTTTTGAATAGCTTGTGCCAGATAACGGG
GGAACAGGGGGAAGATAATGCAGAGTGGGCAAACTCTATATAGCATATAGGCACTTGTGC
AAAAAATAATCTTCCAATTTCAAATGCAAATATTGTTCTGTCCTCAAATCTGTGGCCTTGAG
AATTTGGAAGTCAAATGTGCTCTGAACAACTGGAATTCACTGCCGTGATCTGCAGCATTCT
CTGCCTGGCAGGGCAGTGCTTGGGACATGCAAGATGCTGGAAGAGATGAACTCTGTGCA
GATGTCTCATGCCCTGCAGCATCCACTCTTCACAAATTTCCTCTATTTGGGGTTATACTGCC
AGTATTTCACATGAAATTCATTAGACGTGAACATCTGAAATAATAGATTTTTTATATTGCTTA
ATTCCATAGGTGCCACGGAAAACAAAAAAAGAAAAATGGAAAAACAATAGGTCTCTGTCA
TCAAATAACATCAAATAACAAAACCCAATGGGCATTTTTCTTGATTGAGGGAACAATATTCT
TCGAGTGATCGTTTTGCTTGTCATGTGGCTTGCCAATGGAAATAGGAAAAAAGTACACAC
CTACTATCAGGAAGAAAAAAACTGGGGAGGGAAAGAAGGCAAGGAGCAGGGAAGAAAT
ACATAATTGTTACCACATTAACTAGTCTGTATGCACAATCAGAACGTTTGAGTTCCAATAAA
CCAGATGCTGCCAAAATATGCTGCTGCTTCTATCTTTTCTGAGTGTGTATGTGTATACACAC
ACATACACACACACATATATTTATTTATGGGCAAAACAAACTGACTTTTCCTGTATTTTAATT
TTGCTTGGGAAGGCACAAAGTTGGCAATATATATGTTTCTACTTACATATTTAGGATGTTTAT
TGTGAAAGCCAATTCAGGAATATTTGAACTGTAAGAGTTGAAGACAATTCCATGGGTAATA
AGGTAATGTCTTGACTTGATAGATTTTTACCAGGATCTCTAAATATGATTTTTTAAAACCGAC
TTTAAAATGTACACTTTACAGTGTACATTTTACAATGCAGCTTTGAGGATAAATGATAAACC
ATCACATTAAGGATAAATGTTGATAATAGGATACATCAGACTTCCAGAGTTTAGGAGACATG
TCTTAGAATTAGAGTCAGGACTAGCCTTTTGCATGAAATAATTAATTTGAATAGTTTTACTTT
AAGTCTCTACATCAAAGATAACATTTCCTTCAGGCCTGTGGATTGTGGAGTGCACAGGCA
GTGCAGGCCTGATTTTTGGAGGGATATGTAAAGGAAACAGAACTGAAGATTGATAAACAA
TCCCTGGACATATTGGTGCCCAAGGAAAGCAGATCCATTTTGCATTGAATGAGCAGGGCA
AAACTGATTAGGAAATGAAGGCATATTGCCACAGAATGAAGGGAAAAGGTGGGCTTGCC
CCACTGGAGGGACTTACTAGGGTTCTCAGAAAAGGGAGGATGATGACAATATTGGTGGCT
TCTGACCCAGAATGGTTTTGTGAGTTAGGGCGAGGGGCCCTACAGACACCTGGACGCAC
ACTGTTCAACGTACTGCTGGGGGAGCAGTGAGCGTGTGCGTACCTGACACATCCAAATCC
ACTCCAGGATTGCAGGCACTGTACTATCAGCACTGCCCTGCACTGAGCTGCTGCTGGGTT

TTTGGAATAGCAGTGGCTGTGAGTTTCTGCTGATGTCATCCCACATTCAGCCCTTGCCGCT
AGGCTCCCAGACCTGGGTCTCTAGGTCTCCTGGGGACTCTGCACTACCTGATACAACTCC
TATACCAAATTCTTAAATTTTGTTGTTTTCAAGTAAGCACACAGAAATCTATGTGTCTCAGC
CCTTCTCATGTTATGCTTAAAATTTTGTCACATTCTCCACATCTGAGTTATCTCACAGGAAA
GAATAAAAATAAGTTGTTTCTAAAAGCTTATCAAATAGAACATTTGTCTTAAATTATGTCTTA
TCTACCAAACAGATGTGTTTTTTTAACAATCAAAACAACTAAATTAAAATAGTTAATGCATT
TGGAACAATGGTTAATATAAATAATCTTTAGGATGAAATCAGAGTTCTTTCACTCTTTCTATA
TAAGCCCTGATTAGACTGATGTTTTGTGATTCACCCTGGATGTTGTCAAACCCTGTAGAGT
CAGATGTGTAGGTGGTCTCAATTTGCACTAAAGCTCCCTAAATGCTTCTCCAGTTGTGCCC
ATGGCTAGTTCACAAGGCTATCATTTGTGCAGAAGCTGTTTTCATGAAGAAAAGGAGGGA
AACAATTAGTGTCCACGTTAGGATACTGGAACTACTAGAGCCTGGTTCTCTTTGATCATGA
AAGACTCTGTTTTATCAGAAAGTAATAGAGTAATTCCCTCTCTGCCAACAACAGAAAACCA
TTTTAAAATCTTAGCGCTGACTTTGAAAAGCACTTAGAATGAGAATAAAAAGCAACAGCAA
ATGATCTAGTTTCATCTCAAATACTTGAAGTGATGAAGGCATAATCTTTCTGCAGTTTTCCA
CTTGTGCATTCCCCACTCTTCTCAGCCACAAACCACACCAAAAGGTATGTGGTATATTTCT
GAAGGAGCCACATTATAAATGAATGTATCATTCTTGCCTGAGCATTTGACATCAGGGGAGA
TGGCTAAAAATAAGCAATAAAGGCAGACATGTAACAACCGCAAGCTCCTCTAATAATAATT
TAGAACATTCATTAGACTCCAAATCCAAAATCAAAAGCAAGCATCCATTGTACAAGCACTT
TATAAAGGGGCACCCCTGCAGCATAAAGTGACTATATCAGGCAAAAAGCTTGCTATTAATA
TTCCTATCCCTGGAATTCTTACTCATCCAGATTTCCCCCTTCAGTTACTTTTTCTCTTTCACT
TGAGTAGTTTTCTGAGTGCAGCTGATCTGCATTGGATCAAATGAAGCCAAAGCACAAGCA
ATACATGACTATAAATCACTTTTCCTGACAGTCTGCCCACTTCCTACTGCGTTGTAAAACAA
CAGAAAAAATGTGGGGCAGTAACATGAATTTTTGGATGAAAAGAGTCAATGCCCCTCAGA
GTTACTGAGGCAGCCTAGCCTCTTACAGTGGGTTGGGAGAGAGGGGAAGAGGGGCAGT
GCCTAGGCACTGGATCAAGCCAGGCAGGGCAGAGCTTTGGGACTTCACACGCCGACATA
TTTCCACTAAGAACATAAATCGCTCCTCATCATTTGTGAAACAGTTACCGCATGCGGTAGA
CAGTGCCATATGCATCCCTGGAATCCTTCAGATAACCCCCACAGATAGGCTCTAGGTTTTC
TTAATTGAGAGATGAGACAAATGAGGCTCACGAAAGTCAACTTGCCAAGGTCATCCAGCC
AATAGGCAGCTAGGGCAGGACTGAAACCCGGCCTGTCTCAGAATCCATGGGGCTCCACA
TCTGCATCTACCAGGCATCATCCAGCACAGTAGCAGCGACGGCGGAGTTGGGAGCCTAC
GGAGGACGGTTTCAACCCAGCTTGGCAACCTGCTCCCTGCACGGCCTTAGGGAATTCCA
TGGCACTTGCTCTCCATGCCTCGGTCTCCTCACTGAATTGCCTGCAATGCCTTTTGCCACA
CAGATTGTGGATGAGGGGCCAAAGCACTCGGGATAGTGCCTACCAGACAGGAGCATTCG
AGAAGCTGCTCCTTTATCGGTAGCAACAGGTGGTCTATCCTACTCAGTGCTGCCCAACAC
TGCTTCTAGACAAAACAGGCACACGCCCTTTATAACTCCTATAAATTGGATTTGAGTTGGG
CCTCGATTGACATTGCTGTTTCTTAATATGAGCACAGGAACTTCTACCCTTCTGGCATGGA
AAAATAAAAATGAATATAATGCAAGTAATGAATTTTTAGAAACAATGTCATTTCAGAACAGT
TTCAGTCTTGTGAATTAAAAGATGTGAGCCTTCTATGCAGCATCCTTTATTCTTAAACCAGA
GATGATGAAATCTGTATGAATAGTCACATGTCATGTCTGGTAGGCATAGGTTCAAGTATATA
TAAGGATGTGACAAGGAAGAAACAGTTTCCAGAATATTTAAGAGTCTTTTACAGGTCATAT
TTCTACAGACTCTTATGTTTGAAAGAATGTGGCCCAATGTGGACCATCCAATATCCAAACA
CCCACGACATCATTCTTTGGTTAGATTTAACTACCACCACCCACAGGCACATCATTCCCTC
CCTAAGCATTACCCTTCTGTAGCTCCTTCTAGACCAGCTTTCTTCCTTAGCTTGTGCTTTAA
ATGAAAATCTGCCAGCTTTTCACTTAGTCTCTGAATGCAATGCTGCAGGTTCATAAATGGT
GGATCACATTCTGTAAGGGTATCTTTACAAAAGTATATTACAGGGTGTTAGGAAGGGAGAT
ATATTTGAAGACATATTATGGTTATGCACTTCTGCAATGAGACTGTCTAGTTAACTAAACCAT
AATATTAATTAAGAAATCAGTTTAAAAAGCAACCCTCCCAAATCCCTTTCCATGTATTAAGC

ACACAATTCAGATTTGCTCAGAGAGGACACACACTGAAGGGCTTGCAGAGACTGGGAAA
GGTCATCAACTTCATAAATGGATGCTCTCAAAGTTCACCAACAGAGGATGCCTCAAAGAA
GGGGAGGACATTCGGAAGCAGCTCACCCTCCTGAGGAAGAAAATACTGAAACTGCACTA
AGAGTTCTGATGCAAAAAGATAATTCATGACCCATATGGGGAGTCTATTCTTATATCTTTCT
CCATCTTCCCAAACTCCCCAAACTTCTTCAAAGGAAGAAGAAACTATACAGAAGTATTTTT
AAAATCTACAGAAGTCAAAGCACGTGGCCTTCTTATTACATTGTCTGAAATGTTCCTCTTTC
AAAGATGTTAAGGACTGCAGAATCTGCAAGATTTAATTTTTCCCACTCCTCCATGCTTTAAA
CAAAATCTGATTTCACCATCTAGCTAGGTATATCAACCAACAGTTACATAGTGTGCCTCCTG
AGAATGTTTATCAGGAGTTTTAAAAAAATTTTTAAAATTTCTATTTAAGATATATAAATGAAC
AGATACTTAGATAAACATACACATTATACATGTGGTATATAAATTTCATTGCTTAATTCTGACC
CTAAAGAGTTGAAAAATGTTTTTTCCAAAAAGCATTTTGACGCAAACATCAGGGGAAAAC
CATATATTGTTAACATTACTTTTGGTAAAAAAAATTGTTACATTGTTTTGCTGAAATCTTATTC
TTTTGACGAGCTGGGCCACCAAGAAGCAGCAGACGCTGCAGTTCAGTGTGCATGTCCAA
GTCCAGTTCAACTTAGTGAGTGAACAATATGCAAAGAGCAATGTGTTAGCTGCTTTTAAAG
CAGAGTTGAGTTAGACATAGTCCTTCACCTGGAGAGCATGGCTCAGGTAAGATGAGCCAG
ACAGACCAGGAGAATGTGGAGTGGAGTGCCCTGCTGGAGGTCGAAGGTGGCTGGGGTT
CCTGGAGGGAGCAAAGCCATCTGGCTGAGAGATGGGGAGAGGAATTTGGAAATGTCCTT
GGAGATTAATCTTGAGGGGCAGGTGGGAATCTTTCCAGTAGGGGTCTTACAGGTGGGAGT
GGGGAGCTGAGGCTCTGGGTGAGGGAAGGTGGTGATGAGTGGGGGTGGGTCACATAAG
TGGGCTTCTGTGGAGCACAGGGACCTGCAGGGCAGAGGCAGGGAACACGTTCAGAGAG
GTTTTCAGAGCCACACAGTGACAGGCTTTAATGCCAGAGTGACAAGTTTGATGGAAACTG
AGGGTTTTAGTCAAGAGAATGTCATGATCTGAGCCGAGCTTCGATACAGAGTTTCCCGAC
ATAGCACAACCTCTTCCCCTCACCACGTGATCCAAGGCTTCTGGGCTGCGTGGAAAAGG
GATGGGAAGAAATTAAGTCAAAATTGCTGAACCTCCGTTGCTCAACCTTAGAAAATCCTAG
AATACACAGTATGCTGTGGAGAACTGTTGACAGGCAAACTTGTTCTAAGTTTCAAAATGG
GAAAGAAGATGGAGATATTATAAATACCTATCTCAGGAAAAAATATAGACAGTTTAAAAAA
GTTGCTGTATGTGCATGTACAGACAGACTTGTTACTCCTAAATGCAGGTCATGTGTGGCTA
ACCTGTTTCATAGCTGGGCAGTACTACAAAGCGGACAGGTCAGGGCAGCAGCCTCAGCC
AACGCTGTGCCCGGACGCAGCATGTGCGGCAAGTTTCTCCTGATTCAACTGCACCTTAAA
AGGAAGTATGCAGGCTGGCTGATAGGATTTCATGGATTCGTGACAGGTTAAAAAAAGCAC
AGTGATAAGAGGATGGATTTCTCAGTAGGTGAGAATTAAAATGGTGTTAAAATACAGAACT
CAGCCTCGCTCCACATCCTCACGGTAATTCAGTAAATAGAGCGACAGAAAGAACACGGG
GAAATGCTGGAGCAGAACTGTGTCTGTACCGATGGCATATCACATCTGAGTGTACCTCAG
TAAGCTGGAAGGAAGTGCTGAGAACAGTAGGAACTGCAAAAGGAACAAATGACGAGGA
CTTAGGTCCCAAAGCCCAGTGACACAAGCGTAAAGTCTGGTGGACAGGACAGGGCAGG
CTTCCTCGAGTTTCCAGTCTTCCTGGATACCTTCTGCCCGGGCCCCTTGTTCACATCTGGC
AGCCCTTGCAGAGGTTGTCAGAAGGAGCAGAGTTGTCAGAGGGTTGCTCGGAGGAGGC
TGCTCAGCCGCGGGAGGAGAAGAATCAGCTGTCTGCAAGCCCAAGGCCAGCACGAGGA
GAAACAGTCTTGTGTGGTGGAAAAGCACCAGGTAAAGGTGAGGCAGGGACGCCGACAC
TGGTATTCACAGCAGCAATTTTTTATGACAGTTTAAAGAAAAATTGTTGGAACTGTCAAAA
AAATGGAGCATATTGCCTCTGGCAGTGGCAAGTTCCCTGTCATCCAAGTGTTCAATTGACT
TCAGGGCCCCGCCAGCCCTCCAAGACTAACAGTGATGTGTGACGTGAGGCTTGGCAGCT
GACTCTGGATAGAGCCTCCGAGTGGCTCAGGTGGCCGGCTTCCTGTCTGCGTGTCCCAG
GAGTGGCCCACCCCAAGGACGGCGTGTGGAGATGACAATGAGGCCTTGGCAGGCGTGG
GTCTCCAGCACGGTGCTCCACCCAGGGCGGAAGTGGAGGTCCCCTTTCAAAGCCTTGGG
TGACCACAGCTTTGAGCAGGGTCACTCTGCAGTAAACAGCTGCAGTAGCTCTTAGATAAC
TTAGAAAACAGACCTCTCGTAGGGGAAGAATTTCCTGCAGACATTTCTCCCTCGAGGGCC

TCCCACCCACCTCAGAGGCTGCACCCAAATCCTGAGGATCTGGCAAGAACCATGTTTTTG
AGATGTAATTAATACAACTGTTTACACACAGCATCTGCTGGGGCAGAGATGGCGGTTGTGT
GCAAGCCACAGCTGTGATTGGAGCAATCTGATGACAGTGAGCTCATGGTGTCCCCAAGG
AGGCCTTTGTGAAGTGCCTGACATAATTAAAATTGGGAGGAAAACTAGGTCAGGTGACCC
AGGGTGGTTTAGGTGAAAGGGCTTCAGAGCTCCCAGCTCTGCCCATTCCCTGGGCACTG
TCAGGCATCGTAGAAAGGATGCGTGCTGAAAGGCAAAGCCACCACAGTCTCCGCTGAGG
AGCAGTGCCTCAGAGATGCTGAGAGATGCTGCTTTCCATTCAGAGTGGCCACCACTGAC
ATCTCTGACAAGACTGGCGGCAGTCTGCTGGCTGGGTCTCTCAGAACAAAAATTTAAAAA
AGGATTGAAACCATAAGATGCAGCCCCCCTTGTATTATCTGTAAACACCAGCACATTTGAT
GCATCGGTTCTGTCTCAGATTTCAAATTCAATTCATTAGCCTATCCTTTTCCATTAAAAAGTT
TCCACTTACAGTGTTGAGAATTGGGCCCCTTTCTTTTTTTTTTTTTAATTGCACATCTGTTCA
CAGAGGTTGGCAAAAGACACTGGAAGTGATTGTGAAATCCACATTGTGATTCCTCAGGAA
TCAGATCCTAGAAGGGGGTGCCCAGAGCTGTCGGCACACCGTCCCAGGAGTCTGCCTGT
GCAGCTCCCAGCCAGGCAAGAAGCCCTGAAGGCAGAGTCCCAGGTGGACACAGCTGGA
CGCCTCTCTGACAATGGTGGCTCTGGTGGTGAACCCCTCGGTGTCTCTTCTGCACCTCTC
AAGGCTGCAAAGTGCCAAATACTCTTTTCCAACCAGCTCCCGAATTCCCCCCTCCATCTG
GGACTGCATGTCCTGCTTAGCGATTTCAAGCAATGATTTCACCTTTTCATAGACAAGGACA
TTGTCCTCATCAGGGCTTGCACCATCACTCTGATTTCCAATGCTGACAGCCAAGTGGAGG
TTTTTAATTCTTAAAAGAATGTGGCATTAAAAAGAAAAACTCACCCATCTCATGGTCTAAAA
TAAAGCAAAACAAGTCAGATGGCTCTGCCTAACCCAGCACAGCATCTGTTACATAATCTG
AAACTTCTGTGTAATTGAGTGAGATGTGGGAGAACTCAAGCACAGTCTTTGTAGCATTCTG
GGCATGGAATCCTGATCCGTTAATTCCAGAGACTGAATCAACAGAGGAACACAGAGAAA
GCCACCACCACGATGTGTGAAAAGGAGAGATACGAGAGAGCACACAGAGGGCTGTTTTA
GAAATTGATTTTCATCTTCTGTTTGAGGCCCTGGGAAAAGAGTGAGAGAGTAGAGGGAAA
ACATGGACGTTTATGAAGAGCTGCCAGAGGCCAGGTTCTCATCACTTTCCAGGGCCTCTC
AGGAGAGGCCACCTGAGGCTCCTGTCTGACTCCCCACTGCTTCGAAATCAGAAGACTCG
GGGAGTCAGAATCAAATCACTTAATTAGCTCACAGACAACACTGGACACCTTTTAGGGTTT
ACTGGTTCACATACTATTTCAACTACTTTGTTGAATCTTTTTAACAAGTGGAGGAAACTGAA
GCCTTGAATCACAAAAGGCTGGCAGTGTGGAGGGCCAGGTTTGCCTGTTCTTACTATCTG
TTTGATGACAAGACAAATTAGTTATTTTATTTTTATATTGCCGGAGTAACTTTTTCTTTGAAG
CAAAATTAGAATTATAGATAAGCAAACAGAAAAAATACTTAACAACAATATGCATACAGTAC
TTATATTTCTCAGGATTCTTTTTATGCATACGTAGTAAACAGGTATATTTATTTACAAAAGTGA
GATTTTACTGCACATAATTTCTGATGCCTTAAGGTTTTTTTTTCATTTAACAACATATAGTAAA
GTTTTTTAAAGCAAATATTTATCCAAAAATACATTTTACTGGCTGAAGAGTATTTCATTCTAT
GGAGTTGCCATCACGTCTCTGAGCAGTTCCTGACTGTGGGAAGGTCATGCGTGCTGTCG
GGTCAGACCACCCTCAGGAGAGAGCAGTTCTCCCCAGTCCCCCTAGCCCCATCCCTGCC
AGTCCTGGCTGCCGAAGGCTGCAAATTAGTTATTTGGAGAAACACAGGCTATTTAGGTTTT
GAAAAAATCAGTGAGCTTTCAGTTTCTAACGAAGCTACCAACACCCTCATTTTTTCCCTGT
CTTTATGGTCTGCAACCAGATGGAGGCTAAGGATAAAGATTATTTTCTCCATCAACTACACT
GTCAGGCAAAGCAACTCTGGGGTCTGTATGAGAGGAAATATGGATTTTATTTATGTTAGCA
GTAGTTACTGTTGTGAGGCACTATTTGAGTATTTTTCATTATATTTTAATTTTTTTGAACTTCC
TATGATAAAACCATCTACACTTTTCTAAGTAGGAAAAATAAGGGAATGAGTGTCATGTTTTT
AAAAATTTGGGGAAAGATCAAGAGTACAGAAGAGCATGGGGCAAAAAAGAAGTTTAGGT
GCATTTAGGTGACATCAATAAAGCCCAGTTCTTTTTTTTTTTTTGACAAATGGGATCATCCTAT
AGACATTGTTAGGCAAATTACAGAATCTATCTGCGCTGTCCCTAGTTGGGCACAACATGCT
CCATGGCAATCTGTCTTGCTGCTGTGCAGTCCTTCCCTGGGTGGCTGCAGGGGCAGACA
CAGGGTTTGTAGGTCCTGGGTCTTATAGAATCAGGTACAAAGTGACACGTATATTTAGAA

GGAGAAATTGCACAAATGAGACAAAATGTTTTCCTTATGCAAATTTCATAATATAAATTATA
GACACTGCCAGTGCACTGTTAGGACTCCCAGGGCCTTGGAAGGGGCTTCAGTATTATTGG
CTTTACAATAGATCTGGTCCAGGTGAATGTGCGATGATTTACTTCTCCATTTTCTCTTTGATG
GATATTTGGGCTGGTTACAGTTTTATTGTTTCCTATTAAAAAACACTGGGGAATAAGTTTTAT
GGGGGAGGATTTATCAGCTGAGGTTACTATTGCGATGACTGTGTGACCTTCCCTTCCCCTG
TAGATCTGAGAGCATCCAGGCCATTAGGCAAGGAGGATAGAGGAGGCTGGGATTCTGCG
TTCCAGGCAGTACTGCCTTGAAGAAGTGCAAAGACACCACCAGAATCACCTGGGGGTGC
TTGTGAAAACTGCAGATTCCTGGGCCGTGGAACTGGGGCAAGGCACAGAAATATGTATTT
TGACAAACATCCTCAGTGGCTTCCTCACAGGTTAGAATTCTAGAGCCAGTGACTTGGGCT
TCAGAAAGAGCACAGGTATGCAGCCTCGGGCAAGCCACAATTTCAATTCAGTCTCCTTAC
TAGGGAAAGGGAGCCAGGGATATGCGTGTGGAGGACAGGTGAGGAGCTGGGGGGGTAA
GGAACGGACAGCACTTTCATTCCACTACTGTAGCTGCATTCCCCAAGCACATGTGCTAGC
CTCTGTATTTTAATATAGAACCTCGGCTGCACCTCCCATGACTCCAGGAGGGACCCTGTGC
CCTTCCCTGGCTCCAGCAAGGGCACAAACCGCTCTGCGCCTGCGCCCCTGCAAGCTGGC
AGGGGCTGACAGGTGCTGTGGCAGGGCCTCTGCTGCAGCGTGAGGGAGCCATGGGTGG
GGAGTTTCTTTTCAGTGGGACCAACACGTGTCCACTTCATAAAAGTGGCCGTGCGGCAGT
TCCCACAGGGCTGCGTGGCCGGGAACTCCTGGTCAGTCACTTTATTTCAGAGTGGAAAA
ACATTTTGGCTTCCTAGATCCCTGAGATCCTGACACTTCCTGTCTTCCTAGAAGTAAAATG
GCCATAAACAGAACTTATCACAGGTGACCTCTGGCATCGTATCCCCATCGCCTGATGTTAT
TTTTACAAGGCAGCCGAAAAGATAGCAGGATTTCAAATCTGGCCCTGCCGTTCTGCATTG
CATGTGCAGAGGTCTAGTCCACAGGCCGCCACAAAGTTAACTCAGCAGCAAATTTAAACA
CATGGGGTACACATCACGGAATTCTTAATCATGACCAAGCACCTGGTGAACACAACCGGA
GAATTAAAAATCTGAGGTATGTGTATCAGTGTGCCTGAGAAGATTTTGAAGAGAAAAGGCT
TAGGTGGGGTCTGGGAATGAGGTAGAAAGGGCTGCAGTTGGAGCCAGAGGGCCTGGGA
TACGCTTTTGTGCTGAACTCACCTCAAATCGAATCTTCACTTTCTCTTCTATAAAAAGTGGA
GAAAAACAAGGCTTCTTTCACTCCTCCTGAACTAGCTAATATTACTTTATCTTTATCTCTTTC
TCCTCCTCAACTAGCTAATATTACCTTATCTGCACGTGTGTGCCTGTGTGTGTCTGTGTGTC
TCTGTGTGTGCCCGTGTCTGTGTGTATGTCTGTCTGTGTCTGTATCTGTGTCTCTGTGTGTA
TCTCTGTGTATTGTATGTATGTCTCTCTGTGTGTGACTGTGTGTATCTCTGTGTGTTGTAGGT
ATGTCTCTGTGTGTCTGTGTGTATCCCTGTGGGCTGTGTGTATGTCTCTCTGTGTGTGTGCC
TGTGTGTATCTGTGTGTCTCTGTGAGTGTGTCTCTGTGTGTGCCCATCTGTGCGTATGTCTG
TGTGTGTGTGTGTGTGTCTGTGTGTATCTCTGTGTGCTGTGTGCGTGTGTCTGTGTGTATTT
CTGTGTATTGTGAGCATGTCTGTGTGTAACTCTGTGTGTTGTGTGTATGTGTTTGTATCTGT
GTGTGTCTGCATGCAATCCTTTGAGTCTATAATTAATTCTCTCATTGCACATATCAGAGAATT
GTGGTTCTAAAGGCGAAACTGGCCCACACTCCCAAGCTGCCCTGGGCAGGGAGCTGTGT
GTTGAACCAGGACTGTCTCTCCCAAAGGACAGGGCCTAATTCAGGACCCACTGAAGCAG
CTTCAGTAAAATGACTGTGGCACGCCTGCCTGTGAGAAGGCAAATAATGACAATGACAGT
AATAACTGTCAACCTCATTGGGCCACTCTCCTCACTTAATCCTCACAACAGTCCAGTATCT
TCCCCACTGTACAGAAAAGGATGCTGAGTTTTGAGAGGTTAAGTAACTTGAGCTTTTGAAT
GTCACATGGTGGAGGGCCAGTCGGACAGCAGAGAAGAGGGAAGGTGACCTGGCTTGCA
GGGTTGCTCTGTGTCTTGCGGGACCTGCAGCTTTTCCAGCTTTTGCACTTAGACCAGAGC
CCTGCAGGAGTGGCTTGGGCATGAGCTGTGCTAGCGTCAGGGAGAATCCACCTCTGTGT
CCTTGCTTTCTTTAAAAGATGAATCTTTTAATCACTTTTATTAGGAGAAATAAAAACCAGCG
ATAAGAGAGCCCTGACATAAGGCTGCCGGTGCCAGGGATCACCTGGGTTCAGCACCTAT
GGAGAGGTTTATGGTTGCCGCTGCTGCACTGGGGCCTAACTGCCAAGGTCCAAAATGGT
CCAAAATGTTTCTGTTAAATTGTTTTGGGTCTTTCTAGGTTTTAAGAAATATTTGTTTTACCT
GCAAAAGAGCAGGATCTGTCTAGAGTCTGTACAGTAAACACAAGTGAAAATGTAATCCCA

**TATCAATTAAGATGTATAGAAATACACCTAAAGCACTATGTATTTATTTCTTTTTACCCTACTT**
**CTCTCAAAGTGGATCTGTGATGTCCATGTAAATGCAAAGTCTTGGAGTATCCAGCATATGT**
**GAAGGGAGAAGTTAGAAAGGAAGTTGCTGGACTGCCTTGTCGTAAATTCCTCTTCTTGTT**
**TTCACTGTGGACTCCTAGTAGAAACCCATGACCCAGATGTTCTCACTCCCTCCTCTGAGGT**
**CACCTGTAATCTCCAACTCAAGGACTCTCCAAAAATCTGTGGATCAACTGCCTACATCTTA**
**GGCCCTTGAAAACGTCTGGGCAACTGCGTCTCCTGACAGGATAGGCCAGTCTCCCCAGA**
**TGATCAGTGGGTGGCAGGAGGCTGCACAGTGAGGCCCCCCCTGCCCCCCGACCTGGCA**
**GCCCACAGCTCAGGAAGAGTGCTCTGCAGGCTGCACAAACTTGGCTCTTGCACCAGGCA**
**GCTGCGTGGATGACAGACCAAGAGCTATCTCTCCCAGGATAAGAAGCTGGATGCTGGGT**
**GAAGCCAATTCATTTTTCTCCTCATCAGTCAGTGGAAACTCACTTCCACAGATTGATTTCC**
**AAAGGAATAAATGATATCACCAGATGTCATCCTGTCAAGCAGCCCCCTTCAGAAAAAACTT**
**TTCCTTGGTAATGGGGGAAATTTGACTTCAAGGAGGCAGTACACAGCTGTGTACAATGGG**
**TTTCATCTTTTTACGTTACACACGCTCTACTGTGAGTTCTCAGTGGCTGCACTTCCCATCTC**
**GCCTGCAGCTTTGCCACCATCACAACATAACTGATGCTTTACGAGGCCAATTTCATAGTGT**
**CTGCTGGCACCTCTGCACTCCCTGAGCTACCAGTGACAGTTACGACAGGACCAGCTGGC**
**TAAGCACCTGTGGCATCCAGGCATGATTCAGGCAGCCACACTGCCCGAGAGAAATGTATG**
**GAAGTGAGACCTCGGATCACAGACTAGTTTCATCTCCCTAAATTAAGATAACTCCTTGCAA**
**AACTAAGACAAAACAAAATGCCATCCACCAGCTAAAAGTTTCCATCAAAATTGGAATGAG**
**TCTAAGTGCAGTTTTCACCTCAAAAATGACTTCCTTGTTATTTACAGTCCACAGATATGAAC**
**AGAAATGACCATTTCTATGAAAGAAGATTTTATGATAGGAAAGGACAAGGGAAGTAGGGC**
**TGAGGCGCATCACGGTCGGGAGAAGGTCCTCATTCTGTTTACCATCACTGAGGACTGAGA**
**CGCGGGGCGAGCGGTGGGGAGGGAGGGCGTGAATTCGCTTACACAGTCACCATGCCATT**
**CATTGCTCACTCTACGATACTGATTTTTGGCTATAAATTGGCCACATTAGTGATTCCAAAAC**
**GAGGTAGAGTAAAAATGAGCAGCACAATTCCGAACACAAAAGCAATTCCATTTTTAAAAA**
**GCACAGAAAAATGATTTTCGTCATTTCAAACACGGAAATCAGAACAAAAAAACATGGAAG**
**GCAGTATCTATTCTCATGGGACATGACTGACAAGTGCATTTAAAGCTTGGTTCCAGCAGGA**
**AGAGCTTACAGGCTTAGGCGCTGGAAGGAACCCAGCGGACCCCTGAGCTCATATTTTCTT**
**CTAGATGTCAGTTAAGGAAATCAAGACAAACATAGGCTGAGTTTTCAAAGCCTCATTTCAT**
**TCTGGATTCTTTTGATTAAAATAGAAAGGCTCTTGTTTTGAAAATGCAGGGCAATCTAGGA**
**AGAGGCACATAAAGTAGCCAATTACATGAACTCTGTCCATGTGGCTTTCAGACTTTAGAAT**
**CAATACTCTAATTGGTTATTCTTAAAAATGCTAGCAATAAAAAGCCCTCCAAATTCCTAATG**
**TCTAATCCAATCTTGAAAGGGACAGATCTTTAGTAACATCATCAGAACCACATTTGTTATCT**
**GCGAATTGCTGGAGATACGGTGCCGGTCCCACACCGCCCGCTGGCCCTGCCAGGAGCG**
**GCTGACGCCGGGGCAGGCCGAGCTATCATTAGCCGGGCACAGGGATGATGTTCATCTGC**
**CTGTCATGGTGCTCGGTGACATCTAATACTACGAGCTTGTTTTGATCTGGCGGAGAAGATA**
**TATGAAGTTGATTCCATAAAGCTATTTTGTTAAATAAAACAGCTTTCTTCAAAAACTGTCTTA**
**ACAATAATGTAATAAATATAGAAAAGAGGTCTAGGGAATGAAAATCAGAGTCATTTTATTTG**
**CTGCATTTGTGGGGGTGGGTAAAATGTAAGAGGGTTGGGCTTTAAGCCTAGTTTTGATTGC**
**AAATGCCCTACACTATGAAATCACCAGGAAATTGGAGCCTGAATGTGGCCACTTAAGTGC**
**AGGCCTCCAAGTGAGGCAGGGACTAATACCACTGATGCAAATCTTGCCAAAGTAATTAAG**
**GCAAATGTTCAACCTGACAGGGAGCCACCCAGTCTCATCCTCTAGGGCACTCCTGGGCT**
**CCTCAAATACATTGTAGGACTCAGCATCTAATTGAATTTTCAGACCAAATGCAAAAGGAAG**
**GGAGGGCCGGTTTACAAAATTTCACTTGCCAATATTGTGCCGCAGAGCCTGTTTTCCTCCC**
**CATTCTGTCCTGAGCAGAGCAGGAACCACAAGCAGCATGGAAGAACAGAGGTAGCCCG**
**CATTAAACAGTTCTCTGAAAATTACAGCCTGCTATCCATTAAGCAGCTGTACAATCAAGTTA**
**GTGGGCTGCAACCAGAATTTTTTTTTTAAGGAGATAGAGTGGACTAGGAAATGCTAGGAG**
**TAGATTAATCATTAATAAACCTTCATTAGAGCATGTATTATTTTGCCAAACTTTAGTTCTCTCT**

**CTTTAGTTATCTCTCTCTCTTTCTCTCTCTCAGGGTGTGTGTGTGTGTGTGTGTGTGTGTGTGT
GTGTGTGTGTGTGTTTACAGTGGGTCAGGGTTTGGATAAGTTGCTCTACGAGATGCCCTCT
GCTTTGAAATCCTCTGAGTGTGAGTCTCCCCAGCTGCACACAGCTCCTCTGAGGCTGAAG
TATCCTGCCAGCAACACAAGCCTGCCATTGTTTAAGTGCTACTCCTGGGCCCGGCTTGCC
ACTAGTCAGGACCAGATTTTAGAATGTAGTAGGGACTTGTATTGTCAATTAATGGAACTCTT
TCCACCAATAGCTTCGACATGACATTTCTCAACTTATTCAACTCTCCCTGGCCTTACTCTCT
GGAAGGTAACCAGGAAGCCTGTACTGTCGAAGGGAGGTGGGAGGGGTGTGGATAACTG
CAGGCCCCACCCCCAACCCAACTGTGGCCTCCCCACACCCAGATGTGGCCCCCGCCACC
AGCTATGGCTTCGCCTCTCATTCCCTGGCTGGTTACAAATCTTTGACTTACACATCTTTCAC
TGCATTAGGAACCACTATCACCAGAACCTTTCAAGATGGTAGTTTTCATCACTTTTTACCCC
ATAGTAGATAATTACAGATGAAAGACTGACTTATTTATCATATTTCATCATTGTAGCGCTTTT
CCTGACATGACTGATCCTTTTCATTCAAAAGGCCCATTTTGCAAAGTTGAGGGCATCCTCT
TTCAGAACGTGAAATTGTGCAAATCCAATACAAAACAAAACTAGAAATCGGGGTCTTTAAA
TTAGAAATTGGGGTGAAAATATTGTCACTATCACTTGCCAACTCAGCGATCTTGCACACTG
GTACCTCTCAGGGACAGCTGAGAAGTCAGATGAGGTCAGGACAAAGTGCTAAACACAGT
GCCTGACCCCCGGAGGAACATTGTATGTCTTGAAAAAGCAAAGTCACATTTAATAACAATC
TCATATACAAAAGTATATTGTGTACATCATAAAATACCCAAATATAGGTTGGGTCCATCAACA
TCAGAATGCATTTTTCAGTTCTTCATGGTTAAAATGAGGATTTCTTCCACAAGTAGGAGTAT
TTCACTTTTTACTAGCCTCATTCCTGAAGCCAAGATGATTGTCTTTAGAAGAAAGATGTTTT
TAAAATGATAAGATATATTTTGAATTTCCAAACAGTTAGTTACATATGGCTTTCTCAACAGAC
AAGCCTGGCCATTTTTTGGCATCTCAGGCTGTATGTCTGAGCTCTGATTTTCAATCAGGGT
TTAGTACGGTATTTTTTAAGAGATTAGGTGAAACTTTCAGGTCATAAGATTTATGGTTTTAGT
AAATCATGTGCTTTCTACCCTTTTAAATGCCTTCATGTTATGCTTTAACATCTATTTTGTTATT
ATATCTGAATCCTAAATAATATGGCCAAGTCAAGAAAATCTACACATAGGACCTTGAAGAA
GATTCATGTGGCAGCAAAAGACCCAGGGTCTGGGAAGGGGCCCGAGGTCAGGGGCTTTT
TGGCTGTGTGACCTTGGCTGGTGGCCTTTGTACTGCCAGCTTTTTCATCAGCACACAAGC
GCTGTGTACCCTGGAGATCATTTGAAGTTCTTTAAAAAACAGAAAAAGGACCTGGTCATC
AAAACTAAAAATGCACGTACAGCTCCTGAAAACATGAAGCACTTTGGGAAGCCGAGGTG
GGTGGATCACGAGGTCAGGAGTTTGAGACCAACCTGACCAACATGGTGAAACCCTGTCT
CTACTAAAACTATAAAAATTAGCTGGGCGTGGTGGCACGTGCCTGTAATCTCAGCTACTGA
GGAGGCTGAGGCAGGAGAATCACTTGAACCCGGGAGGCAGAGCTTGCAGTGAGCCCAG
ATTATGCCACTGAACTCCAGCCTGGGCAACAGAGTGAGACTCCATCTCAAAATAAATAAAT
AAATAAATAAATAAATAAATAAGTAGCATCACCTAGGCTCACATAGGTGAATTTGAACTGAA
TGAGTCCTTTGAGCAGCTTGTATTGACAGGAAAGAGGGCATCCTGCATAGCTGGTTAGAG
GGAGGCCCCAAGAAGCAAATCCCGTGTGGTGGTCTCCAAGCCTGGCTGGAAAGAAACAT
GGGCTCCGGCAGGTTAGGGAAAGGTGAGAGGTGGGTCTGACTTTTGGATCATGTTTCAG
ACAAACCACAATGGAAAACACATTTTTGAGACAACTGGAGATGTGTCAGTAGGCTGGTTG
TTGGGTAACATAAGAGATTTATGGCTAATTTTTAAGTTCGATGACAGCATTGTGATTATGTTT
TAAAAATCATGCTTTTAGAGATTCATAACCATGAAACGGAGATAGTGTTGTGTGAAAGGAT
GGGATTTCTTGGATGAGACATGACTGGCCTTAAAATAGTTCAGAAATGGAAGGACAGATG
TGGAAGAAAACAAACTTGATGATTTGGGGTTCTGGATATATATTAAAACTTATTCAAATAAC
TGAAAATTTAAAAATTCCTCCATTGAAAATCCTATTTGAATGGCACCCTCTGCCAGGCCTC
CCTAGCAAGCCCTCCTTTGCGTGCACAGAGAAGTACGTATGAACAGTGCCCAGTGTACGG
GCTGCAAAGCCCAGAACAAGCTAAATCCCCTTTTTAGTGACAGCCTCACAGAAGATGAGG
GAGCCATTACATGCATAGGCTGACTTACACGTACTTCAGTTCAAAAAGCTCTCAAGCAAA
GTAAAATTAAGTTAAAAAAAAGCAAGAGGCAAAACAATATGTATGAATGACATAAGCTCAT
ATACAAAAATATAAACAAAACCTAAATGATCGACCTTTATTTGTACATATATTTGTATGTATGT**

AAAATACACAGAAAAAATGCTGGAAGGATTCTTCCACAACTCTTGACAGTGGAGTGAAAG
CATGGTTCTTGCTTTTTACCCTCTAATTTCCTGAAATCATTATACTTTTTACAAAGAGGAATA
TTTTTGCATGATGTACGTAAAATACAGCCAAATAAATAAGTAAGTAGAGACAAAAAATGAT
GGAGGGCGTGGGAGATGAGGCAGGGAGGCTGGATGGCCAGTCTCCAGAACTTGATGTTT
TCATCGTCAAAGGGACAGGACTGAGTTGGATAACATTTTGGTTTCAATTGCGAAGGCCTTT
CATTCCAGAAAGAAAACTGTGTCAGCATTACAGCCATGGGTTGACACATGGTTAAGGTCC
CATTAGCAGCACGATGGAAGCTCCCCTGATAATCACAGGGAAAGCCAAATGACGGGAAA
GGTCGCATGCCCAGAGGAACTCCCCTGTAATTTGTATTGTAATGAATTATTACCTTAACACA
AGCATGTGCCAGGCATCTAATTTGCAAGTGCGGATGCCTGGCAGTTGACTTTAAAATCACT
CTGGAATTATCCCTGTATAATCTGAACAACCAGAAGTGTGACCAGGGCTTTACCAAAAGG
GAGTTGCACAAAAGGGTAAACTGCAGACTGGGAAGTTCCATGAACATGGAGGTACTGGA
CGCATCACGCTCTTTTTCAGGAAGAAGTCCTACACACATTTTCTGCGTCTGTTTTATAGACT
GGACTGTGCACTCTTTCATGGGGCTCTTTGTGCGAAGGGGGCCTCCCCCATCCGCTCCTA
CCTTCAGCGTAGGGCCAAGTCCAAGTGGAATACAGCCTCTGAAAACATGGCTGCTTCTCA
CACTCCTGGGAGCCAGACCACTGCTCCTTTCCATGTACCTCAGGCTTTGTGGGGGCTTTT
TATTCAGGTGGGAGAGAAATGGGCACTGGGGCAGGTGTGGTAGATGCCAAAGAAGTGAG
AATGTGTGTGTGTGTGTGTGTGTGTGTTTATACCCTGAATTATATCAGAATGGAAGAAAGGAA
ATAGTATGTCCATCTGTTATCTCCTCTGTGGGCCTACCTCAATTAGCCACCAACACACATAC
ACACCTGTGCTGCCTTTGATGAAGGAATTAAGAAGGCCCAAAGTGACTCATGTCACTTCC
TTTCTGTGGCCTGCAACTCCAAAAGGAGAAGAGGAGAGGACGAGGGAGGGTGGGAAGG
CCATTGCTCCTGGTTCCGGGGTGCCTCTCCCCTTGCCTGCTCAGGTTCCTGTAGGCCTCA
GCTGTTGGTCAGTCCTTTCCACATCTTCGAGGAGACTCTGCCTGACAGGGCTGGCTAGAA
GCTGGGTGCCTGGGTGTTCATGCTCTAATAAACCCAATCGGGAAGTTTCCAACTCATCAC
AACATCACAAAGAATGTGCAAGAAAATGCCGACTCTCTTATTTCTCCTTCAAAGTTTGATT
AATGATTCTAGGAGGAACCAGAAATCCTTGGTGCCAATCAAACACATTTTGCCATACCTTT
TAAACATTTTACACACAACCCATGTTTCATGAAAGCATGGCTCTTATCCCTTCGGGTGCATA
GAATAAGACTGCCGGATAGAATGTTTCTGAGACCTGCTCTCTAATATTCTAACTCAGTAGG
CCCTACGCAGGGCCTGAGAATGTGCATTTTCAGTAAGTGCCACTGCTGATGTTGATCAGTT
CTCTAAGAAATTCTGGGTTAGCTATGGGATTATAAGCATATCCTCAGAGTACAGGTTTTTTT
TTTTTTTTTTTTTTTTTTAACAATTGAGAAACTACTTAAAGTGAACCTAAAATGGTGTGAGCC
TGAAGTGCTTGCTGGGCAGCAGACAAGCTTTGCCAGCCTTGTGTTCTCAGTGGAAATGCA
GCATCCGAGCCATCCTCTCTTCCCCCCGCTTGTGGTCAGCTCTAAATAGCACACTCACAG
CGCGGGGGGAAAATATTTTTCCCTGTTTCAAGTGGGCAGTGGAAGTAGTGAAAGCCTAAG
TAAACTCTGACCATTAGATAATGGGCCATTATAACTCTGGATGACTTCCTGAAGGACCCTG
AAAAATGACTTCTCATTTCCTGCCTGCAGAAAAGAGAAATATTAGGATAGTGTTGTGTGCA
AAAAAATGCAAGCTTGCAATGAGAGATGCAGAGTGTGAGGGAGAGAGGCACGAGGGGG
TGGAGAAAAAAGACAGAGAATTTGAGGTTGACTCACGGCTTTGAAGGGAAAACAAGAAG
AGGAAGAAAGTCTGTTCTCCCATGGGTCGTCAAACCCCACACTTCACACATCTTTCCCAT
GGGGCTGCCAGCTGCCGCCACATCACAGCTGTGCTGCCACCCAAGTGTGCCGCCTCCAC
AGTGGGAGCGACCTAAGCGTGGCCACCTCTGCCTCTGGAGGGCCTGCCTGCTGGCCAG
CGGTGGCCGCCCCAGACCCGGGTGACTGGGCAGCACAGTAGCCCGCCTGGGCAGGGCA
TGAAATTAACGAGGCAAAAGAAGCCACAGAGAGATGAAAGCATAGAAGAAAAAAAGTCA
CCCTGCAGCTCAAATTAAAACAGAAAACTCTGCTGCTATACTTGGTCTTATTCCAAGAAGC
TGCCAGGAAAGAATTGACTCACAGAAAAGCCACAGCTTCATTTTTTCCCATACTCAAGTG
ATATGTCTTTCCAGGGCTTTTTTCTTCAAAGGGAAAGGGAGCAAAATAACCTCACTTTTCT
GCATTACCATAGACTATCTTATATAATTTTTTTACAGCCTGTAATGGCTGGTTGAGATACCAG
CTGACAGTGTAGTGGATGAAGGCTCCGGGGTGCAAGCCCCTGGCTGTCCTGGCCAGCTC

TTTAACTCGAGCAAATTACTTACTCTCTCTGAGCTTCCCTTTCCTCGGCAGTAACATGGAG
GTATTGAGCAGAGAAATTGTGAGGCACTGTAGACACGAGATAATTCACAAAAATTGCACG
GCACGGGACCTGGCACATAGTCAGCACTTCACAAACAGTGGCTCTCATAAGGTAGTTAAT
AGTGGTTTCAAACTATGTGTCAAATCTAAGGGAGGAAGAGGGGCTTCTAGGGGGTGGGG
ATGGGAGGGAATGTTCCAGTACAAATCTTGGTATTGTCAAACTTGGGGCAGCATGCTCAG
GAGACAAAACACGGAGGCACATCTGCACAAGCTCAAATTCTACTTCTGCCACTGCATCAG
TTGGGCCACTGTGGCTCCTCATTTCCTTATTTTTAAAAGAGCTGATGTGAGAACGAGCTGA
TTTAATGTGTAAAATACCTAGTGCCAGCAGACAGAAGGTACCCGCTACACACCAGCTCTG
TACTCTGCCTTGATCCTTCGCAATTTTCGTTAGCAGAATTCCACCACGCACTACTGACTTTA
TGGTATGGCTAAAACCAGAATTCTAAATGTTCAAGGATTCTTGTGCAGTTACTTCAGCAGG
AGTTTTCAGAAAGTGGGAGAAATACTATTTATGAAGCACCACCCATGTGCTTTAGTCTTTCT
ACATTGTTAGGTTTTCATGATAATACTTTAGAGATAAGAACACTCCTCATCTCATGGACAAC
GGAACGGGCTCTACAAGAGGGCGTGCACAGTGGACCTGGGGTCTGACGCCCACCCTGC
TCTCTGCACCTGGAGCCCACCAGCATCCTGCTCCCCATGCTGCCTCCCCTGGAAAAAGA
GACACATGCAAATGATGAAGCATGCAGGAGCCACATGGAGGTTTTATAAACTTAAAAGGA
TTTACATGAATGCCTCAGAGAAGGGGCAACCCAATGTCAACTCTGATACCCCAGGCATGA
TACCACTGACAATGTGTCTGCTGGCATCAGTTGGAAAACAATGCAAATATCCTTCATGTGA
TTAAACACAACATGCCTGTGACATTGTTTTTTTTTCTAGCCAGGCTGACTGTTGTCCTGTAG
GCTTCCAAAGGCTGCTGGTGCCCTGACCTTCAGCTCCTAAATTACCATCTGAAGGACAAG
GACTGGGGAAGGGAAGGAGAATTTGCATTTGAGCGTACCATTCCGTAGGTCATTCCAGAG
AAACTGTGCCTTCCTCCCGGTTGACTCTCCGCTCACTGAGTGTGACAATATGCCTGACTAT
GGAAGATCACGGATGGGGCTGTACTGCCCTGCTACCGAGAACCTGGACAGAAGACAGCA
GGTGAGCCCTGGAGGCTGGAATGAATGGCTAAGAGCATCTCAGGACTCCACTTCTCTGA
AGGCAAAATCCAATATGCCAAATTCCTCTTCCTAATAAAAGCCCTTTAAATATTTGAAGACA
GCTATTATAACTCCTCTTAGTTTTCTCCCAGATAAATATCCACATTTGCCTTCATCCGTCACT
TGACTTTTCTCAAGTGTCTTCCCCATCCTCTGACCATTCCAGTTTGTAAACATTCCTCCCAA
GTGTAGCCTTGTAACTGAACTCTTCTAATCAACAGAAGCTTGTCCAGCTTTATTATATCTCT
TAATGCAACTTGACCCAAATGCATGATAAATATCAGTCCCATGGTTAGGTTATATGGCAAAG
TTGAAGTTGTGCAGATGTAATTAACATCCCAAATCAGCTGGTTTTGATTTAATACAAAATGA
AGTTATCCTGAGTGGGCCTGACTTAACCAGGTAAAAGCCATGAAGAATGGACTGAGCCCT
CCTGGAGGCACATGCTTTCCCACTGGGCCTGGAGAAGCAGGCGGCCATGCTAAGAACTG
CTTATGGAGAGGGTGGCCTCTAGGAGCCAGGACCTCAGTCCTACAGCCTCAAGAAACTG
AATTCTGCCAAGAACCCTTATGAGCCTGGAAGAGGACCTGGAGCTCCAGATGAAAACATA
CTTTGTGAGTCCCTGAGAACACAGACCCTGACCCACAGAAACAGAGATAAGAAGTGTGT
GTTGTTTTAAGCTATGACATTTGTGGTTTTGTGTGCCAACCTCATAACTTGGGCTTACATCT
CATCATGGTAAGCCAATCTCTATCTTTCATAGGGGCATTTGTTAAACCACCTTTCTTCCACT
CAGGATATTTGTTACTGATTTGGCCTTGACCCTGTTATTAATAAATTCCCTGATTTTGTTCCA
TTCATTTGGGTCTGTATTTTAGGATCCCAATCTTGCTAACCTATATGTTCTAATCCCTTCCTC
TTGATTTCTTGTATTCCAAAAACCTGAGGCACTCATTTAAACTGGCCAATCATTCCCAGGAT
GTTACTTCTTGACTCATCAAAGTAATCAACAGAGACAATGGGTTTCATTTTACCAAAAACTT
CCAGTATATTTAAACATTAATGCATGTCTGCATTATATTAGTTCACATATTTTTAAAACACAAA
TTCGACTAATAAGTAGCCATTTTTTTTCAGGAAGAAAAAGATTCGCTTTCTTCAGTAGATGC
AGCAAAGATCTCCCCTTGGCCTGAAAATAATTTAAAACCATGATATTAATTGTACACACATG
CGCACAGTGTGCCTCGCCATGTGCCTGTTTTGGTTATACTGCAGACAGGCTTTCATGCCCT
TCTAAGAGTGACCAACCATGTGTTTTCTCATAAGACATTCACCAAAGATGACATTCAGCAG
AGAGGTCTAGACCACTGACGTCAGGCGTCTGTTACAGTCCTGGATGAGGGAGGCCACAG
TGGAGTGAGTCAAGAAGAAAATCCCAGGGAAACCCTTTACTGGAGTTCAAGGCTGAGTT

**ATTGTCTAAAGAATACCAAGGAAGTGAACATGATGATTGAAATACATTCTGGAGATCATAT
GCTTCAGGCTGTCAGCGATGCTGGTGACATTGCCATGGTTAATGCTACTAAATAAAGTCAA
CGTCTCCAAGAACGAGTGCATAAATTACTATGTGTCATCAGTTGGATTGTTCAATAAGGTAT
GTTCAAGTCCTAACCTCCAGCACTTGTGAAGTAACCTTATTTGAGAATGGGGTTTTTGAAG
ATGTAATGGAGTTGAAATGAGGTCTTACTGGTATAGAGTGGGCCCTGATTCAGTGATGGGT
GTCCTCATAAGGAGATGGAAACACAGACACACAGAGAGGAATGTGAAGGCACAGAGAC
AGACATGCAGGAGAACTCACTGTGAGGATAGAGGTGGAGATCGGAGTTTTGCATCTACAA
GGCAAGGAATGCCAAGGGCTGCTGGGAACACCAGAGGCTGGGGAGAGGCCAGGGGCA
TCATCCTCAGCAGGTTTCAGAGAGAGAGACTCTACTGACACCTTGATTTTGGACTTCTGG
CTTCCAGAGCTGTAAGCAAATAAATCTGCAGCTTCTTAAGCCATGGCAAGGATGGTGATCT
GTTAGAGCAGCCCTGGGGAAGCCACACACTACGGCTGCCCTGCACGAGATGTCTCAGGA
GGGTCTCAGATGAGGCTGCTGCCCGGAGCCACAGGGACGAGTGGTCTATTGTATGTCTAC
ACTGGAGCGAAAATGAACCATGAAATGAAAAAAATGAAAACCCTGGGTAATAATCCTTAT
GAGACAGGAAGAAACAAGAAACAATCATAGTAAGATGCTGTCCTGCTTTCTCCCTTGGGT
GGTTTTGTGTCTAGGCAATCTCCTTTTTAGCCTTGGATGCCACCCACTGAGTTTAAATACAA
CTATTGCTGTGAGTCTCCAAAAGGCTATAAAGTCACAGGTAGGAGACAAGGGAAACAAAC
AAGCCAAAACACCAAAGCTGCTTATGAAGAAAATGTGGGCAGGGGTGGGGTACAGAGG
GGAGGAAAGGAAAGGGAGGGGCATGTAGGATTGTGCATCCTTTACAAGTAGGAGGTGTT
CAAGGACTACCTTTTTTCTCTTTGAGGTATCAGGAATGACCCAAGTTCCTTCACATATCTGA
CACTCTAACAGCTTTCTCATCCTGATATCACTTGGTAAGATCAAATAACTTTGGTTTGATCA
CTGATCCACTAATGTCCCCAGTTACCTACTTTGTACAGTGAGAGGAAGGCCCTAGATTGGG
ATTTCAAACTGTGGGACACCACCCAGCCTGCATTGTGAAATCACTGGAGTGGGTCACAAC
CAGCATAACAAGGAACAAAAAGAAAAATAAACAGAAAAGAATAGACAATATCAGAGTGAT
ATCGTAAAGGCAGTATTATGAAAGTTTAGATACAGCTGTATGTGTGCCATGATATTAAAATG
TATTTCTTACTTTGGTAAGGTCAAAAATGTTTGAAAGCCACTGAAATAGATGTTATCTGGGA
AACTTTCCTGTGCTGCTTTTTGTGAGCCTATGATTAGCTCCAGAATTCCTTACAGAGTAAAT
TAGAACACACTGACAGCACGTTTTGGAGAGAGAAAAAAGCTTGCAAATAAAAAAAAAAA
AACCAGTCTAAAACAATCTCTTCCTGTCACAGACACTGATGAATTGTGCAGTGATGGCCC
CTCCTCGCCCAGGGTTAGTGGGTGATTTGGTAATAGCCTGCTTGTCCTGGGAGCACGGAA
GGATGGTTTTGCTTTAGCTATGGAGAGACCGAGGGGCAGAATTCAGGGGCTGCTCTCACC
TCTGCACGATGGCAGGTGGCTGAGGACAGGTCAGCTCAGGCATCTCAGGCTCTCGCTCA
CAGACACCACACTTTACCACCTGGCCTTTGTCGGGAGTCTGCAACACTCATTGCGAAGAA
GAAAGCCAGGTCCAGTCTAAACAAGACTCCTTCGTGGCAGCAGTGTGAACAGTCTAGGA
AAAGATGCAGTTACCCTGCTCAGCACAGCCCCCTCCAACCGTCCACAGAATAGAGTATCT
CCGTTACGGTGTGTGGCCCTGGCCCAGACTGTACCAAACAATACAGTTCCTTCTGCCGCC
AAAAGTCCATGATTGATAATTGCCCTGTTAATCATTTCCATTCTGATCTGCAAGGAACCCTG
CCAAAAGTATGTTTCAAAATGCTTTCTGTTCAAATTAAATTGCTGTTGTTTTACATGAGGAA
AAATACTACCTGGGAAAATGTTTGACAAAAATGTGGCAATTAAGTGCACTTTTGAAGCACT
TAAAATCCAGGATTGAGCTTTCAATCTTTGATGTGCACCTTTTGCAGGGCATTTCTAAGTAG
TATAGTGGCCAAAAGGAAGCTTTATCATCCTGACTGGCACCCATACGTCCTGATAAATGGC
TAGTGTCACTAGATCACCCCTGGGTCCTAGTTGTACAGGGAGGGCTATCCCACACCTGTC
CCCACCACATCCACGACAAGTGAATAACATTTATTTTATCTTATTTTTAAATTTACATAAAGA
GACAGTATATATTTATGGTATACAACATGATGTTTTGATATATGTATCCATTGTGGAATGGCTA
CATCAAGCTAATTAATGACTGTATTACTGCACATACCTTTTTTTGGTGTGGTGAGAACACTT
AAAATCTACTCTCTTAGCAATTTTCATGTATACAATACATTGTCACTAACTACAGTTACCATG
ATGTGCAATAGATCTCTTTAACTTATTCCTCCTGTCTAACTGAGATTCTATGTCCTTTCACCA
AATCTCCCCAAGCTCACACCTTCCCAGCCCCTGGTAACCACCATTCCACTCTCTGCTTCTC**

TGAATACAATTTTTTAGATTCCATGAAGGAGTGAGATCATGCAATGTTTTTCTTTTGGTGCC
TGGCTTATTTCACTTAACATAATGTCCTCCAGGTATGCAAATGACAGGATTTCCTTCTTTTTT
TTTTTTTAAGACTGAATAGTATTCCATTGTGTATGTATACCACAAGCAGTGACTGGCAGTTA
CTAAAAACAAAACAAAAAGAAACAAAAGCTCCTGATTATTATAACTATTAAGTCTTGACATT
GGCTATCATGTTTTCAACAAATGGTGCCAGAGCCACTGAACATCACAGGCAAAAAAAGAA
CCTTGACCTCAACTCTTGTGCAAAAACTAACTCAAAATGGGCCACAGATTTACATCTAAAC
AAAATACTATGAAACTTGTTGAAGATGACCTAAGGGAAAACCACAGGACATACTTTGGTG
CAGACTTCTCAAACCTGACACAAAAAGTATGATCCATAAAATAAAAAAATTAAAAATATATT
AGACTTCAAAATTTAAAACTTTTATTCAGCAAAAGAGCCTGGTAAGAGGTTGGAAAGATAA
GTTATGGACTGAGAGTAAATATTTGCAAACCACAGATCTTTCAAAGGCGTCATATTGAGAA
TATAAGGAATGCTCAGCACATACAAATGCCCCTCCTGGCAAGAGCCTACGTGTGTTCAGC
TCTGCTGCAGACACCTGGAGTGGCGGGACCCTGGATTTTCACGGACTACTGGGCTTGTTT
CCTTCCTTTCTGTCCTCAGGTCTTAAGGTTCCGTGGGAATGATGAGAGCTATGAAGCCAG
CAGGCCCGGGCCTAGGGTGGGGCACGTTCTTGGGCCCAAAACTTAAGACGGCAGCAAA
GGACCCTGTCTTCACGATAAACCGTGCAGTATTTTGAAAACGAACAAACAAACAAAAAAA
CGCCATAAAATCCATGCTGAACAAAATCTCAAAATGTAAATGCAGACAAGACCCAGTGCG
TGCCTGTCTCAACCATCACTCCAGTCCTGGCCCTGGAAGGAAAAAAATTCAAAGAACAC
GTATTTGGGAGAAACAAAGTAAAATGTGCAATCCTAACACAGTCCATTTCTTTTTGTAAGT
GCTTCTCACCACACCTTCAGCACGCACAGAGCAGGCCGCAGCTCCTGCATCTTCAGCAG
CACGCCGACCCGCCCCTCTTCCCCGCTGCACAGGCCACCTACTCTGAGTGTGCATGCCT
GGCTCTGGGCGGAGCACCTGCTCCAGCTCTGCAGGCCGCCCACACCCCGTGCCTGGCC
TCTCCTCTTCTGACCTTTCCTTCCCCATCTCGAGTCCCCTGGTGGTAAGTTTTCCCGCCCC
CGCCCAGCCTTAGCGGGGCTTTCCCTCCAGCCTCAGCCCATGCCTGCCTCATTCCAGGG
CCTCGCAGCTCGAGCCCCTCCTGGCTCCTGAAGGCTGTCCCTGCTGCCTGCCCGGCCCA
CGGGAGAGGAATGGACGAGAACTCCCTGAACTTACGTTCTCTTTTCCCTTCTGAAGCCAA
ACGGAATAGAAGTACCCGCCTGCTCCCTGGGGTGGGGAAAGAGGTCTGGAGGGCCCCG
GGCACCGGGAGTCTCCACCCCAGACGCGTGCCTGGCCGGAGACAGGGGAGTGGGGGC
TGCATGGGGAGTGGAGAACCTGAACTTTCTGTCTCAGCAACGCCACTCCACTGCTCCATG
GAGAAAGTGGACGCTCTGGGGACATGCACAGGATCCGCGGGCTGCCCAAACCATCTCA
GGATGTCCCCTCTCTGTCCTCCTGAGCACAGCCACAGACCGTGGAGTCACCACAGCGTC
CACTCTCTCACCAAAGCGACTGGGCCCTGCACGTGGGATCTGCATGGGCAGTAACTCGA
TCCCACAGATGATCAGTGGTGGTGCCCGTGTGGCCCCCACCCTGACAAGTGCACGTGGA
CATTTTCCTGCACGTGCGGAAGGACGCAGGTCAGTCTGGGCAGCAGTTCCCCACGGGGG
ACCAGCTCAGAGGGAAGGAGGAGGACGCACATAAACCCTATGCACCGAGGCATCCCAG
GGTTGGTGCAGCTAGCACACATGCTTTAGGAGTGACAAACACGAAGTGAAAAATAGTGTC
TCTACGTGGCGTGGTACCCTGGGGATGGGGCTGGAGCCTCAAGATACGGCTGCGTCGAG
GCGGCTTCCTGGCACCGCTCTGGGCTGTGGCGGTGGGGTGGGGTGGGGGGTGTCCCAG
GACAGTGAGGACGCGGGACGCGGGGCTTGGGGTGTGTGCGGGGCCCAGGGCCTGTCC
CATGGACAGGGTGGTCCTCATCCCCTCCCTGCCGCCTTGCTCCTTCTCCCCTCCCCACAC
GCCACACACCCAAATACCCCAGGACAGGAACACGGCACGGTGGACAGGAGCAAACCCT
GCACAGTGCCCAGAAGGTCTGCAGGAGCTTCTTCTTCCCTTTGACTCGCGCACGTTATTC
TAAACTGTTCCTGGGCGTACCTTACCCCAAGAGGGCCAGCCACTGCCTCCCCCTCCACAC
CCACGGCGTCACCGGTCCCTGCAGATAGCGTCTCCCTGGTGCAGCATTACACCCTTGCAA
AGCTCCGCTCGCCAAACAAAGCTGCGCTGACGGCCTCCAGTTCCTGCTCAGGGTTCAAT
GTCCAGCTCTTCGGAGCTGCCTCTGTTTAAAAGAAAAGACGGGCCTGCATCTGGAAACTA
CCTCTCCCAGGTTTCTTTTTTAAGAGCATTTAACCCATTTTTTCCCCAAGCACCAAAGAGTT
TCTGCAGCCCCATAGAAGCCTCCCGCTCTTCCATGAGAGCCTGGAGTTCTTGCACCGCCT

GTTTCCTAGTTCTCTAGCAGTCTTGGTCTTTCGGTGCAAGCAGCTGCCTTACCCTGAGTTA
CCCTGGCTTAGTTCCACTTGGAAGTAAACATTTCTGTCATTTAATTATATCATAATCATGCTG
GAAGGAATTAGACCAGCAAGTGTGAGGCAGGGTGTGCAAAAGGGCACACAACACACAC
ACACTTTTCACTGTCAGAGAAAAAAAAGGAGGACACATTTTTACTTCCAATGTAAAATGACA
AGATCAAGATCAGAAAATTTAGTGATGAAATTGTACTATGATTTATATTGCGCTGAATTTTAT
TCACTATTTTAGCCACACAGGAAGATAATTTCACAGCCTTAAGATGATATTTTTTGCTGAAA
TTTTATTTAATTATTATCAAGGAATTTTTCTGTGTGCTTGAGGCCCTGTGCTAGGCACACTG
ATGTCTCAGTTTAAAAATACCACACCCAGCAAAGAATAGTAGTCAATAAAACTGCCATATTT
TCTGCTAGGCTTACAAAGGATGAAGATCTGACCGAAATACCATCTTTGGAATCTTTGAGGT
CTAAATAATAATCTGTAATTGACCTCATGAAGACATTGAATGATTTACCAACTCATCCATTAG
TACATGGTGTTCAATAGTTGTACTCACATATTGGGATTGACATGGATTTTTGTATGGCTTACA
TTCTGCCTTTGTGTATCTTATGACTGTAGTGAACTCAGTTTTCTAATGTCCAAGCAGAGGAT
GGCTTAGGTCTTTGGCTGCTAGTCAATTTCCCTCATTCCCTAACTGTTTTATTTGTCATATAT
TTATTGATAGCCTACATGACAACAGCTAATGATGGAGTGGCAGGCATTATGAAAATAGATA
AGCGGGTGGTCAGAGCTCTGATGGAAATTATGTTGTAGTATATATATTCTGACATACGAAGA
ACTCTTCAGGAGTGAATGAACCATGTGCTCTACTAATGAAGTTCAAAGGGAAAGAAACTA
CAGTATTCCTCAGTGGGGGTGGGTGCCATGCACAATAAGTAGGTTTCATTGCAATGAGCAT
TGAAAATCACCTAGATGATAATAGGCGAAGGGGTGTGAGTATGTTGTTTTGTTTGCTTGGC
TACTTTTGGTTGGTGTGTATGTGTAAAAATAGGCCTATCAGATGATGGAATAAAAAGAGGA
ATCTTTTTCTGGACATTAGTTGTGTTTCTAAGTCTAGGCCACAGAGTGTCCTAAAAGTTGTA
ATCATTAAAGTTAATATGCCTAACAGTGCTTATCAGGATTTTTTTTCCAACCAATGAAACTAG
ATTTTACTGGTGCCCTTGAGTCTTGATAATAAAACTCAAGGACATCAGTCTTGTATTTGCCT
GCATTTCCCATTAGATTGAAGGAAGTTACAAGGCTTTCTTGGGTTTTAAGAATATTTGACAT
GCGTTACCATTTCTGTTGATTTATTCATTTGTACTGAGAATCATATTTTAATTCAGAATTTCAG
CAAGGTGTAGGCTTGAAAAATAAAGCCCTCATGCAGCTTGGATTTCGTATTAATACATCTC
AAGCTTGTACAACACTACTAACAGAATGTTATAATAAACCATACCAGTATCCCAAAGAGAT
GCTTTAATACCAAATGTCTCATTTATAACAAATTACTAATGGACAATTGCTAGGCTGGACAA
GTGAACAAGGACAATTAATTGAGGTAAACAGGCTTGGACATTTCCAGTTGAATGGAAATTT
CCACGCGATGTGCTTGCTTGTCTCTTTGTAAAGAACTGTTGTGCAGAGTTCCACTATTGCA
CCCCCACCGCTTGAAAGCCAGCCCATCTAATTTAATGCTCTTTTTAGCTCAGCAAATTCTG
ATTTGGAGAGTTGATGTGCATGGATTTTTGTGCAATCTTATTTACAAATCAACTGAAAAAAA
AATAAAGAGTTCCAATCCCAGAGTACACTGTGTAGCTCTGAGCGTGCCAGAGAAGGCAG
CTGTGGTGCACTTGTGATGGCCTCTGAGGGCTCTGGGTGCTGTCCTCACTACGGTGGTAG
GTTCTGCTGCCTGAGTGTATCCATTACAGGAACTCGAGCTTCGGGTGGCCCATTATAACCC
AGCTCAAGAGGATGTGTCTCCAAGTTTCAGGAGGAACATAGCTGTACATCAGCTGCAATG
TTCCTGTGACCTAATCTAGGGGGAAATTCAGATTGGTAAATTTGTTCCAAAACACATTGTT
GCATTGTTGTGGTAGTGGTTGCTAAGTCCATTCCCCAGTGAATATATTCTGAAATTAGATAT
AAACTATCTATATTATAAAAGCCCTAAAACAGGAAATTCTGTCATGTTGGTCACAGCTCAAT
TAATAAAATGACCTCCATACAAAGAAGGGTTCAGAGTGAATGAAAAGTGGCCACCGGTGT
GGAGGAAGCTGGAGGCCTTCACAGATAAGTTAGGCCCAAACTGTCCTAGGTAGTATCCTC
TGGTTGGTGGAATTATTTGAAACATTTTTTTTTTTTTTTTAGCTCCTGCTTCAACTTTAATTTG
CTCTGCCCTTTAGAAGAATGCAGGCTGCAACCGTGGCCCCTGCCCTGGCCCTGGCTGCC
CACTGATGGTCTGGGCTACCTACACACCCTTTGGCCTTCTATTCAGTTACTTACAAATGCTC
TTTAAAAGTCTTAAGAAGATGACAAAGTGATTTTACTGGTATTGTGAGTTATTTTAGTTGCT
CTTATACTGATTTTCCATGAATGCGATTGTTTTAATAGTAACCTTTGCCAAAGATTTCTAAAA
TAGATTTCTAAAATACCATGCAATCTCAAGGTTCAAATTCTGGATGAAAATAATGTTTTGTT
GAAATGATCTTCCAAAGAGAACACCGTGCTTGGTAATTAGAATGACAGGTTAAGCATTATT

CTTCAACTACCATAAGGAAAAATAATCTTGTCTTTGCCACCTAATTGATTTGCTTACTAATTG
ATTTAAAATGTCCTTTCACTTTAATGAGTCTGTCTATTTACAGTGGTAGTTTATAAGGTTAAA
GTATGCATTCCCAAATATTAGATATTCTTTAATCATGTCGTATATCCAAGTTGAAAACAGTAG
GTATTTGGGGAGCAAGCAAGACGATATTTAAAATAATTAGGTATTTTTTAAAAATTCATATTA
TTCTCAAAAATAATTCTGGAAAGACTTCGGTGGCCTTTCACATTGAATGAGAGAATATGAT
GTCACCACTTACACTTTCTGTTTCTGTTTTATGTTCACAAACCAGGCCAGGGGGCCAGCTG
TAAACCCACATTCTATTTTGATTACATTGTCATAACTTTCAGTTCTAACTGGAGAAAGCCAG
AATTTAAATCTAGCCAGAGTATTAATGCTGCTCAACCAGGAGGGAAATGAAAGTGGGATCT
CAGAGTGAACATGTGGTCCTTCCGAATTTCCTCCTGGACCTGGACAGGACTCATGGTCGA
ACCCGAGATCCTGAAACATTTTCACCATGACTCAGAGTGAGCAGGAAGGGTGGGGCTGG
CCTCTTTGTTTTCTCACAAGGTTAATAGCGTGTACCTTAGAATGAGCCAGAGCGCAGCAG
GGGTAACAGCACATGGCTTAGCCTTCGCGAGGCAGATGTGGTGAATGAGCCGTTGAGAC
CGCCCCTGACAACGTCAGGCTGTTTGCAGTACCCATGGGCCTCGCCCTCTCCAGACTAG
GGGAAGGGTGTGTCTGAGATGCAATCTAAGAAATGATCATTCTTTAAATTCTAAAGGTGAA
GAGATGTAAACAGCATTTTAATTATAATTTGCCTAAAATGTGTTTCTTGTAGAGGATAGGCA
TCCCTGTGCACATGCAAAATCTTGGAAGAAAAATATATGTTGAGAAGGCTTGACTAAGGG
ACAGAACAAATAGCACGCTGGTTTTCTGTTTTCCTTAATGCTGATAAGGTCAGAAGAGCTG
TGAATGTCTCTATGTAATTAACAGGGTGCAAAAGAACAGGAACGAGATTCAGGGGACTCC
AGAAAGAACTTTAATTTCTGTTCCTTGGTTTCCTCATGGGACACATGGGACGAAGTATTAA
TAGCTATAAATGCCCATAACAACTGAGATTTGATAGTAAATTCCAAATTAAGTTAACACTTG
AACTTTATAAAAAATACACACTTCATTTCCCTATGGGCCATATACAAAATGTGAGCTGCAAT
CAAAGTCCTGGCTAGGGAGTTTCTGGTAACGTGTCTGTATTATGGGATGTACACACTGAGA
AAAGCATTGATCACTGCACCTGCAGAAGCCTTTCCAGCATGTGCCTCACCCACATACTGC
CTGAAAGTCTTCTGAGAATCATTCTATAGTCACAAAAAAATCCCCTGGGAAGGTCCTACAC
TCATACCTATTTTACAGATTGAGGAAGTGGGTTGAGAGGCTTGATAACTTGTGCAAGGCAG
CGTGGCTGGTAAGCAGACACTCTTCACAATTGCACTATGCTGTCTGCAATATCCAACTGCC
CTTTCCTTGATCCGGCACATAGTTTGAATTGAGGGAAGTCTTTCAGTATTTAGAAAAATCAC
AGATGACTAAATGGTAAAATAGCAGAAACAATGAGTGAGAAGTTAGATATTTGCTTCATTC
CAAGGGAAGAGAAACGACAAAGTAGTCAGGTAGTTAGCCAAAAGCTCAGCCTCCAAGAG
CAGTCAGATATAGCAAAGCCTGAGAATTCCTTATCTAAAAGTAGTGAAATGTGAAAGGCTA
ACATGAGAAATTTCATCCTGATTATAAGGAAATTTTCCAGACAGTGAAGTGAATTGAACTG
TAATCATGTACCTAAAAAGAAATCATGAAAGTCCTTTGTGTATGGCATTTTAGAAACTGACA
AGCACACAGGAAAGAAGGCAGGTTCACTGACTCCAGACATCTTTCCATCTGCACTATCTT
TTTAAGGATTACTTTAAGGACCTCAAAAGGAGACAAAACAGCAGACTGTTAAAAGCAGCC
GTCAATATTTATCTAAACACAATGAAGCAACTGCTTGTGCTACGACAAGCAGTAATAAAAT
GATGGGACTACCACTGTTGGTCCTATTAGTGGAAACAATTAATAGAAAAATGGGCCAACAT
AAAACTGTTGGTAACTTTGTAAGGGAAAATCACTTCCCTAAAAAAGGGAACGGCATCAGC
ATCACCACTCATGAGAGCAATGATTGACACTTCACATGCACCAGCAAACCTCATCCAGCC
CGCAGCCCTCTGAGGAAAGCACTGCTGACAGCTCCATGGTGCCCACCAGAAAATGCAGG
TGCAGCTGCTTCAGGAGCTTACTCAAGTCCCACAGCTGGTAAGTGGGACAGCTGGGATTT
GCACCACATTCCTCTGACTCTGATGAACGTGCTTTTGTCCGTGATATAATGCCACAAATAG
AGCTCTAAAATCCACCTTGAGTAGGCAGTATACTCCAAAGTTGTGCAGAGTACAGACTAGT
AATAGGTACAATTGCTAGTAATAGCTACAACGCTTAAACTGATGTTGCATACATTTAAAATTA
AGTTTGCATTTGAACTTGATAAAACGTGTGCATTTAACTTACTTCCCTACGCAGGTGGCAC
CAAAGCTGTATTTGCCCTCGGGGTTCACATCCATCTGGTACGTGGTGGGGTTGTAGAGCA
TGAGTGGGGGGCAGGTGTCCTTGCACGTGGCTTCGTCTCGGAATTTGCGGCAGACCTAT
GGAGGAGAGAGGACACGCTGGGCAAGTGAGGTAAGTACACTCAGCGCCACACACGGAG

**GAAGCGCAGCACGGTGTTGACTCCCGTCAGAAAGCCCAAGTTACTGTCTTTGCAGCCTA**
**GTGTTCCACTAGTGTGGGAAAACCCAGCTGGCGCCCTCTGCAGAGTCAGAAAGACCCTT**
**CTCTAAAAAGTCCATTTTTTTCTTAGGTGCACTGTCCCACCTATCATAATTTATGGATTATGA**
**TAAAAATACGTCTCTTTACTAACGTATAGATGATGGAGTTCACACATTATAGCGGTGACTAT**
**GGAATGAAATTATGTTTAATTTCCCTTATAAAGGACATTGGGCTTCCCTTATAAAGTACAATT**
**GAGGGTATATTTTATAATGTATTGCTGAATGGCAGAAAGCCTTGCTAGAACAATGCCGTGTA**
**AAGGAGCCAGACCCAATACAGCGAGGGCTTTTCCTAGGACCAACCCCACGCACAGTCTT**
**CTCCCTGGGGCTCACATTAGAAACAGCTCAGGCCAACCAAGTCACAATTCCTGGGTCTG**
**GGTCCAAACTCTGGAATTTAGGAATCTTCTCTGAAACCACAGAGTGCTCTGGATTTAGTCT**
**TAGGAAGGGCTTCGTCACACCAGCTCCCACTCCAGGAGTCACATCCACCAGCCTTGGCG**
**TCTCCAGGACTTCCAAGACGCTTCTGACCTGTACCCTCCAGTGCTGCTCAGAGCCCGCTG**
**GGGCAGCAAGGCTTCTGCTGGCACTTTCTGGAGCAATGGGGCGACAGTGAGTACTTCAG**
**GACCCCTTGAGGGCAGGGTGCAGCTCAGGAGGTGCCTGAGCAGTGGTGGAGGAGCCGA**
**GGGATCCGTGTCAGGAGGCCCTAACCCAGCCCCAGCCAGCAAGACCTGTGTGGCTCTCC**
**ATGGGTCTCTTAGGGACCTCACAGTGTCAAGTCTCTAGAGGTATAGCCGAAGTCAAACAA**
**TAAAGAGAGTGTGCTGCTGGCAACAAGGCACATTCATGACAAATACTTGCATGTAGTATTT**
**AGTGCTTGTTCTGTGCCCAGGTCCTAATGACGGACATCACAGGCATCTCATCATTTAAGCA**
**TTCAACTGGGAGAGACTTTGTTATGCATCATTGTCAGGTAAGGAAATGGAGGCATGGTAGT**
**CCCATCATTTTCCCAAGGTCTCAGTGAGTGGTGGAGGCAGGATACTAAAGATACTTTGTCA**
**CCAGGCAGAGGGCAATATCCTGTCTGTTATGACAAATGCTAATTGACAGCTCCCCCACAG**
**GAAGTCTTCTGTCCTGGTGTGGGTGGGGCGGGTGCAGCCCCAGCCTGCTCCAGGGAGG**
**CTGCTGGAGGGGCATCTTACCAGGCAGTCGCTCTCCCGGGGGCCTGTGCAGCCTGCAGC**
**ACACTGGTTGTGGCAGCAGTCACTGGGGGACTTGCCACGGCAGCGCCCGGAGCACTGC**
**TGGGCACAGATGATTTTGGTCACTGAAAAAGAGCAAAGGTTCCCTGTGAGCTGAAGAGT**
**GAGGGTAGGATCATTTCCCAGGGACGTCGCAGAGCACTTGCAGACTTTTTCATGAAAAAG**
**TGAAGACACATTTGCTTCCAACGAAACCACATTCAAACTAAGTCAAACCATGCCCTTTTCT**
**GATTGCCTCTTTTTTCTCCTGATTTATTTTTAATCACTTAAGAGAACTCATGAGATCAAGTCT**
**AGACTTTGAGACCTGCTAAATAATTAGATGCATCTTGGGCAATGCAGAAAGCACATACAAA**
**GAACAAAAATACCTGCTGGGAAGTAGTGTTTAGAAGGGCCTAACAGCCCCCCCACAATTC**
**AGCTTAAGGACATAGTGTCCGTTTTTGAGCTCATGTCCCCAGGTTCCTTGGCCACTGAGA**
**GGCTGAGCCAGACAACCATACCTAGGAGAATGGCCCTCTCCATCGTAAAGCAGGGTCCA**
**CATGGAAAAGTCCCCATGAGAACCCGTGCTCAGGCTTCTGTGTAGGTGGGTACCAAAGA**
**AAGCCAGGGAGGGAGAGGAAACAAGAAGTCTGGTGACTGCTGTCAACAGGAGCAAGTC**
**TTCCATTCCTGGGAAAGTGTGCTGTGTTCAGAGGACAGGGCTGGGGTACCTGCTGATGTT**
**CACTTATTGTTGTTGTTGTTGTTGTTGTTTTGAGACGGAGTCTCGCTCTGTCGCCCAGGCT**
**GGAATGCAGTGGCACAATCTCGGCTCACTGCAAGCTCCGCCTCCCAGGTTCATGCCATTC**
**TCCTGCCTCAGCCTCCCGAGTAGCTGGGACTACAGGCGCCCGCCACCACGCCCGGCTAA**
**TTTTTTGTATTTTTAGTAGAGACGAGGTTTCACTGTGTTAACCAGAATGGCTCGATCTCCTG**
**ACCTCGTGATCCGACCGCCTCCGCCTCCCAAAGTGCTGGGATTATAGGCGTGAGCCACC**
**GCGCCCGGCCCCTGCTGATGTTCACTTATTTGTGATAGGCTGAAGCTCCCAAAATAGCCC**
**ACCCTGGATTAACTGCATGCGGTGAGATTTGCAAGACTGTCCTTCACCTACTGGTTCACAT**
**CTGACCCTGTAATAAAATAACTCTTTGGAGTCTTCAGAGGGAAAACAAATCCAATCACCTA**
**AGCAAGTGAAGGAAGAGAGGGGCGGTCACACATGGGTCTGAGGCTGTTCACTGACTTAC**
**GTTTCTGGCAGTTCTCCTCTCCTGCACCCCAGCAGCTCCCATTGGGACAGCTTGGATCAC**
**ACTTTTGGCCTGAAATGTAAAAGTAATAGATGCATATCTCAGTGAAATGATGAGAAACTGA**
**TGACGCCCTTTCCCGGGCCTGAGTTAAGCACGTTCAATAAATGAAGACAATCTTTTCAACT**
**TATTCTCTGATTGTTTGGCTGGCTGGTTTGTAGAATCACTTGAGACCCCGATAAGAAAAGA**

AGATGAACCATGAACTGCTTTTATTTACATGTCAGCTGCGGAAACTGAGGCAGAGAGTAAT
TACTGACATGGGCCACATAAGCAAGAATTTTTCATGAATTTGAGGTTCCTTGTTTACAATCT
GCTTAGGCTTGGAGCATTCCCTGGCATCTCCTGCCACAGCTAGAGTGTCTCGTCTGGCTG
AGCCTTTGGCATGGACCCACTACTTCCTTGCTGTGTCCAGTCTTGAGTGACTGCATGTGTC
AGCCGTTCTGGGATTGTTAAGTCATAGCCTGTAAATAAAACCTTCTAGACTCCAAACCAGA
GAGAAGCACAGAATGGTGCACAGAGGGTGAAATGTTAACAGTGCAGATCACTTTCATGA
CTGTGTAAAACCTTAGCTATCTGGTCAATTTTTGCAGTTTTTATAATTGGCCGATTACAAATT
ATAGCTGCAAAGGTTTTCCTGTCATCTGTCGTGTTTTCTTTTTATCAGTATTGAGAAGTCTG
AAGGTCGAAGTGCTATATCTATATAGAAAGAGTAGGTTAACCACCAAATTTGTTAAAGAGT
CCAGTCATGCAGATATCTTTTACTGTGAAGACATAGTAAACTAGAGGTGTTCATGGGTCCC
GAGGTCCCCTGCAGTCAGGAAGAATTTAGATTTCAAACAGAAAAGCTGACGATTGATCTC
CCATGTGACTGTGAAGCCAGAATACCACCAAGGCAAAGGGCGCTAATTGCCGCGCTGGT
GCTCCTTAGAGACATGAAGGCCGGGGAGGCTGGCATCACGACAGTGACCGCGGGGGAT
GGACGTTGTGAGGAGGTAGGCAGAGAGACTGCAACAGAGGTCTAGAGAAAGCAAGGTC
CTACGCAGGCGTGCTAAAAGCAGCTAGATCTGTAGCGTGCAATCCCATTCTTTCAACTTTT
TACCCTGTACTCACATATAATTTTCAAATTTCTATTATAAGCTTGTAATCATTGTATAAAATAA
AAATATTTAAAAGAAAAATGCATCTTTTTCTAACGGGAAACATTTGCCAATAAACAAACAA
GATCTGTTTATGATTCTAGTATTGACCATCTGTTACAAAAAAGAAATGCAGGCACATCGTAT
AAAATATGGATAATAGAAGAAAGCAAAAAGAAGACATGAGACTCATCCAGAGTTAATCAA
TATAGACATATAAGTCTATGTATATACAGGAATTGGAATCCCACTATACATTGTGTTATAACC
TTAAAAACCCATATAAAACATGGCAATTTTTCCATGCCTTTAAATATGCTTCTATCTAACATG
ATTATCAATAGCATGGCTGCTGCATAATTTTTTCAACCAATCTCTTATTTTTAGATATATAATT
TGTTCACAAAATTTATTATAAATAAGGCTGCAAGGAATTCTTGTATAAGAATATGTTTCTACA
TCTCTTATGATTTTCACTTTCATTATTTCTTTTTCTCTTATTCCGAAATGTGATATTGATTGGTC
AAAGCTATACATACTTTGAAGCTTTAGATACATATTGCAGGCATATTGCCAAGCTTGGTTTT
ATTAACATATATGCCCACAAGCAGAACGTGAGAATTTCTGTTTCTCGACACCCCAGCTGAC
CCAGGTGTTTAAAAGCAAAAGGAACCCAAATACTTTGCAAAGGAAGGTGTGATGCTTTCT
TTTTCTATTTTGACCTACTTATTACATTAGAAATAACATTTTACTTTATTTCTTGGCCATATTTG
TGTGTGTGTGTGTATGTGTGCGTGTGTGTGTGTGACAGAAAGAGAGACAGTGAGAGTGA
GAGAGAGAGGCTGCTCATGTTCTGGGCCAGGTTTTTATTAAGAGATTCATCTTTTTCTTAGT
CTTTTAAAGATGACTTCACAGAGTGATATAGCAAGGATATTAACCCTCTTCATGGCAATCAG
ACATTGCACAATATTTTCCTGGTAGCCATTAACATATTTTACTTGAAAGTTTTCTTTTATGATG
TCAGATTTTGGCTTTTCTTCAACTTGATATTACAAAATATTCATATTTATGTTTCCTTCTGGCA
CTTACATATATATTAAATATTTTATTATATGTATTTAATATGTATTTGGAATTCATAGTACAAGGA
GTGGGTGATGGATTTAATTTTCTCTGAAGTAAACCCATTCATTCTAACTTCTAATGCCCAAC
TTCAACATAAACATAAAAGCTAGGCACACTGTGAAATTTTTCTGAGTTTTAGACATCTTGTC
CATAAAACACTTAAATCATCTCATGAGATTTTTTTTTTTGGTGAAGACTAAATGAGATCCATT
CTGAGCAGGAACGTGTAGCCATAAAGCACCAGTCAAGAGAGGAGCTTCCTTTACCACAG
GAGCCTGGCAGCATTCCCAACTTGCCTCTAGATGGTAAGGAGGAAATTTCAGGTATCAGG
TCGATGGTGAATTTGCACTCTCCAATGCGACTAACCCAACATACCTAATTCCTTTTTAAAAA
CCCTTTATAGTTCTCTCACTTTGATATTTTCACCCCTAATAGTTAAGTCATTCCAGACAATTT
ATTGAATAAGGAATTATTCTCTTATTGGCTTGAAATGTCAGTTCTTTATAGACTTTTGATTTTG
CTAGGCTTATTCTTACTGTCTTATTTTTTGTTTATTCTTAAAGCAATGCTATATTGGCCTATAG
TTCAATGCCCTGTAGTATTTGAATCTTTTTACTGCATGAACCTACCCATCATTCCCCTTTCCA
GCACTTTCTCTGTGGGTCTTTCCCACAGTTGTTGTACAGAACATGGAGACCTGCCATTGTG
GGCTGCCTGACCTGATCCATGGCAAACACTGGCCCCATAACCGTTCCAGGTTCATTAAAA
AAAAATAAAATGTGCACAGTTTAACCTCCCCATTGTCCACCAGACAGCTGAGGAAATGCT

AACCTTGATCTCATAATATGAAATTTTGCAATGTAAAATAAAGCTTTCAGTTTCTTGCAAAC
ATAGTAGGTGTCAGGATGGGTATCTGGTGGAATAATATGTCTATAAAACAAAGAGCCTCAA
TACCCATGTTCACAGCAGCATTATTCAAACAGTCAAAAGATGGAAGCAACCCCAGTGTCT
GTAGATAGGTGAATGGACAAACAAAATGTAGTGTCAACATACAATGGAATATTATTCAGCC
ATGAAGAAGAAGGACATTCTTAACACGTGCTACAGCATGGATAAACCTTCGGGACATTTT
GCTAGGTGAAATAAGCCAGACACAAAAGGACAGATACTACATGATTCCACCTGTATGAGA
GCCCCAGAGCAGTCAAATTCATAGAGACATCAAGTGGACTGTTGCCTGCCAGGGCCAGA
GGCTGGGAAGGAAGGGAGAGCAGAGAGTTACTGTCTAATGGGTAGAGTTTCAGCTTTGC
AAGATAAAAGAGTTCTGTGTGTGGACGGTGGTGATGGCCACACACCCATGTGGATGTATA
ATGCCACTGACATTGTAACTTACAATGGTTAAGATGGAAAATCTTGTATTACGGGCATTTCA
TCACAATCCATCCATAAAATGAAAATAAAACAAATACACAAAAATCCAAAGCCCTGTAGTT
GATGATTTTGAAGATTCACGTCAGCTCTATTTTTTTTTTTTTTTTGTTCTGAGGGCAGAAAA
CAAACAGGCATCCACTGCACAGTTGTGGAGAAACAAAGAAGCAGGAAATGGTGATCACT
TCTTACTCTAAAGAACAGAAAGTCCAAGCGATCAGGAACGGGCGCAGCCCAGGGGAGCT
GGGTGAGTGTGAGCTTGCGTCTCAGAGTCCCCCATCAGTGCTGGCCTGAGGCAGGGCGC
CACGCTGACCTGCGCCTGGTTCCCTGCCAGGCTTCCCCACAGGCTGCCTCACTGGGGAA
TGGGACAGGGACCAGGCCACCAGTATTCAAGGAAACATGGACAGACTTCCTCTCCATGG
CAGTGCCCAGCTGTGTGTGTGACTCCATGTGGCAGTGTTCATGCCCTCAGCTGGGGGTG
GATTCAGCCCTGGTGCTTCAGGAGGGCCGGGTAGCCTCTGGAGACTCAGAAGCATGTAG
GAGTCTGGCCAGTGTCAGCAGGGCAGGCTAGGAGAGGCCCTGGTGCTGCCAGCCCCAT
GAACAGCAGCTCTTCTCAGGACACAAGTGCTCTGCTGTCCCTGCAGGGGATCTGCCTCC
GGTGGCCTGAACTGCCAACCAGCTGGGCTGCCACCCGCCACAAAAAAACGATCTCTATG
TCCGTGGTAAATACATGCTTTTCTAGTGGTCAGATATTTTGCTTACTACTTTTTGTTTCTGAT
TAATGTGACTTACTAAACATGGCTTCTCTCCCCTGCCCCAGTGCTGTAGAGCTGTCCCCCA
TAGGAGCTGGAGGCAGAGATAGTGTGTATGCGACACTTACAGCTGCCCAGGTGGTTCTG
GAAGTCCATCGACATGTTGCTGAGAAAGTCACTGCTGACTATGTCCCGCCACTGGATGCT
CTCCACATTGCACAGGGCAGGGTTGTTGCTGAACCGCACGGCGCCATGCAGGATTTCTG
CGGGAGAATGGAACTGCGGTGAGCACTCTTTCCAGCTCCTTTAAATTCCCATGAAGGATG
CATGTGCTCTTCGCATTTATGTTCTGCTAAAGAGATGCCATGGTGCGAACAAGAGTCCCGC
AATTAGCCCAGTGAACTTCCCGGGGGGCGGCCAGATAAAATTGTTAATGATGGTGCCATG
GAGGAGCTAGGGTTTTCTCTGAATTGGGCTTTGAGCAATGATTAGAAATTTAGAAATATGG
GGAGGAAGGAAGAGAATGGTTTAGTCAGTGGTCACAGCATAAATGATATGCATGTGGTAA
GCTGGGTCCCAAGGGGCTGATTGGCTGGACTGGAATGGTCGCATGATGTAGGGGGTAGG
CAGGGCAGGGAGGCAGGTCCTGAGTGCTACAGCCCTGGATGCCAGGTGAGGGGCTGGC
ATCCAGGGCTCCACCTTGCATGCCCAGGCGGAGCTTTTCATTAGAGAAAAGGATATGAGA
GTGACACGAAACAAGAAGAATCTGAATGTATGACAGAGACAGGTAGACTCCAGGGATGC
ACAGGATGGTGAAACTGTACAGCTGTTTCCAAAGAAGAACTAACAAGATGTGGAAAAGTC
AGGGATGAACACATTCAAGTCTTTACAAAGTGGGAGTGTTCCACTGATGACATTAAACATG
AACATTGACAGAATGACTTATGTGTACAAAACCCTGTGCAAGTGCTGTGGGCGCTGCGAG
GATGAATGAGATCCACTGGCTTCGATCTGGGAGCTGACAGGCTTGGGGACAGACATGTC
ACACACTCTAACCACCTCATGCCCAGGGAGACCCTGACACAGTGAGGCTGGGTACCTTG
CTGGCTGATTTATAAGCACTATCTTCATTCATTCTCACAATGGTGCAATGAGCATCTGTGAC
TTTAATAGAGAAAAAATTGACCTCTGATTCATGAAGCAATGCCCAGCTCCCCAGTCAGGG
AATGGGCATGTTGTGTGTACAGAGTACCAACGTGGGAAAGGAGGGATTCACCCCTGAAG
ATCCTGGAGGCTGCTAGGAAAATGAGCTTCCCCAGAAGGGCACAGGCACACATCTTAGA
GGAGTGTGGAAGTGAGGGCTGAAGGCTTCCTGCAGACAGGAGACCCAGTGCAGGGAAA
CAGTGGAACAGGCTTCTTCAGTGGATGCACAGAAGCAAGGAACTGAGGAAGGAAGCTTG

GGGTCATAAGGGCTGAATCGTGGTGCTTTGTTTATCAGGTTGGAGATGACAGCAGAACCA
TCCGTGAAAACACTCATGAAAAAGCCATGCACAGGTGGATGGAACTCAGGAGAGGACAG
GGCTGGAAACACCAACAGGGATGTCCAGGCTGCTGTGGCAGCTCGGACAACAGACCAA
AGACCTGAGAGAGAGCAGGGGGCGGGGCGAGGAGAGCACAGAGGCACAGTCAGAGCT
GGGCCTTCAGCACACCACAGGCAATGACTGCCAGAGGAGGGCCAGCACACGTGAGGGA
GGCTGCAGGAGCCGCTTCCCGGTGGCTTAGGGGAGGCTCACCCTGCAGCAGGCACTAG
GTTAGCTATTAATGGTATCACATTAACTAATCTGCATCTATACAATCCCAGCCTTGGTTTAAA
AAACAATCATACCAAGGAAGTGGAAGGAGCTGCTGAGGCCTGTTCCTACAGTCACTTAAT
TTTCTGCACCACCTAGGAAGTATGTCACCATTTTTTAGCAGCTATGCCATCCACATTTTTGA
AGATGACGAAACAGAGACACGGAGGGATTGGAGAATGTGCTCCCGAGCCATCTGGCTGC
TGAGGGACGGGGCCAGGCAGGAACCCAGAGGGTCTGGCTCGGAGGTCCAGGCACCCG
CACAGGCTGCCTCCCTCCGAGACAGCGTGTGAGGCGCACTGTAGGAACCGGTGTACCAG
GCCATCTCACAGGAGGCACGCACAGCACAAGCATGTGCGCGGCTGCTCTGAGCAGCAA
GGGGTGTGGGGGAACCACACAATTTTATAGGCGGTGAACTGAGGAGCCATGACCAGAGG
TGCTGGCTTGGGTTCCAGCGCTGCACCCCTGTTGGGCTCCTGTCACTCCGGATATGACTC
CCTCTGCTGGGCACCCCTTTCATCACACCCCTTTCCTACCAGCACTGCCTCCCTGTCAGA
ACTGCATGGCACAGGAGGGGCGTTGAGCGCCCTGGGGCAGAGAGTGGTGGCCATAGCT
GGCCTCTCCATTCACTAGCTGGGGCCCTCAAGGGAGTTACTTGACTTCCATTTATTTATTTG
TACGATGGCAGAGCACAGCCAGTCCCCGGCTTAGCATGATTTAACACAGATTTTCGACTTT
GCCATGGTGTGAAGGCAATATACATTCAATTAGCCCTGAAATCCGACCTAATGACGCCTTC
ACCCTCGGCGGGGTCCAGCCAGATAGGATGCCATCATAAGTCAAGGGGCATCTGTACAGT
GCCTACTTTGAAGAAAGGTTGTGGGCGTTAAATGAAGTTACCAATTTAACATTCCTGGCCC
CAAACCCACTGCACAGTGAGGCACAATCACTGCTATCTCCAGTTTTTTAAAATCAATTTCC
CTTCTTTTCTATCATCCTTCATTGTTTCCTCTCACAGGAACCTCACGAGTCCTGCTCACAAA
AATTCAAAGTTCTAGAACACCTCATTCTCTATCCTTAACTTACACAGGGCTTTCTGTGTAGC
AGCGCGACTCTCGAAACTGTCCTCAATGTGGGTTGATTTTTACTGTATGTCTTTTATCCTTG
AGGTCCAGGTAATAAGACTGGCAGTTTATTCACTGCCTACACACATAGATTCACGATTCCT
GCTCAGCTTGTTTGGTCATACTCTCTTTTACTCAGAATCTTTACGATTTCTTACTATCTCTCA
GTATTTTTTCATGTCCATGTAAACACTGGTACCCCGCTTAAATCTTCCCATTCCAGTATCTG
CCATCTTCTCAGTAACTTCCCCTGCAGTATCTTACACACAGCCGGCACTCAGTGCACGTGT
ACTGGGTAAAAATAAAAGCCTTCTCCGAGGTGGAATTGAGTGACAAGCTCGCTGGGAGC
CATCGGAACTGCTGTCTGCATTTATGAACCCCCAGCCTTGGCATCCCAGCCTCTCACCCT
GTAAATTTCTCATGGGCAGCTCCTTCAGTCCGGTTTTATTTGCATCATAGTTAGATAAGACT
GCTAAGGCATAGGAATTTTCGTAGTACATATTTCCTCTGATGATCTGCAGGTTTTCCAAAGG
AATTCGCTCCACTGTGTTGAGGGCAATGAGGACATAACCAGCCACCTCCTGGATGGTCTA
AGAAGAGAAAACATAAATGCGTGATTTCTCTTTTGAGGAACTCAAGAACTCAAGGTCTAA
AATGGGTCCAGGGAGCCCTAGGACGCCCAGTTGAACCCTAATGCACACGAGTGCTCAAC
AACTGTAAAATAGGTAGACGATTGCAGTCATGGACACAGGAGGCAGCATGTCCACGATCT
GCAACAATCCCTGCACGTTAGCCTTTGACCCCTGTGATAATGAGCAGTAAGCATTGTTCTG
GAACACTGTGCTCTCATGCCTGACGGGGTTCAACACTCTCCAGTGAAAATGGAAATTAAA
AAGAATTACTTTGTATCTGTCTTGCATTATTTCTTATTTAAAAAGGAAAGGTAAAAGAAAAA
CAGAAGAAGGCTAAGTAAACCCCATCCAAAATTCAGTGAAACTCTGAAGCCTTCAGAGCT
GGCTCCTATGGGGTCATTAGCGCCTATCGTGAAATAAGAATTCTGTGACCAGCTGAGTGA
AACTTGGTACCTTGAAATCAGCTATAGTAGGAGTATTTACACTAAGGGAATAGGCAAATGG
TGCAAAGCAGGGCTTTTAAAAAAAGTTTTTAAGAGCGAGTATGGCAGCACATCATGACTG
AGGTAATGGTCAGGGATAAACGTCAGTGTCTATGATGACATGGAACTCCAGATTAGCCTGT
TTCTATTTGATATCCTTTTCTAGTTATTACTTGACATTAGCTGGTAAAATGGCTTTCTCATAAA

ACTCTCGTTTGTAGCTCTGTAAGACTTGTCATTGCCATAGCAAAAATAAACACACTTCAAG
TGGAATTCTGCCCAGGCCTTTCTCCACTTAGATTTTCTCCAATTTTATTTGTGTAGGAAAAT
CAAAGTCACCAACCTTTAAGAAGGAAAGATCATAATTCCTCTGCACATAGGTAATTTCCAA
ATTCCCAAGGACCACCTCACAGTTATTGAACATCCTCTGGAGGCTGAGAAAATGATCTTC
AAAAGTGCCCAACTGCGTGAGCTTGTTACTCGTGCCTTGGCAAACTGGAAGAAAAAGAC
AGATATCACATGAAATACTGAGAAATGCAGAAAAACTAAAATAAAACAATCCCTCAAGGTC
CAGGTATTAAGGGTAGCAGTTTCTTCACTCTCTCCACACAGAGACTCACCCATTCCTGCCC
AGGTCATCTGGGCGCAAAGCTTACTAACTTAGTGAGCTTTAGTGACTGATAATACAAATTC
TCACTAAAGGACACAGCAAAGAAATTTGTGATAAAAGATCTTTAAGTTTATATAAGGTGCA
ATTCATAATAAGAGATAGCTTGTCCCTATAATGCATTGAATTGCATTAAAATGCCAAAATAAA
AAAATGTTGAAATTTAAAGGGGTGGGTGGTAGATGAAAATTCTATGAAAATCTTGAAGACA
GGATTAACCTGTAAGAGCTCCAGTAGGCAAAAAATAAATTTCACAAGGTATTTCACGAATA
TAACTGTAGGGTTGAGTTAATAACTATATAATGAACTTTTTACTTCTCAAAAATATTGTGAGT
CACATTGTTGTGCGTGCAATGAATGTTTACAAACATAACATACACTGGGCAGTAGTGTACA
CAACACTACACCTAAAATCAGGGACAAAAGTGTAAATTAAAAACCTAACTGGCTGGAGAT
TAAAAATATTATTCAAATGAATGCATCAAAGAATAAATCAAAGGTCTAACTGTAGAATATTC
AGAAAATAACAAACATCAAAATACAACATTTAAAAACACTCTCAGCTAGTGAGAAACCCA
CACATCCCTATGCTCGTCCACACTCCATGTTGATCCACCCAGCACAGGCGGAGGCACAGT
TACTTCCCTCACCACCAGCATACAATCTTGTATGCTTGAGTTCCTAATGTTTCAAGGCATCA
AACTTAATCCAAGAAAAAATTTTAAATTAAGGGACAAAGTTGGACCATAGAAATGGACGTG
GTGATCCATTTTTAAAAGCAAAGGAGAGGTTTATAATAAAGTTGTATTTACAGAACTAGTTC
TTTGAAAGGCATGTTTAGGCAAGCATAACAAAAGAGAGTCTTGGGCCTTTTGAGGCAGAG
AGTGTGTAAAGGGACTATAATTTGAACCCAAATTATCTTAAGAGTGAAGAAATATGGCCTC
TGGCTTTGGATGTTGGATTGAGCCTTGGCCAGGTTCTTCCTCATCTGCAAAGGCACTTCTG
CAATCACAGTAACCAGGTGGTAGAGCACAAACCATGGCCAAAATCAATACACTGGCACTT
TCTTCAATGCTTTTTACATGTGCATTTCTATTTTACAGGCTTCTTTGTTCCTTCCCCATGGGA
ATGATTCTAAAGAGTATCATTGGACTGTCTGCCCTCTCAATGAGCCATGGAAATGAGCACA
TGTTCAGACAGCACATAACCAACCCAAAGCACTTACACAAACATTAAACATGAAGCAATC
TTCACATCCTCTTTAACTCTGTGGGGATTTCCCTGTTCTTCCCTTCTGTGTTCTCATTCAATT
TCAGTGATAATGATAGCTTTCACATGGGAACCATGACAGTTGACAAGATGGAGTCCCTCTC
TCCTATCACCAGGGGCAGAATTATTCCATCATTTTTCCAACCATAACTCTGGATTGACCCAG
CGTCTCTGGTCTGCTTTTCCTCAGGAGTATGGGGGAAATGAGGTTGCTCTGAATGCAACC
TCAAGTTGGAACAATTGTGTGGTTGTCTGCTACACATGCAGATAGGGCTAAACTGATTTCC
CAATTCTAAGGTTCTTAAAGATGTCAGCACTGATGTAGGTACCAGGTTGTGACAATAACAT
GTGGCCTTTCGGTTGGCACACCTTGGAATAAAAGCTGCTTTCCCAGCTGACAGACTATGG
CCTCTAACTCTCATTCCAACTAGATTGGCTGTCGTCCAGGTAGCCGCAAGTGTTAGTTGGG
CCCCTCGTGCCAAACAGATGAACAGGGTCAGCTAGAACCTAGGGCTGGAACACGTTGCA
GAAGAGTTTCCTCAGGATGCAGAGAGATGAAAACAGGCAGCTGCTTTCATTTTAAATAATC
AAGGACGTCAGCTTGCAAGTTTCTCTCAGCTGAACCAAAGAATTTTTATTATTCAATCACA
GAGTAAGTAACTTTTGTCAGTAAAATAAAGGAAAACATGCCATTTAAGATTAATCTATTATA
GAAAACAGTATTTTAAAAAAACTGTACTTGTGCTTTTATTTATTTATTCAAAAAGTGCTAAAT
AAGCCCCAAATGACCCCCCAAAAGCCCCTTCAGGGCTTATTTAGCACTTTTTGAATAAATG
AATAAAAGCACAAGTATTAGAAGCACTTCATTATCTATGAGGATAATGACCCTTCATTGTCT
ACAATTAGCATGTATAACTTTTATATTCAGAAAAAGATCCTGTATTTGGGAGACAGTAACAA
CATTACATATTCTTGATGTAAATCTTAATTCACAAAACATTTTGGGAAGCAACTGGACAATT
TGTTTAAACAAGTAAGTATGCTTAAATTCTACGTCATGATGATTCCACTTTGGGAAACGTAT
TCTGAGGAAACAGACCTTATGCAGAATGTTGCTCATTAATTTTGTTAATAATGGAAAAAATT

GCAAAGAAACTTAAATGCCCAATTACCCAACATGTCTATACTTGAGAATAAGAAGGCAGTT
GATATGCTATTTGATTCAATATAAGGCATGTCAATCTTGAATACTATCCTGGGTTAATTACTA
AAGTATCAAAAGAAAACACATAAGGTTGTTTCTTCTGAGGCTTTCCCCTGTCACATACCCA
AAAGGAGACAGCAGCATTTAACTGAAATCTAGGATGAGGATCACAATGACTTGGTCTAGG
TTTTCTAATTAAAACAAGGTATGATCTTGGTTCCATCCTTATACCAAATATCTCATGTTAGAA
ATATAAAGCAATGAGAAAATTTAATGTGAGACCACTTATATGAGTAAATTACTTAAAAATTAC
AGTCACAAATTAGGTAGATTTAAGACAAAACTGCAAAGAGCAAGAGGTTTTTAATTAATTT
AAGATAAGGTTTTTGTTTAGCCACTTCCTCATTGGGTGACCTTGAAGTACTTCTCCAAGCT
CTCTGTGCCATGCTTTCCTCAAATATGAAAATGGTGACAATAACGTTGATCTCTGCTGTGG
TTTGAATGTGTTTTCCAAAGTGCGTTTATTGGAAACTTAATCACCAATGCAACAGTGTTGA
GAGGTGGAAACTTTAAGAGGTATTATAATTAGGTCATGAGAGCTGTGCCCTAAGGAATGGA
TTAATGCCATTCATGTAAGAATGGGTTAGCTATCACGGGAGTGAGTCCCTGATAAAAGGAT
GAGTTAAACCCGCTTTTTCTCTTCCCCACCACCTCACATGCATGCTCTCTTGACCTTCCAC
CATAGATTGACACAGCATGAAGGCCCTCACCAGCTGCCAAGCAGATGCCAATGCCATGCT
CTTGGACTTCCCAGCCACCAGAACCATGAGCCAAATACATTTATTTCTTTATAAATTACCC
AGTCTGTGATATGGGTATAGCAACATAAAATTAACTAAGACAACCTCCCAGAAATGTTTTG
AAGATTGAAAAAATATATAAGCTTTTGACTTTCTTTTATTAAAGTTAACTATTGAGAATGCAA
CAGAAACATAACATAACATGCATCTGGACCCACGTTAGGGGATGTCAGCCACTCTATATTG
TCTTAAGTTATTGTCTAAACTGAGATGTTTGGATCTGCATATGGACTTACTCAAATGTTGGC
CATTAAACACAGTTACTTTATAGAATATTTATCAAACCAGAAAAGAAAGAAGAAAGAAAAA
TGAATACATAGTACTTTAAGGATTACATAAAAATATTTATTTAAGACTAGGTATTTCAGTAGTT
GATTCAAAGAAAGAGTTGATATTACTTTAAGTCTATACATTTGAAAACCAGACAATTAAAAG
TATTTGCATCAAGTTCCATACAAAATGCTAGTGGTTTGCCTTACCCTTTGCAAATCTGAAGA
CAAATTCTACTGCATCAAGAGCAGCAGGCCCTATGACTTAGGACTTCAAAGCACTAACTTT
TTGTCCTAAAATTTTATTATGATTTATAAATGTATAGAAAAGCTGAAAGAATTGTACCGCAAA
CATCCTTATGTTAGCATTTTACTACATTCACCTTATCACATATCTATTTTTCTGTCTGTCAGAT
ATGGTCCTCTAGTCTGCCTTTGGATGCATTTTAAAGTAAAGTAGAGATATCACTATACTTCA
CATTGGTATATCACTAACCAGAGTTCAATATTAGTTTGCAGTCATCATTGTTGTTTTTAAGGT
AACATCCACAAACTGTGAGAGGATTAATCTTAAGTGTAGCATTTGATGAGCTTTAACAAAT
GCACATGCTGGAGTAATCTGAATTCCTGTCATGATCGAAACCCTAACTGTCATCCCTCATG
CCCCTCCCAGTCGATCCCTGCCCCCATACACCAGAGGCAACTGCTGTCCTGATCCATTTT
CACTACAGAATAGCTTTGCTTGCTTTAGAACATAGATGGACTCACACCACACTGTCCTCTT
GAGCAAGCCTTATGTCACAGCACAGTAGTTTTGAGGTTCATCTCTGTTGTTGTGCTCCACT
TGGCTGTCAGGGGCAGGTTCAGGAAAGTGGATATACGAAGTTATCTGTGCATTCTTGTTG
GACATCTAGGGCGGGCCGGTGGTCTCGTTATTATTTTTCTTGAGAAACTCCTCACTTTATTT
TAACTGAAAATAAAAGCAATCTGCCTCCTTTAAGTTTTTAAATGCCTCTGGTTTTGTAGGGT
AGGCCCTGTGGCCTTCCTCTGCTATTATGGGGTGTATACCACTCTGCCCACAAGCCTGGG
CAAATCTATATTACTTATTAAGACACAGTATTAGAACAAGTTTCTCCAGATTTGCCCAAATC
CACTTAGACACTTAGAATCGTCTAAGTAAACTATAATTGTTTTAATTTGTTGTTTTCTTAGCT
ATGCATTAAAGGCTGCTGAGGTATGAGCTGGGCTGTTGTCAGTCTCTCCCCTTGGCCTCC
AAGTTGTCTCTGCTTCCATCCCCAGCCCAGATCTTCGCATCTTTGCTTCCTTTGCATCTTCG
AGGTACGCTTAAAGGCCTACCCTCTAGAAAAGCCTGTTCTGCAACTACATTAGCTTAAGTA
CCCACCATTACATCTGTCACATTATCCTGCATTCTCTGCTCCCTGGCAGTGATCACTACTTA
TTATTAAGTAGGAATAACTTAATAATAATCATGAAATTATTATTTGCCTATGAATTTGTTATCC
ATCTTTCCCTACCAGATATACAGATGGTCCTCAACTTTCCATAGGGTTATGTCCTGATAAAC
CCACCCTAAATTGAAAATAGCAGTCGAAAATGGCATTTTGTAGACATGATAGGATTCAAAA
ACACAAAACACAATATCCCCAAAATGCTGGTGACACAGTCCACTGTATAGTATCAGCTGTT

**TAGCCTCTTGATTGCATGGCTGACTGGGAGCTCTGGCTCCCAGCCATTGCCTGGCATCGC AAGAAAGTAACATATCAGACATCAACTAACCCAAGAAAAGATAAAAAATTCAACATTAGAA GTATGGTTTCCATTGAATGTGCATTGCTTTCACACCATCGTAAAGTCAAAAAATCTTAAGTC AAACTGTAGTAAGCTGGGGAGTGTCTGTAAGGATGTGTGTGTGTGTGTGTACACATATGTG TATATATAGATAGCTTATTGGACCTACTGGTATGAAATGCACACACACATGTTGGATTTATAT TTTTGTTGTGAAAGCAATGGATGGCCATGTGACCAAAGTCTTGGATGACTTGTCCTGGGTT TCTCTGCCTTTCTCGTTTCCGTGTGCCTGAGGGAGATAGAGGAGGAGGTGGGCTGTATGG GAAGAGTGACAGTCCTTGCCACTTTCTGGTTGGTCATTATAAAATATTTCTAGCTTAGTCAT ATATTTAATTAGGCTTTAAAGGCCAGAGGAGGATTTTTTTCCCGCTTTCTTGCTGAAAGATT TAAGAAAATCTCAAATAAGATGAATGGATAGTTCATTTATAGGATGAACTTAAAAATATGTA AGGAAAATATAGCGAGTTTGCCAGAGGTTTCATTGAATCAGGACTGGTCATTTCTTTTGCT ATACTAAAATCTCAAAGTGAACATGTATACACATCTCAAATGTGCTCAAATGCACGGAAGC ATGACAGGAGGAAGATACACCGTGTTGTTAACAATGACCTTTTCTGAATGGTAAATTTCTG GGTGTTTTTCTTATTTACTCTTCATAGTTTTTGTGTTTTTATGTTTTCCATAGTGACCATGTATT ACTTTATGCTAAAAAATTACCAAAAATTTAAAAAAATCTGTAAAAAATTTACAAAAATCAGT GTTAAAGAAGGCTAAAGAAGGCTATCTTAAAAAATACATTCCTTAAAAATTGAGAAATAATC ATGTACATCTATGAAGACACTCAATTTACCACATAAAAAAATTTTATAAATTGTGACTATAGA ACTAGACTAGTCTATGAAATTGATTTAAACCACACAATAGTTTGTAATAATGGAAACTAGAA AACCTATAGTTGTGAGTCATAATGAAATAAGTGGTTAAACCAAAGTAAATACACTTATTTAA ATTTTCATTGTGGGTTTAATCAAAACGTTTGTGGGATGGAAAAAGTCATAATTTTGAAGACA ACTTTTATTCACAAAGCAAATTTTATGTTTTTTCCTCAGTTGTTCCTTATTTTACCCTATTTTAT TCCACAACTAATAAATAGTTCTTAACTGATAGGCATCCAAAGATACCTGTCAAACTGAATGT GAATCTATATTGGAACTTGAAAATACACATGTGATACTCAAAAAGTCCTTTTGAATTTCTGA CAATCTTCCATGTACATTATAATATGCCTTCTGGACTAAACATGCCAAGCTGCGGCATGCCC AGCACAGCTGCAAACTCATGTATCTCTAATTAAGTTAACAAGCAGAAGACTTGAAAATAAC ATCTCTTTATTTTGAGTTAGGGAACAGAATACCCAAACTGGTGGACTTCCAACCCCATGAT TCTCCTTCTAGCCATTAAAAGAATCCAAAGCCAAGCTTGTGATTTGTAAAATATACACTAAG TAGAGTGAGAATACCTCTTGGGAGTTCCCACCCTCTTGGGAGTTCACTGATGCTTCTGATC ACAGGAAGGCCAGTTCACAAGGGACTGTGGAATATGCTTCCAGTGACATCATCAAACTTC TACAGTGTGAAGATTGTGATGGTACCATTATGGGATGATCTCTAGCCAGTTATCTCAACATG CTGGGTCTCAGTGCTCAGTGTACTCTGATTGACATGGGGCCAGCTCGGCGAATTTCACAG GGCAGTTGTGAGCAGTACTGAGATCATGGACATGAAAGGGCCTCAGAAATCATAGATCTC TCTAAATATGGAGCATGACACTGAGGCTTCACATGGCAGCAACTATTGCAATCTATTTCTG CCTATGTGCCTTCAATAATAACTGTTTTACCTTCATTGGTTTTTAAACTAATAATCTTAAA**CTG AGGATGTCCAATTGCAA

>A23_P_1.1kb (1108bp) CG11.1.1_ chr12:68898148-68898149

CGGGCATGAGCCACTGTGCC**TGGCCTGGTGTTCTTTTTTTTTTTTTTTTTTTTTTTTTTTTTT TTTTGATTTGAATAGTTCTATTTTTTATTATTGTTATTAGAGACAGGACCTTCCTCTGTTTGGAG TGCAGTGACACAACCATAGCTCACTGCAGCCTTAAACTCCTGGGCTCAAGGGGTCCTCCT TCTTCAGTCTCTCAAGTATCTAGTACTACAAATATATGCCACCACACCCAGATAATTTTTATT GTTTGTAGAGATAGGGTCTCACGGTATTGTTCAGGCTGGTCTCGAACTTCTGGCTTCAAGC AATCCTCTCATCTCAGCCTCCCAAAGCACTGAGACTATAGGCATCAGCCATCATGCCTGG CCTGTTCTCTTATTTTTGTTTTTTTTTTTTTTTTTTTTTTTTTTTTTTTTATTATACTCTAAGTTTTAG GGTACATGTGCACATTGTGCAGGTTAGTTACATATGTATACATGTGCCATGCTGGTGCGCT**

**GCACCCACTAATGTGTCATCTAGCATTAGGTATATCTCCCAATGCTATCCCTCCCCCCTCCC
CCGACCCCACCACAGTCCCCAGAGTGTGATATTCCCCTTCCTGTGTCCATGTGATCTCATT
GTTCAATTCCCACCTATGAGTGAGAATATGCGGTGTTTGGTTTTTTGTTCTTGCGATAGTTT
ACTGAGAATGATGGTTTCCAATTTCATCCATGTCCCTACAAAGGATATGAACTCATCATTTT
TTATGGCTGCATAGTATTCCATGGTGTATATGTGCCACATTTTCTTAATCCAGTCTATCATTG
TTGGACATTTGGGTTGGTTCCAAGTCTTTGCTATTGTGAATAGTGCCGCAATAAACATACGT
GTGCATGTGTCTTTATAGCAGCATGATTTATACTCATTTGGGTATATACCCAGTAATGGGATG
GCTGGGTCAAATGGTATTTCTAGTTCTAGATCCCTGAGGAATCGCCACACTGACTTCCACA
ATGGTTGAACTAGTTTACAGTCCCACCAACAGTGTAAAAGTGTTCCTATTTCTCCGCATCC
TCTCCAGCACCTGTTGTTTCCTGACTTTTTAATGATTGCCATTCTAACTGGTGTGAGATGAT
ATCTCATAG**TGGCCTGGTGTTCTTATTTG

A23_R1_1.7kb_CG1.2_chr7:55636272-54753591

GACTTGAACC**AGCATATAATGAAAAGTGACCAGGTGCCAGGCATTTTGTAGGAGCTTTTAA
AAGTGTAATCCCATTTAATCATTTAGATAATCACAACTCCTCTATGATGCTACACTTGAGAA
AACTGAAGTTCAAAGTATGAAGCAATGTGATGCATGACAGTTATTAAATGGTAGACATGTA
CAGACAAACTCCACTTCTGTATAGCTCTGTCCCTCCCCCTTATATTGTTGTCACAAGTTACA
TCTTCATACAAGTGATCCTCCCACCTCAGACTCCCAAATAGCTGGGACTACAGGTATGCAC
CATCATGCCAAGCTAAATTTTAAAATTATTTCCCTTTCAATTCAAGGGGGGATGGGGTATGT
TTTAGAGACACGGTCTTGGCCTTCGTTGGAAACGGGATTTCTTCATTTCATGCTAGACAGA
AGAATTCTCAGTAACTTCTTTGTGCTGTGTGTATTCAACTCACAGAGTGGAACGTCCCTTT
ACACAGAGCAGATTTATTGCATTTCCTGTCTGGAATGAAGATTATGAAAAGGCCAACAACT
GTGAGCTACAGTGGTTTAGAAGAAAACTTAATTTCTAATTTTTATTTCTCTGGTTTTTTGTTC
TTGCGATAGTTTACTGAGAATGATGGTTTCCAATTTCATCCATGTCCCTACAAAGGACATGA
ACTCATCATTTTTTATGGCTGCATAGTATTCCATGGTGTATATGTGAGACATTTTCTTAATCC
AGTCTATCATTGTTGGACATTTGGGTTGGTTCCAAGTCTTTGCTATTGTGAATAATGCCGCA
ATAAACATACGTGTGCATGTGTCTTTATAGCAGCATGATTTATAGTCATTTGGGTATATACCC
AGTAATGGGATGGCTGGATCAAATGGTATTTCTAGTTCTAGATCCCTGAGGAATCGCCACA
CTGACTTCCACAATGGTTGAACTAGTTTACAGTCCCACCAACAGTGTAAGAGTGTTCCTAT
TTCTCCACATCCTCTCCAGCACCTGTTGTTTCCTGACTTTTTAATGATTGCCATTCTAACTG
GTGTGAGATGATATCTCATAGTGGTTTTGATTTGCATTTCTCTGATGGCCAGTGATGATGAG
CATTTTTTCATGTATTTTTTGGCTGCATAAATGTCTTCTTTTGAGAAGTGTCTGTTCATGTCC
TTCGCCCACTTTTTGATGGGGTTGTTTGTTTTTTTCTTGTAAATTTGTTTGAGTTCATTGTAG
ATTCTGGATATTAGCCCTTTGTCAGATAATGCTAGACAGAAGAATTCTCAGTAACTTCTTTT
TGTGGTGTGTATTCAACTCACAGAGTTGAACCTTGCTTTAGACAGAGCAGATTTGAAACTC
TCTTTTTGTGGAATTTGCAAGTCGAGATTTCAAGCGCTTTGAGGCCAACGGTAGAAAAGG
AAATATCTTCGTAGAAAAAATAGACGGAATCATTCTCAGAAACTGCTTTGGGATGTGTGCA
TTGAACTCACAGTGTTTAACACTTCTTTTCATAGAGCACTTTGGAAACACTCAGTTTGAAG
ATATTTCCTTTTTCACCATAGGCCTCAAACTGCTCAGAATTATCCCTCTGCAAATTGTACAA
AAATACTGTTTCCAATCCGCTCAATCAAAAGTAAGGTTCAAATCCGTGAGATGAATGCATA
CATCACAAAGAGGTTTCTCC**TCTGAGAGAT

A26_P_685bp_CG1.1_ chr4:57148028-57180989

AGCTACTGGG**GGGGAAAGAAAATTTTAATGCACAGGAAGCAACAGATATAGAGGCCCTGA
GGCAGGAATGAGCTTAGCATCTTTCAGGGATGCTAAAAAGGCCATTGGGAATGGAGCCCA
GAGACAAAGCCATCATCAGCAAATAAATAATTGGCTTTAAACATGAAAGTGTCTTCAGAAA
GCTAGCTTAAAAATGTGCAATTTTTCTTCCTGATTCAACTAGAACTGCGACGTAAATATGAC
GCATATTGCTACATGTTTGCGTGGGCACAAATCTTATTGGCAGTTCTGGTAAATAAACGTAG
GACCAGTTGGGGTCTCCCTGACGCCTCTCACCACTGTGAACACAGTCTCTGATAACATTG
TTTTGTCACTCTTCACTCCCACCAGCAATGAAGACAGATAAATGGCTTCCTGACAAATGGC
CCCACTTTCCAAGGTCTGGGGCTCTGAAGGAAGGAGACGTGAGCCTTGAAGCACTTCTG
GAGTGAACTCCTTGGAGGGTTTGTGGCCGTGAGCTCTCTGGGGCTGCTGCATAAGGAGC
CACTGCATTATTTCTTCTGAAAAATTAGATTTTGCTTGCAACCCATTAAAATTTCCAAAGCTT
CATGAAATGGATTCCAGACTATGTATCATTGTAATTTGCAACAGACAATGCCTGATCATAGA
AACAATAGAGATTTAAAGTGAAATAT**GGGTAATTT

A31_P_1183bp_CG2.1_ chr12:65905357-chr4:57086268

TATGTAAGAA**AGTGAAATAAACAGCATAAATAAAAACAATCACAACTTCTGGAAATCAAAG
ACACACTTAGAGAAATGAAAAATGCACTGGAAAGTCTCAGCAATAAAATTGAACAAGCAG
AAGAAAGAACTTCAGAGCTCAAGGACAAGGCTTTCGAATTAACCCAATCCATCATAGACA
AAGAAAAAAATAATTTTAAAAAATGAACAAAGCCTCCAGGAAGCTTGGGACTATGTTAAGC
ATCCGAACCTAAGATAATTGCTTTTCCTAAGGAGGAAGAGAAATCTAAGAGTTTGGAAAAC
ATATTTGAGGGAATAATTGAGGAAAACTTCCCCAGCCTGGCTACAGATCTAGACATCCAAA
TACCAGAAGCTCAAAGAACACCTGGGAAATTCATCACAAAAAGATAATTGCCTAGGCACA
TAGTCATCAGGCTATCTAAAGTCAAGACAAAGGAAAGAATCTTGAGAGCTGTGAAGCAAA
AGCATCAGGTAATGTATAAAGGAAAACCTATCAGAGTAACAGCAGATTTCTCAGCAGAAAT
CCTACAAGCTAGAAGGGATTGGGGTCCTATTTTTAGCCTCCATAAACAGAACAATTATCAG
CCAAGAATTTTGTATCCAGCAAAACTAAGCTTCATAAATGAAGGAAAGATACAGTCTTTTC
TAGACAAACAAATGCTGAGAGAATTCACCACTGCAAGCCAGCACTACAAGACTGCTGAA
AGGAGTTCTAAATCTTGAAACAAATTTTTGTGATACACCAAAATAGAACCTCCTTAAAGCA
TAAATGTCAAAGGACCTATATAACAATAACACGATGGGAAAAAAAAGACATCCAGGCAAC
AAATAGCATGATGGATAGAATAGTACCTGACATCTTAATACTAACATTGAATGAAAATGGCC
TAAATGCTCCACTTAAAAGATACAATATGGCAGAATGGATAAGAATTCACCAACCAAGTTT
CTGCTGTCTTCAGGAGACTCATCTAACACATAAGGACTCACATAAACTTAAGGTAAAGGG
GTTGACAAAGATATTCCATGCAAATGGACACCAAAAGTGAGCAGGAGTAGCTATTCTTATA
TCAGACAAAACAAACCTTAAAGCAACAGCAGTTAAAAAAAGAGGGACCTTATATAATGAT
AAAAGGACTAGTACAAAAGGAAAATATATAATAATAAAAGGAC**TAGAAATTGC

A31_P_15005bp_cG1.2.1_chr12:65663901-65666658

CCAGGATTGTGATTGAGCCT**GAACTTTTAAAATTCATTTTATCTAATGAATTTGGGCCTTTTT
AACCATGTGTCCCTCTTTTCTTCAACTTTTCTTTAACTTACAGGAATTTAATTTTGTTAAAAG
AAAATAATTATCCTTGATTATAATTTGAACATTGGAGATACTAAAAAATATATATATATATATTC
CACCTTGGCACTAACTTTTCTTTGTTCAGATATGGTAGAGATGGATGATTTAGAATAAATGA
GACAAGTGGCCAGTCTCTATGTGTGGCTTCTGAAGGAAAAATTTGTAGAATATATTATTTAA
AAACTAATAGTTCAAATGTAGGCAAAGATAGCCTTTATTTTATGTTATGTATTAAAAGTGTAT
ATTATAAAGTGTGGGGATGCCCATACCTGGGGACTAAAATTATAAAGACTGATTGAGGACA
TACTTGAAATACAAAATTATTTATTTAATGTTGACTCCTAAGTAGTTTAAACATTTAATTGTCA**

AATTTTAAAGCTGCAAAGAATCTTGTTATCTAGTTCAATGCTTTTATTTTATAGATAAGAAAC
CTGAGAAACAGGAAAATCTAGCTCAAATGAAGAAGCTAGTTGGGAGAAAGTATTTTAACA
GGGAGCTAGTGAATATTTTGTAGGCAAATCACAGGTCAGGAGAGTGGCAAATGTGACAGC
TAAGCCTTAGTGCAGGATACCTAGAGTAGGACAGGAATATTTTTGAGCCAGGTATAGAAGT
GTATCTTCTTTATTGAGGCAGATCAAAAGCCAGGAGAACTGAAGAGTGAAAATGAAGATT
CCTGGCCCAGACAGCTAATTTGTAGAAGTATTGAAATCTGGAGGTCATATCAAGCAAGAC
ACAGTTTCTGAAAAAGAATGATGAGTTCACTAGACTTAGAAACTTTTAAAATCAAATAATTT
TTAAATCACCATCTATTTAATACAAATTTGGAGCAAAAGAAAGAGTTGACTGTAATTTAAGA
AAATTCAAATTTGGAAAAATTAGCTCTGGTCACTTTGTGTAATTCTGTGTTACCATATTTTGT
TTTTGTTGGTTTTCCTTTAGTTTCCTATTAAAAGGCCTTTTGGAACAAATTTTAGCCTTTAAT
TTTGCTAACCTAAGAAGAAATGGACTTTTTAAAAAATTAAGAGATAAAGGACCTTTTATCTA
AAGATTACATATTCTGGTAGAGTAAAATGAGAGCAACTCTAGAACTTAGAGAAGACTCATG
TAGTTGGGTTGATTAAAAAGGAGAAAGTCTAAAGAAAAATGTTGGGTGAGCTGCAGTCAC
GTAGGGTGGTATAAAGGGCAAGGCTTTGGAGTTAAGACAGTCTTTTATTCAAACAGTGACT
CCACATTCAATTGTGATACAACTTCTCTGGATCATCAGTTTCCTCATCTGTGAAATAAGGAT
TATAATACTTGCCTCATAGAAAAGTTGTTGGGTTTAAAGAGACCCAATGTAGTATGGAACA
CCTGGTATACAGTTAGTTCTAAGTAAATACTAGCCTCTTTTCTCATGAGTGCAGGGGAGCG
TGCATGTATGCATGTTTGCTCCTCCAGTGTATCAGGGGCACTATCTATATCACCCTTCTCAG
ACTTTGGTGATTTTACTTGCTTATTTGTCTTCCTTTTCATAGAGCCAAGGTATTTCGTGTATTT
GGTAAAGGTCCAGAAATAAAAAGAAAAATTCTCAGTGATGGAACTCTGCATTATCAGATAA
AGATCTGTTTTAGTAATGCAAAGTTGCGTTTTAAAATGTGAATGGATTTTCAAAAAAATTGT
GAAAGCATATAATTGAAGTTTTAAGAAAAAATTTTGTTTAAATCTTTTAAAACCTTTTTCTTC
TCCTTTTACAGGTCATCCTCATAGAATTTTTACCAAAGACACCGATTTTTCGACCAGATTAG
CATTTACTTTATTTATAGAGACTTTCCAAGTATGTTGTCTTTCCAATGGTGCCTTGCTTGGTG
CTCTCCTGGTGGTGACATAACATTGGTTCTACAGAATCGTGTGGTGTTTTTTTTGTTTTTGT
TTTTTTTTTTTTTAAATAACCGCATGTTCTAAGTGTGCATTTTTGTCAATCTTTGCAACAGTT
ATTTCATACAGATGTTTAATACTTAAGTTATTGTGCTCTTTTCTGTTATGTATTCTGATTTTCA
AGGATTACTTTTTTGTATTATCAAAAAAATACATTTGAACTTAGCATAAAAAGTGGCCAGCC
TTTTTTATTTTGTCACCAAGGTACACACAGTCCTTTATTTATAAATTCCTTAACAGAGAAAAA
CACCTTTGTAAGGCTCAACTTACCTATTCCAGCAAGCACACTTTTTCTGTCATTTTTTCTTT
CTTTTCAAATTTGATATTGTCATTATTTTAAAATAGTAAGTTTTCTTTAATAGTCTTTTGGGAC
CTAACATACCCTTTCTCATACAATTCCTAATGCTCTGTTTATGGCAGATAATCTGTAATGTTA
TGAAGACCTATCAAAAAGTTTTAAAAGTATTTCTGTCTTCAAAGGTAGTAAGACAGGATTA
AATTTTTATTAGAATAGACAAATCAGTGAATGGTATGCATGTATCTAGTGGTTACTAGAACTC
AGGATCACACAATATAGTAGCATCACGATCTGTGTATATTTTTGATCAAGATGATAATAATGG
CCTTACTTGGGTTTTTATCGTTTATCAAATCTTACATACAAAAGAGTGGAAGTATTCCTTTAC
AAAATTTCTAAGGAAAATATTTCTTCCAATCTATCACAATTATAGAATGGATATATGTTTCTG
AAAAGTTTTTGAAAGAAAGCAAAAGTTCTAGAACTAAAAGTAAGCTGGTATTTAATATCCC
GTTGATATTTAGAAAAGATTGTTAATAAGAAATGGAGGATGCATTTAGTACTATTTTTATCCA
CTAGTTCACTTTCAGTACAGTTATGTATACTTGTTTTGATTGAGAGTGTGACATACATGTTAA
ATCAGATTAGCTTGTTTCTTTTAAATATACATATACACAAATACATATAATTTTTTCTCCTTTTT
GTTGTGCATATCTCTATGCATTTTTAAACTTTTAGATTTGTGAATGACCTATGTGTAAATTTTT
GTTTTTATAAACCAGAAATTATACAAGTTTTAATGTGTGTCAAGAACTTGTTCCATACAACT
GTGGTATCGAGCAATAATGTTAATAACTTTTGGAATTATATAAACTATGCTTAATAATTTGTAT
TGAGAATTGGTACCACTATACAATACTTTTTTCCTGTATTAAATCTTTTAAATACCAGTTTCAT
TAATTTATATACCTGTATTTTTAAAAGTATTTCTTTGTACTTGCCAAAGTACTGTAATCCTATAT
AAATTTGAATAGTCAAATGCCAGTATCTCATGTCCTATATCCTGTCTTGGATATTAGGAGCTA

TCTTAAAAACTAGGATTTAAGATTTTTTTTTGCTTGAGAATTATATGTTTTCCTTGCCCCCCA
AGTAACAACTTGAATTTGTAGAACACTTTTCCTGTGAAGAGTTCAAAGTACAAAGAACTCA
AATACTTATTTTATTTAACTTGTATCTAGGGTGTGTCCTACCACCTTAAAGTAGTGCAATTAA
AAATAATGCCATCTAGAAACTTCAGTATTCCTCTGTGGTTTCAGTGTGAAAAATAATTGTAT
CTTCATACAACGCTAAATCTGCCTCAGGATTCAGAAAGATCAGAAAAATATAATGACTGTG
TAAACTATGCAACACAGTGTTCCCTGTATAAAATTGTATAAATCCATAGAGCACCATATTGA
TAAGCTTTATGAGAAGAACTTATCTATTTACATAATTTATTCTCCTGTTATTAGTCTACTGTGG
ATAATAATTTTAAAATTACTATATTCCCTTAGGGATATTTTAACTGTAAGATAATTTTCATAGTT
GATTATATTAAGTATAGAATCTTTAAAAGCTTTTCTGATTTTATTCAGCCATGTGCCATTTGCT
TAAAGATTCCCTAGAAACATACACTTCAATGTATATATAGAAAATAACTGTGAATACTTAATA
TGTGGTATTATAAGCCAGTTATGCCTTGGGCATTTAGATTTTCATGCTTGGATTTTCATTAGT
AACCTGTAGTATAACTGTATCCTTTCTTGAACCATGAATTTAAGTGCCCACATTACTGGATA
GTAGAGATGCCCATATTAAGTAAACATCAGTCTAAGGAAATATTTAAAGTGATGATTTTTCT
TCAGCTTCTGTTTTCTGACTTAAAGTTTGTGGGACATAAACTGTAGAGCTTTTTAAACTGGC
CAAGAAAAAATATTATTGGCAGAAAGTTTGGTATATGGATTCTACATAGGCTAAGGTAAATA
ACTTATGGTCTTTTTAAAATACATTACTGGCATTTTACAATCAGTCACCATTTCTGTATAGAA
ACATTCTTTTCTCAACAAATGTACGTTGACAACCCTGTTATATTTTAATATATAACTCTGGCT
ATCTGGGTTAAAATGATACTCCAAAGCCTAATGACTATAGTAAAACTGATGTGACACATTTA
GCTTCTAAGGTAGATGTTGGCCCTCCCTTACATGGAGATCACCTTACCTATCTAAAATGATC
ATCAGATGCTTTATTAAGACTCTTTAATGACTAGATTTCATTTATTTAAAATATAAACATGTCA
TTTTGCCCTAAATTCAACAAAATAAATTCAACAAAGCCTTGAATAATAGTAAACAATGATTA
TTTTTTCTTATGAGAAAAACTAGTTTTGTAATTGAATAATTTTAAATCAGGAAGAATCCAGCT
TTCCCCAAGGTTTTGTCTTTAGTGTTATTTTTTCAGTGTTAGAGAGCTTCTGGTTGTTTCTTT
ATATTAATAGACGCAGGCCAACCATAAAAATCTTTGACAATATTCTGTCTCTATCTTTGGAT
GTAAAAGGACATTAAATAGTAATACGTACAGTTGGACCCACAGTTGCATGAGTTCAAACTA
CTAGTTTTGTATAACACTGTGTACATAGTCAGACTGTTTCAACTTTTGTGGTATTTAGCTTCA
CAATTATAACCTTTAACTGCATGCTTGATGTTTCACTGTAAATATTTGCTTTGACTTGCTGTT
AATCAGTAGTACTAAATCAAAACTATAGTGTGTTAAAGCTGTCTTGGACTGAACTCCAGCA
TCTAGTCCTGCACCCTCTTTCTTTTTATCCTTTTGGACACCATGGATTCTTTTGCTGTCCTCC
TGATGTCCAGAGTCTTTACCTAGGCCTGAACATAGTTCAGAATTGCATTTTTATCTTTCTTC
CTCAAAGCTCCCAGTAAGAACAGTAATTTTTTTTTTTTTTTTTTGAGACAGTCTCACTCTGT
TGTCCAGGCTGGAGTGCAGTGGCACCATCTTGGCTGACTGCAACCTCCACCTCCTAGGTT
CAAGCAATTCTCTGCCTCAGCCTCCCGAGTAACTGGTATTACAGACGCCCGCCACCACGC
CCGGCTATTTTTTTTGTATTTTAGTAGAGACGAGGTTTCACCGTGTTGCCCAGGCTGGTCT
CGAACTCCTGTGCTCAAGCAATCTGCCCACCTCGGCCTCCCAAAATAAGAACAGTAATTC
TTTTGTATTCTTAGGTAGGTGTTTAGAAGACCAATTAAAGTAAGTCGCATTAAGTGATGAAT
GGCTGACACCCTCAGTTCTGCTAAGGTGGTACTTGCATTTTAAAAATGCATTTTAAAAAGA
GATTTCTTAATTGATGCAGTGGTGCAATTGATGTTTCCTTCTGAAAAGACCCTAATGGTCCA
CGCTTTCTAGGGAGGGAACAATAGGATGGTTGGACATGGGGTTGCTTCTCAAACCTGTTT
TTGAAAGAAGGTGATTAATAATTAGTTTTTGGGAGGAAAGAATAGGCTAATTTTAAACCAA
ACTATTAATTATCCCATAGCCGTTTGTGTTTATTTAACCTTTTCTTAAATAAAGTAACTCTTAA
TTTTTCCTTGATATCATGGCCAACTTTATGTTGTTTTTAGGAATATATCTTGAGATTTGAAGA
ATGTCTTTAAATCTTGAGGTATCTTGTTCTGCACTTCTATGGAGATCTTTAGATAATAATTAA
GAGAAGCCTAAATATTATTGGTATTATTAATCATAATTCTATCTGTTTCTGTAATATGGAGCAT
TATCTCCCCCACCCCAATGGCATATTGAAGTAACAGAGGTTGTTTATAATCTTAATAGGTAG
TACTGAAAACAGAAATTCAGGATGTTGATCCTGATTTGGAACTCATTTCTGGTATTCTACAA
GATCATCCAGCTACTTGGCCTTTTTGGTTTCATTTTCATTTCCCAAGCAGGGAGATACTATG

ACTACCTCATGGTGTTATCTTGTGAAGCAATATTTTTCGATAGTTTATTTTCCTGTTATTAAGT
CCTATATAAAAGTAAGTAGTTAGATGTTTAATAATAATGATTGAATAGGTTTTTCAATTTATTT
ACAAAGTCAGCGCAAACTTGAATTATTGTCAGGTTTTGCATTTAAAGTGGAAGAGATTGGG
TATTCCCTATATATACTTATAATGCTTTGCATGTTTAAAAATCAGGGAAACATTAAATATAGTA
CTTTGAAGAAAGTTTCTAATAAGATTTTTAAAAATATAGTTACGTCTGAGATACTTAAACAAA
ATCATGCCTTTTTTTATATCCCCTCTATTTCATAGACATGGTGCTCTGATATTGAGTTGATTAC
TTTTTTCTCTTTATGAAAACCAGTATATTGCTTTCCTAGCTCTTTATCTTAGAAAGTATTTTGA
TGAAGTGACCATACCCGGTATCCTTAATTTTGGCAAATAAGCTGTATATTGCTCTGTTGTTG
TTTATTTAAAGAAACATAGTGTTAAGCAGCCTTCAAGAGACATAAGTGATGGGTCAGTGTT
TTGTCTACTCTACTTTATTGAGACATAACGTACCCTCAGACTGCAGAGTTTCTATCCTATTGT
GTCTTCTACTGGTTCTTGATGTAGTATAACTGGTTCCATTTCTTTTTCTTGGCATAAATTCTG
TATAGATTTAAAATATTGTAGAAGGAAAGAGTATTTCCTCAAGAAACCTTTTGCTCTAAATTA
AAGAACTCCTCTTCCTTTAGTAGCAGCTATTTAATTAGGTAATTAGAACAATCTCATTACATG
GCCACCAAGTCTTTATAAACAGGAGCTTGTTTAACGCAAGTATCAAAGCACTTTTAAATTC
TGTATTTCTTTTTATGTTTGTTGAGGTTATTAGCATTATGCCCTCTCAGAAGGGTAAAGTATT
GTGAAACTTACCGGTTGTGGCAGTAGAAGCAAGGCAAACATGATTGGAAGGCAAAATGC
TGAAATCTACTGGCTGAGAGAAATAACTGCATTATTATCCTACTCTTTGGTTTACAAGGGTG
AATGTTATGCATTGGTCAGGAATTGCACACAGCTTTTTGCTTTCTTTATACATTCTTTATATAT
TACAATGGAAATTATTTTTAAAGAAAAAGAAAAAATATTATTCTAGATAGAATTTTCCTTTAT
CATCATGTCTTTTGAGAAGATAATGGGAGGCTACCTTGAAACAAATATACTTAGATTTATTTT
TTATAATATAAATGTGACCATTAAAAAAAAAAAAAGGGTTCCAGGAATTTGTTCCTTCATCTT
TTTTTTTTTTTTTTTGCCACTAATGTAATTTATATCCTTAAATGAACATGGAATAGTTTTCATAT
AAGATAATCAGAAATTTCTGATGGTGAGCACTTATGTAGAACTAAGAGCAGGGTCCCTAAA
AATATGCTTTCAGATAATTATACAGCATTCAGGTAAGTCTAGTAACATTAGAAAATGAAGAT
AACTGGTAAAATATAAAGTTGAATGTAAAAGTTATGTATTATGTTTTTACTTTCATTATTAAGA
TATTAATCCCACTGCCAAATAAACACGGTTTTTTTTCTTTTTTTGAGACAGAGTCTCACTCT
GTCGCCCAGGCTGGAGTGCAGTGGCATGATTTTGGCTCACTGCAGCCTCTGCCTCCCAG
GTTCAAGCAATTCTCCTGCCTCAGCCTCGCAAGTGTCTGGGATTACAGGCGTGAGCCACC
AAGCCCAGCTAATTTTTGCATTTTTAGTAGAGAGCATTTTGTCATGTTGGCCAGACTGGTCT
CAAACTCCTGACCTCAGGTGATCCGCCCACCTCAGCCTCCCAAACTGATGGGGTTACAG
GTGTGAGCCACCACACCCTGCCTATTTTTCTTAAAATGAGATTGTTTTGTTAGATAAAAACT
TGTTGGCCTTCATAATTATGAAGAAATACATAATGGGTTATATTTAAGGTGGCTTAGTTTTTG
GTCCTGTGTTGGTTGTAGTATATTTAGTCTTGTAAAATTAAGTCTTCATTGATTGTATTGATTG
TTCTTTTTTTAGTAAACAAATGGAGACTAATTTGAGCTAGGGATTTTTTTTTTTCCCCAGTTT
TGGCTTGCTTTGGCTAGTTGAGTTTTGTTGCTTTTAAGTGAGTCTACTACAACATGTCATTT
CAACACACCATTCCATATTCAAAACTAAAATATGGGCTAATTCACAGCATTTTGTCTTTAAA
GAATATAATATATTAAATAAGATCGTAAGCCAGTGTATTTGGAATTGGAAAAGGGAATCTGG
AACTTAGATCAAATCACTTTAAAGGAAATTGGCCAAACTAATAATGTCTGTTAAAATGACCA
TATGAATGAATGCGTACACGCACACACACCTGCTTCTGTAGATTTGACCTGTACAATGTGT
GGGGAAAAACTAATTAAATTGACGTCATTATGAATATGAGTTTAGAAACTGTGTACAATGAA
TTTAAATAAAGTTTTATGTTATATGTTCATGGACTTACATCATAGTTGGAAAAAAGAATAGGG
GAAAAATGTAGCTTATATAATATAATATTACCTTTTTTTTCCCCACAGTGGTGTTTTAGTTATC
CCTTAAGTTTAAAATTAATGCAGCTTGTAAATAGGATTCTCAATTATCCCGTTTCAGTTTTTA
AACTTTCAGTAAGAGGCAGAAATAATTCCAGTTGCCACTTCTGTAAATAGACGTTTAGCAC
TATTTGAAATAGCTTATCTGTGGCATTGTATCAAGTGACTATTTGTAATACAAATTTTTCATTT
TACTCAATTGACCCAGATTCCAGTCTAAACCTTTACTAATATAGATCCATATATGGAAATACA
CATACACATACACACGCAGAATCTCAGTACACTAAGACATCTCTTTGCAGTAGACAGTAAA

GGTAACTCCATCACCTGGATTGTTTCAGTTAAAATACAAAATGACAGTTGATTCTATATGAA
ACATAAAAACTAAAGGGAAGTTTTGTGAGTTAGTCTTGTTTCCTTGAGTCATCATACTGTAG
TTTTTAAAGAATTGATAACTTATTTTCTGAATTTTTATTATTTTATTCACATTTTTCATCCTGAC
CAGAAAAACGTGAGCATTCTCAGAAGTGACTTGTTTCTTTTTTCACAAAGAGAGACCTATG
CAAAAAATGCCATTGCTGCTTGCACTAGTTACTATCTTAGAATCCATCACAGAAAGCCCAA
AAGATAGCACAACCTTTTGTACCTACAGTCTTAGATATTGTTAGATAGCCATTCAGTTTTGTT
TGCATTGTGTATGAGATGAATATTTCTTGAGACTTTTTTGTTTGCTCTACTAAAGAAATCAAC
CAAGGGAAAAAAAATCCAAAAAACAAAACAAACAAAAAACACCCTCTAACTCATGTGAA
GCATGCAGGGTGTTGTAATTTCAGTTTGCAAAGCGGTGTGATACAATGTTGCTGAGCCAA
AAGCACACGATAGATTTAATGTAGCCAATTGTGTACTTTTGAAAAAAAGAGTAACTGTTCC
CTGTGAGTTAATTTTGGATTTCTTCAAAATCTCTCTTTTAGGCTTTGTGCTGGATTCTTCCAG
ACAAATGCGTATTCCATGTGGGCCACTGTTCTGCTAAATGCTCTCAGTTCTCCTTTTGCAAT
CAAGATTATGTTGCTAAGGAGATTTTGCTTTTATTGCTGCCATTGATTTGTGTTAATACGTTT
TGAATACCTCTCGGTATTCTTTCGTAGACAAGGCCAAAAGAAAAACTTCCCCGAGTTTCCC
AATTTACATTACAAATGGAGCGGTGGTGTAATGAAGCCGATATAAAGTGGGCAGTTGAAA
GGAGACTAAAGAAGGGCACTCAAGTGAGAAATGAAGAAATGTGCCGAGTGCAGTCTGTG
GGCTGCCCTTGGTCCAGCTGCAGGAACTGGCTCTGGCCAGAGGTAGAGTGAGTGAGCAC
CTCCCTCTGCAAAGACCACCCACTTTTGGAGCCTAGGCTGGCACACCAAAGATTACCTCA
TTGATTAATACAGGGCTTGAAAATTTCTGAGGTGGCAAAATCACTTCTAAATCTCTTTTTTG
AGTCCTTCCTAGAGCATGACTGAGGCCACACTTCCTGTGCCTTATGACCTGAAGTTGTAAA
GGATAACCTATGGATGGACCAGTTCTTTTGGTTCTGAGGACTAGTGTATTTCAAATGTTGG
AATGTATTCCTAGGTGAGGAGGCTATTAACTAGATTAAATTGGTTTTAGTTATATGCTTCTTA
CTTTGTGTCATATTTTAGACTCCAAAAACTGTCTTCCTACCAAAAAAACCCAAGTATTCCAT
CTGTGAGTGGAAAGAGACCAACTAGATGTGTCTTTAACCAGCGAAGTGGTTTGCTCAGGT
TGCAGTGCGTGGAGGTGCTAAAGAATAGCATGGGCCGGGCGCAGTGGCTCCTGTAATCC
TTGCACTTTGGGAGGCCGAGGCAGGTGGATCGCAAGGTCAGGAGATTGAGACCATCCTG
GCTAACATGGTGAAACCCCGTCTGTACTAAAAACAAAAAATTAGCCGGGCGTGGTGGC
GGGCGCCTGTAGTCCCAGCTACTTGGGAGGCTGAGGCAGGAGAATGGTGTGAACCTGG
GAGGCGGAGCTTGCGGTGAGTCGAGATCACGCCACTGCACTCCAGCATGGGCAACAGA
GCAAGACTCTGTGTCAAAAAAAAAAAAAAAAAAAAAGAATACCATGGTAGCACAGTCATAC
CCTTTAATGGCTCACATTGTTCAAACAGCATGATATTAACTTTTTTTTAAAATCAAAGCACA
AATGCTTCATGCATATAACTTAAAGAGTTTTAGATATTTTAAGTTAGAAATGCCAAAAATATA
AAAACCATGAAGAAAAATTTCTATAGAAGTTAGGAAATTAAAAATCGATCATAAGGGAAGC
ATCTCAGGTTTTTATTGCAGTATTTGGAGATGAAAAATGACTTTCTATTGACACGATTTGTTA
TTTCAGCAAAATAGAAGTCTGTCAGTCTAATGTTGGATGATCTGATAGTCTTTGCAAGAAAT
AATTGCAGAGAGTTTCTGCAATCACTGGGAAATAGAAATCTATAAAAACATCTGAAAAAGT
GTGAGACTGGGCCACTGCCTAGGGGGATAGCCCTGTTATTTCTATGGAACAGGAAAAAAA
AAAAAAAAACTCTGAAAAGGAAATTCTTAATTCCTGTATTTACAGTGCCTTTTTCAGTAGGT
TTTTCCTGCAAAGGGTGCTTGTCAGTTACTTTGCTTTAATGGATAAAATCTCAATTTCCAGG
CCAAATGTAACTTTTTTTGGTCTTACATCTTGATTTACTTGTACTTAAGAGGTTTTTTCTCCC
TGTAATATTTTTAAAGCTTTTAAAATTAACTTCAGCTTAAAGACAATATAAGTATGAATTTCTG
TTTCCAGCCTTGTAGTTCTGACTGTATTTGTCAATGGTTTCTATATTAACGATAGCATTGTTC
CATTGTGTAATCCTAAAGTTAAAAATTCTAAGAATGAGAAAAACAGCTATGAATAAATTATA
AGTGAAAGATAGAAAAAACACATATTTCTATACTAAATATTGAGAGCTTTGTATTTTTCAAGT
TGGAATCAATATATTGAGAACTGACTTTAAATTTTAAAACTCTTATGCTGAGAAACCCTGAC
AGGTAGATAAACCCGTTTGTCTACATAAGATGTTAGCAGTGTAGTATTGTGCACATCAAGC
ATGTTACAAAAACTCTAAGTTATTCTCTGAAGATACATAAGTTTTACTAGAGATACTCTGGAT

**TTATTAGTCCTGCAAAGCCAGCTTTTTATAGTCTTCTGTTTGTAACATATCTGCTGGATGGTA**
**ATTTCTGCCATGATCATCTGACCAGTGTTTCTTCTATATGCATTTTAGAGCAAAGATGAACT**
**AGGTCATCTATAAACAAATGCATTACATGTTTTAAATCTGTCTATCAAAACTGTTGACAGGT**
**GTGTCAAGAAAGAGCTCTGAAGAAAGCGTGGGGGAGGAGGATGGGAGAGGCATTTTTTG**
**TTAGTTTGTTTCCCAAATTCTGTTAGCATTGGGTCTTGCTTCTAAGCTATAGGCCACATCCA**
**TTTAAAAGAAGGTGGAGTCTCAGAGCTGGAAGAAACCTTGGAAGTATCTAGTGGAGATGT**
**GCATAAAATCAGCTGGGAAGTTTTTTGTTTGTTTGTTTGTTTATTTATTTATGAGAGTCTTGC**
**TCTGTTGCCCAGGTTATAGTGCAGTGGTGTGATCAAGGCTCACTGAGGCCTCCATATCCCA**
**GTCTCAAGCAATTATCCCACCTCAGCCTCCCAAGTAGCGGGGACTACAGATGAGCACACC**
**ATGCCTGGCTAATTTTTAAAGATTTAAGAGACAGAGTCTCACTATGTTGCCCAGGCTGGTC**
**TTGAACTCCTGGGCTCAAGCGATCCTCCTGCCTTGGCCTCTCAAAGTGTTTGGATTTACCT**
**GCGTGAGTCCTTTGCTGGCCCTTTAAACAAATTGTAACATACTATTCAAGTATCTAGGTTTT**
**AACAGGGTCAGCTCCTTTGTTTTATCTTCCTCTTCAGTTAAAACCATTCTCCAGTTGATTTT**
**GAATGATTTAATGAATGTCATTTTGATCTTGATACAAATGCATTCCAAAACTGTAATTTTGAT**
**GAGTAAATTAGAATTTTAATATGGGCTTAATTGTTTTTTTTGCGACGGTGTCTCAGTTTGTCA**
**CCCAGGCTGGAGTGCAGTGGTGTGATCTTGGCTCACTACAACCCCTGCCTCCCGGGTTC**
**AAGCAATTCTCCTGCCTCAGCCTCCCGAGTAACTGGGATTACAGGCGCCTGCTACCACGC**
**CTGGCTAATTTTTTTATTTTTAGTAGAGATGGGGTTTCACCATCTTGCCCAGGCTGGTCTCG**
**AACTCCTGACCTCAGATGATCCACCCACCTCAGCTTCCCAAAGTGCTAGGATCACAGGCG**
**TGAGCCACTGTGCCCAGCCCAGTAGTTTGGTTTTAAAACTACACTTGGCTCTACCCTCCAC**
**GTTAGAATTGGGAGGAAGGAAAGAAGACCATGTATATTTTCAATATTTTTATCAATTATATCT**
**TCCGGCTGGATGTGGTGGCTCACACCTGTAATCTTAGCACTTTGGGAAGCCGAGGTGGGA**
**GGATTACTTGAACCCAGGAGTTCAAGACCAGCCTGGGCAACATAGCAAGAACCTGTCTCT**
**ATTCTTATTTTTAAAATTTTAAAAAAAGATATCTTCCCTTACCTCTCTAACCTAATTTTATAAT**
**GACAATAAGGCTTAGGAAGTTTATCACTTATCTTCCATTGCAGGTTGTTAGTGATTTTTAAA**
**AATCCCAAAAGTGATCATTTAGAATCCTTGTAAGAAAGCATGACCACAACTGGCTCACTGT**
**AACTGAAAGAGCATCGCCTATGCCTGGGGAACTGAAAGGTAAATTGTTGGATTCCAGCCA**
**CTCAAAAATTGTTTATCTGTCTCTGTAAGTGGTTAAGTTTTAGTTCTATCTACTGCCCAATCT**
**GAGAACTGTGACTTAGACATTTAATTATTTATTGCTTGTCCGTGTGAAAACATTTAGAATGG**
**CTGGCAGCCTACCAGACCCCGAGTTAAACATTCTACAGCCTGACTAGACTGTTGGGTAGC**
**CAACTTCACATCAGATTAAACACATTCTCTTCATGATGAATAGCTTGGAACCTAAAAACTTA**
**ATTTCGTTACAAAAAGTGCAAGGGGTGTGTTGTTTTCTAAATTAAAATTATTTTATGTTTTAC**
**TGCCACCGATCCAATTGTGGTTGTCAAATGTATTTTCTTTGGTGCTGGTAGGATGGAAGGG**
**AAAGGGACAATGAGGGACTTTTGGGACTAATAAAATTTAAAAAAGGACTCCTACAAGGTT**
**GGACACAATGCAGCCTAGGGATGAACAGTCATGAATTTTGTACTGAAAAAATTTACATTTC**
**ATTAACATTTTAAATTACAAAGGCAGGTCTACAAATACCGATTTGTGTGTTTGTGTGTGTGT**
**GTGTGCTCATTGCATTTTCCAAGCAAACAAGTGAATAATGTGCTAACTGTATCCTTTGTTTT**
**TTTATTTTTGATTTCTCTTTTTATTGTAAAACGTAAGCTGTGATGTGTAGTGACATTAATTTTA**
**GTAAAACCTACTTTACGTTCTCTATAGATAATATGCACATAAAGTCTACATAATGTTTTCTTAT**
**GTAATAAATGCTTTGGGATAATGAAGGGTACTTCGAGCATCATTCCTCAGCTGTTTAGTAAA**
**CCAGCTCTACAGAGACTTGCCTTTCTAATTCTGCAGCTACATGGGTAGAAATTAATCTACTT**
**GAAAATGAATTAGCAATATAAAAATGTAATATTTTATGATTAATATTAACTAGAAATTAAGACT**
**GAATTATTCAACTTCAGTTTGCCATTCTTTGAAAGAAGTTACATTTTGGGAAGTTCTGTGAA**
**GTTAGAAAAAGTTTGAAAGTGTGTTGCGGGAAATAGTGGCTGACTAGTGTAATGAATTGA**
**AACGTTGGAATCAAGAACTCTGTGCCCAAGGGACCTGGCTGTCCGCAGGCTGTCATTTG**
**AGGGTAAATGACAAGCAGGTGCCTGAGAAATGACAGGAAAGGCCCTACTTTTGGTCGTT**
**GGCATCAGTACTAATAACCTTCTG**GCTTTAGATTCTTACACTCT

A35_P_81bp_CG1.1_chr12:58486056-chr1:203364418

AGTGAAGATA**TCACTAACCATCTCCCTTGGCCGTGGGTGGAAGATCCCCTTGCCCCATGTG GCTCCCAGGTGCACCGTTGCACCACCTTGC**TTGCTAAATA

>B171_850bp_CG1.5_Junction5

ACCAGTTAAG**CAGTCCAGTCCCCTCCAGGGACAAGGAATTCAACATATTTTAATGCAGTCC ATCATATCTATTGACTGCATTTAAAAGTTTCTCTCCATATTTAAGCCGACATATTTCTTCTCAA ACCCTTACCTACAAATCCTAAGAATACCCTTTGGGGTTATGCGTAATGAGAAAGTGAAAGG GGCTAATAGCCACTTGTGTATCTGTTCTCTCTTTATGTGTCATTTTTACACAGCGCAAAGGC AGGTGTACCATGAAGCTAATGAAGCTTCAGAGCCTCCAATCACTGGAAGTTGCGGAGTGT TCTAGATGGAGGGGAACAGATTGTGTTTAGGAAGTGTGTCTGGTAAATTGGCTAAAGAGA TCTCAGAAGAAAGGGACCTGAAACTTCAAATCTCCAGTAATTTGTTGTGATTTCTTTTCTC ATCTTTAGCAAGTATTTACTTTTGTATCAATTTTGTACTGGAAACTTCATACTATTTTCCTTAA AGAGGGCTCCAAAAATTGTATTCAGCCCCAGAAAATCTGGATCCACTCCTGGTTTGACCC ATAGCAGTCCTGTTAAATATATTTTCAGTATTGTATTTATGTCTAGGCATTTACATGAGAAAA ATGAACCCCTGTCTTTTTTAACCTACTGTGGTTTTTATTTCGATTTGCTATGTTCGTTTTTTGT TTGTTTGTTTGAGATAGGGTCTCACTCTGTTGCCCAGGCTGGAGTGCAGTGGTGCAATCA CGGTTCACTATAGTCTTGACCCTCCCAGCTCAAGTGATCCTCCTACCTCAGCCTCCAGAGT AGCTGGGACTACAGGCACATGCCACAATGCCCAGCTAAGTTTTTAATTTTTTGTAGAGA**CA GCAGAAG

>B171_89bp_CG2.1_Junction4

GAGATGTTGTAA**GATTTTCATACTTTCAAGTGTTTTTCATACTTTCAAGTGTTTTTCAATAATA ATAATGACAATAATGATACTTAGAATAATCATGATAC**TAAGTCTGATTTTC

>B171_753bp_CG2.1_Junction5

AACAATGAAATAGA**GCACTCCTGACTTGTGTTACTAGGTCCTGGCAGAAAGCAAATGGCA CACTGCACTTGAGGGGTAATTTGAGGAGAGGTTAAAAAATAAGGGGACAATTTATAAAG GGTCGGCGGAGTACAGGAAAACCACAAGAAATAGTGTACAATCTACTAGGGCCAGTAG CAAATCAATGTTTCCACCCCCAGGCCCGAAAAGTCAAAGGAGGGGAGTGATTACCTGAA TTTTTTGTATTTTTTTTAAGTAGAGACAGGATTTCACCGTGTTGGCCAGGATGGTCTCAAT CTCCTGACCTCGTGATCCGCCCACCTCAGCCTCCCAAAGTGCTGGGATTACAGGCGTGA GCCACCGCGCCCGGCTAATGGGACCCAAGTTTTCATGTACACCTTGTACACATAGCCTG AAGACAGTTTTTTGTAATATTTTAAATAATTCCGTGCATGAAACTAAGTTTTCACTGCAAC CTGTCATATGAGGTCAGGTGTTAAATTTTCTACTTGTGGTGTCATGTGAAAAAGTTTTCAG AGGCTGGGTGTGGTGGGGAACCACAGTAGAATGGTGGCAGGAAAGACTTTCTTTTTCAA ACATAGCCGAGCCTACATGCCTTGCGAATCTTAGAGGCTCAAAGAGAGAAGAGGAGCCT TCATCCCCTGGGTACCACTTCTCCTTCCAAGTCCANGCTCTGGGGCAGGNACCCCCTCG GATGGAAAGGTTCTGATGGAATCTCCCCACAGAGGTCTTGCTGCCTCATCTGA**CACATCC TACTACCGAT
